# NCBP1 stress signaling drives alternative S6K1 splicing inhibiting translation

**DOI:** 10.1101/2024.05.12.593755

**Authors:** Dalu Chang, Mahdi Assari, Chananya Suwathep, Khomkrit Sappakhaw, Chayasith Uttamapinant, Marcus. J. C. Long, Yimon Aye

## Abstract

Subcellular stress profoundly influences protein synthesis. However, both the nature of spatiotemporally restricted chemical cues and local protein responders to these cues remain elusive. Unlocking these mechanisms requires the ability to functionally map in living systems locale-specific stress-responder proteins and interrogate how chemical modification of each responder impacts proteome synthesis. We resolved this problem by integrating precision localized electrophile generation and genetic code expansion tools. Upon examination of four distinct subcellular locales, only nuclear-targeted electrophile stress stalled translation. We discovered that NCBP1—a nuclear-resident protein with multifaceted roles in eukaryotic mRNA-biogenesis—propagated this nuclear stress signal through a single cysteine (C436) from among its 19 conserved cysteines. This NCBP1(C436)-specific modification elicited alternative splicing of >250 genes. Mechanistically, global protein-synthesis stall was choreographed by impaired association between electrophile-modified NCBP1(C436) and SF3A1, an essential component of spliceosome, triggering the production of alternatively-spliced S6-kinase, whose expression was sufficient to dominantly inhibit protein translation.

**Summary:** Nuclear reactive electrophile stress suppresses translation via an unprecedented gain of function mechanism. First NCBP1 is labeled at one of its 19 cysteines. This triggers alternative S6K1 splicing, which is dominant negative for translation.

## Introduction

The canonical protein code contains 20 amino acids, equipping nascent proteins with a versatile array of building blocks. When positioned appropriately, these elements alone can perform awesome feats, including proteolysis, sigmatropic shifts, and redox relays. Combined with cofactors, more possibilities arise, e.g. decarboxylation and proton-coupled electron transfer. Nonetheless, the pressures of selection have frequently forced nature to accessorize its canonical amino acid toolbox. These embellishments occur classically in the form of posttranslational modifications (PTMs) ^1^ of specific residues that change chemical properties, fundamentally altering regulatory modes, and often times, function, of proteins.

Numerous efforts have been expended into decoding PTM-signaling modes at the pathway and protein-specific levels, giving way to the ubiquitin and histone codes, and our understanding of phosphosignaling networks. These signals implicate themselves—often in a synergistic manner—in almost all cellular processes. Such regulatory understanding has, in turn, had important ramifications on drug design and disease etiology. Indeed, these canonical PTMs are enacted by specific proteins, ‘writers’, that code these modification-specific signals. These messages are, in turn, interpreted by specific downstream proteins, ‘readers’. However, over the years, an alternative PTM-driven signaling paradigm has arisen, whereby signaling occurs when electrophilic small-molecule metabolites—without the involvement of specific writer proteins—covalently label specific proteins^2–4^.

A protein’s intrinsic electrophile reactivity and the signaling consequences of electrophile-driven PTMs are still mostly unpredictable. Yet, several pathways have emerged to be responsive to non-enzyme-assisted PTMs orchestrated by innate electrophilic metabolites, oftentimes resulting in dominant loss-of-function or gain-of-function signaling through modulation of a specific protein of interest (POI)’s function/activity. Examples include kinase isoform-specific signaling, transcription, antioxidant response, and mitochondria-associated programmed cell death^5–7^. Nevertheless, despite attempts to decode these non-canonical PTM-signaling events, scant paradigmatic examples remain. Locale-specific examples are particularly rare. We remain unaware of how specific localized reactive chemical signal inputs propagate along specific pathways involving defined mediators, leading to functional biological outputs. Accumulating evidence indicates electrophile-responsive pathways are triggered even at low electrophile modification stoichiometry [‘ligand occupancy (LO)’ hereafter] on a single electrophile-responder protein. We have also disclosed organ-specific electrophile sensitivity of an otherwise multiorgan-resident protein and how this tissue-specific electrophile responsivity regulates global stress management in live nematode worms^8^.

Among the limited instances of biological processes directly influenced by such locale-specific substoichiometric electrophile sensing and signaling, protein translation remains a largely unexplored arena. This is despite the emerging phenomenon surrounding dynamical changes in translation efficiency in response to localized subcellular stress^9, 10^. Here, we used an unbiased proteome-wide screen to identify novel locale-specific electrophilic-stress responders, in tandem with a genetic-code expansion^11^-based translation reporter platform. This approach identified the nuclear protein NCBP1 (CBC80) as a novel sensor of the native electrophilic lipid-derived metabolite, 4-hydroxynonenal (HNE). Despite numerous cysteines within NCBP1 sensing HNE, only HNEylation through one specific site (C436) downregulated translation. Translational stall was choreographed by reduced interaction between HNEylated NCBP1 and an essential splicesome component (SF3A1). This event triggered splicing deregulation of numerous genes. One affected gene was S6K1, which formed a new alternatively-spliced isoform, which we termed S6K1-X. S6K1-X was dominant negative for translation, explaining the majority of HNEylated NCBP1-induced translation inhibition and how NCBP1(C436)-specific HNEylation impaired translation. We thus provide one of the first examples of a specific protein—electrophile labeling event impacting translation. We thus further document an intricate molecular mechanism of translation regulation that proceeds through a gain-of-function splicing event. Interestingly, S6K1-X was selectively upregulated in cell-based Huntington’s disease models that also manifested increased HNEylated proteomes. These data overall display the huge richness of reactive small-molecule signaling, and highlight the unpredictability of the important non-canonical PTM-signaling events.

## RESULTS

### Nuclear electrophile release suppresses protein synthesis

To cast a new lens over how localized small-molecule stressors reprogram translation, we set out to unite genetic code expansion-based translation reporting-(GCER)^11, 12^ with our recently-developed function-guided proximity mapping (Localis-REX)^6^ (**Fig. 1A**). Localis-REX deploys the self-labeling protein, Halo, as a permanent anchor for a photocaged electrophile; in this instance, 4-hydroxynonenal (HNE). The photocaged electrophile, “Ht-PreHNE” hereafter (**Supplementary Fig. 1D**) releases native HNE for experiments studying signaling, or an alkyne-functionalized analog, HNE(alkyne), to enrich HNEylated proteins or validate labeling. The REX-setup thus enables generation of HNE in a specific locale (organ/cell-type, subcellular locale, etc.) in live cells/animals with high spatiotemporal resolution (**Supplementary Fig. 1**). Following Halo expression at a specific locale (**Supplementary Fig. 1A**), treatment with Ht-PreHNE(alkyne), washout of excess unbound probe, and light exposure, liberates HNE(alkyne) (*t* ^photouncaging^ < 1 min) in Halo’s vicinity and hence at significantly-elevated concentrations within this locale. Post-lysis, enrichment of HNE(alkyne)ylated native proteins enables target-ID of local electrophile-responder proteins. Conversely, GCER measures dynamic changes in translation (**Supplementary Fig. 1B**) in response to electrophile build-up in a defined locale. Comparing effects across Localis-REX-assisted HNE build-up in different locales furnishes precision information about HNE’s locale-specific effects on translation.

**Figure 1.**
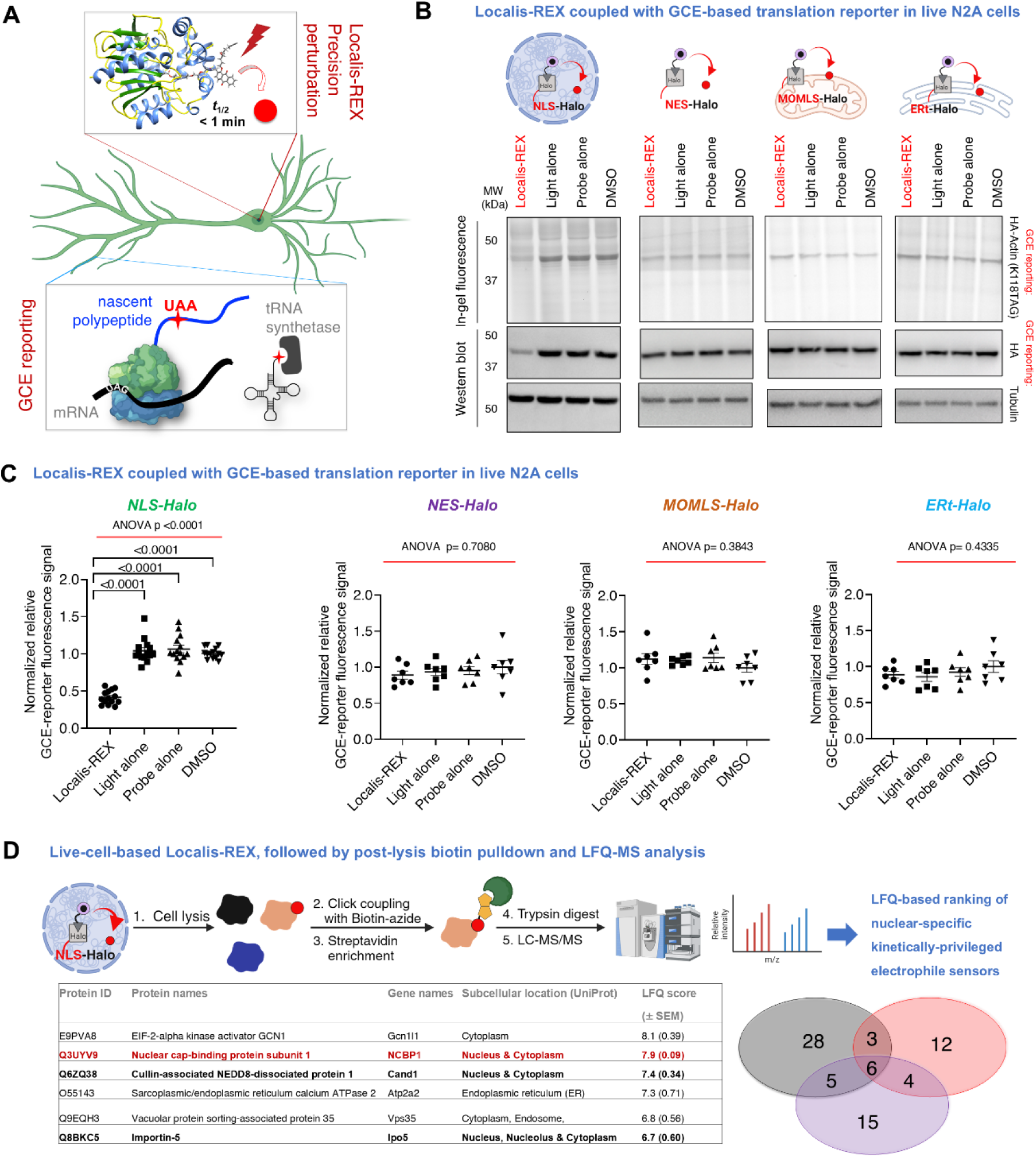
Global protein synthesis is inhibited by nuclear-targeted electrophile delivery. **A)** Schematic of Localis-REX—GCER. *<u>Top box</u>*-(see also **Supplementary Fig. 1A**): the small-molecule electrophilic stressor (red sphere) is liberated rapidly (*t*_1/2_<1 min) upon illumination (5 mW/cm^2^, 365 nm) from the photocaged electrophile covalently bound to locale-restricted Halo. Electrophiles released compete among the best electrophile-sensors within Halo’s proximity, enabling mapping of kinetically-privileged first responders with controled timing, chemotype, dosage, and locale. *<u>Lower box</u>*-genetic code expansion reporting (GCER) ^11, 12^ assesses perturbation to protein synthesis. See also-**Supplementary Fig. 1A,B**. **B)** Differentiated mouse neuroblasts [neuro-2a (N2A) cells] expressing Halo fused to different localization sequences: nuclear localization signal-(NLS); nuclear export signal-(NES); mitochondrial outer membrane localization sequence-(MOMLS); or ER targeting sequence-(ERt), were subjected to Localis-REX—GCER (see also-**Supplementary Table 1,2**, **Supplementary Fig. 1A-D**). In gel Cy5-fluorescence and anti-HA blot report expression levels of HA-actin(K118TAG). Anti-tubulin antibody serves as loading control (n=7). Also see **Extended Data Fig. 1** and **Supplementary Fig. 2**. **C)** Quantification of Figure 1B and **Supplementary Fig. 2**. ‘Normalized relative GCE-reporter signal’ denotes anti-HA signal normalized by anti-tubulin. p values were calculated by Tukey’s multiple comparisons test. Data show mean ±SEM. *n*= 14 independent experiments for NLS-Halo, *n*=7 for rest. **D)** Localis-REX coupled to LFQ-based proteomics (*<u>top row</u>*). The six top-ranked kinetically-privileged nuclear-specific electrophile-senor proteins (*<u>table, bottom left</u>*) from Localis-REX performed in differentiated mouse N2A cells ectopically expressing nuclear-restricted Halo. See also-**Supplementary Table 3**. Venn diagram (*<u>bottom right</u>*) shows the number of statistically-significant enriched hits from each replicate. A stringent threshold was set to limit hits to focus on proteins with highest probability of having phenotypic relevance, see **Extended Data Fig. 3**; lowering threshold could yield more hits. Note: the proteins identified are of mouse origin, but conserved in humans (**Supplementary Fig. 6,7,8**,**10**).

To probe effects in different locales, we fused Halo to four distinct localization sequences, NLS, NES, MOMLS, and ERt (**Supplementary Table 1-2**), directing Halo (hence localised HNE upregulation), respectively, to the nucleus, cytoplasm, mitochondrion outer membrane, or endoplasmic reticulum (ER) (**Supplementry Fig. 1A**). In differentiated mouse neuro 2A cells (‘N2A’ hereafter) (**Supplementary Fig. 1C** and **Methods**), following treatment with Ht-PreHNE and washout, correct localization of probe-bound Halo was observed in all four instances (**Extended Data Fig. 1**). The resultant locale-specific effects on global protein synthesis was assessed in N2A cells ectopically expressing established GCER-components^11, 12^ (**Supplementary Fig. 1B and 1E**).

Initially, we used site-specific incorporation of a bicyclononyne-lysine (BCNK)^12^ into synthetic HA-tagged actin(K118^TAG^), by amber suppression, to assay translation efficiency (**Supplementary Fig. 1B**). Using either in-gel fluorescence analysis, tracking the Cy5-signal arising from actin(K118)-incorporated BCNK (post Diels-Alder-coupling in cell lysate), or western blot measuring anti-HA-signal corresponding to full-length HA-actin (45 kDa), protein synthesis was only prominently affected when electrophile release occurred in the nucleus. No effect was observed when HNE was released in other locales (**Fig. 1B**-**C**, **Supplementary Fig. 2**). An orthogonal whole proteome labeling-based GCER approach^11–13^ validated the same outcome (**Supplementary Fig. 3-4**). Independently, bioorthogonal noncanonical amino acid [azidohomoalanine (AHA)] tagging (BONCAT) ^14^ also gave the same results (**Extended Data Fig. 2**): AHA-incorporation reporting translation efficiency for both the whole proteome and Halo itself were reduced following nuclear HNE build-up, but not other locales (**Extended Data Fig. 2**). Thus, three independent readouts indicated that nuclear HNE stress inhibits translation (**Supplementary Fig. 5**). The fact that release of similar amounts of HNE in three other locales did not cause translation inhibition agrees with our previous data that HNE diffusion between cytosol and nucleus is minimal^6^, and further underscores a nucleus-specific effect. This short diffusion distance of HNE is likely due to cellular detoxifying enzymes that restrict HNE diffusion ^4, 15^.

### NCBP1 electrophile signaling stalls translation

We proposed that (a) specific nuclear protein(s) is(are) responsible for propagating this nucleus-specific signal to drive translation downregulation. Localis-REX function-guided proximity mapping generating HNE(alkyne) (**Supplementary Fig. 1A**)—coupled with enrichment proteomics—can quantitively index endogenous locale-specific HNE-responder proteins^6^. In nuclear-restricted Halo-expressing N2A cells, post Localis-REX-assisted HNE(alkyne) upregulation and lysis (**Supplementary Fig. 1D**), HNE(alkyne)ylated proteins were biotin-modified via Cu(I)-catalyzed Click coupling with biotin azide, and enriched using streptavidin. Following elution, HNE(alkyne)-modified nuclear first-responder proteins were identified using label-free quantification (LFQ)-based proteomics target-ID (**Fig. 1D**). We note this workflow is different from some of our previous published protocols ^5–7, 16–20^ in which we compared responsivity of nuclear and cytosolic proteins^6^. This difference—and the fact that the cell line used in this instance was different from our previous reports—likely explains why our previously-reported hits were not discovered. Localis-REX function-guided mapping revealed 6 protein hits (**Fig. 1D**, **Extended Data Fig. 3**, **Supplementary Table 3**). We note that previous global mapping of HNE-responsive proteins using state-of-the-art iso-TOP-ABPP uncovered a similar number of hits, albeit from lysates in non-organelle-specific contexts^21^.

Among these 6 proteins, nuclear cap-binding protein subunit 1 (NCBP1), CAND1, and IPO5 feature the nucleus as one of their resident subcellular components (**Fig. 1D**) based on UniProt annotations, but the nucleus is not an annotated locale for Gcn1l1, Atp2a2, and Vps35. Nonetheless, Atp2a2 and Vps35 are ER and endosome-associated. Noting also that the ER is in continuity with the nucleus^22^ and endosomes can associate with the nuclear envelope^23^, these proteins may be ‘minority’ privileged electrophile responders at locales distinct from canonical environs—a phenomenon recently unmasked by Localis-REX^6, 24, 25^. We chose to focus on NCBP1, CAND1, and IPO5 where UniProt database clearly indicates nuclear pools. Since human and mouse POIs were highly homologous for all three proteins (**Supplementary Fig. 6-8**), we focused on human POIs.

We used T-REX^16, 18^ to validate electrophile sensitivity. The key difference between Localis-REX (**Fig. 1A-B**) and T-REX is that Halo is fused to a specific POI in T-REX (**Fig. 2A** vs. **Fig. 1A-B**). As liberated HNE occurs in the vicinity of the POI, T-REX assesses the POI’s electrophile sensitivity. Our previous studies show that parameters derived from T-REX are independent of specific POI expression levels^16, 18^: a POI achieving a higher percentage (%) ligand occupancy (LO) (i.e., modification stoichiometry) under T-REX is more sensitive to a specific electrophile than another POI with a lower % LO. Following T-REX in live cells, % LO can be assessed by Click assay in lysates followed by in-gel fluorescence analysis (**Fig. 2A**). Alternatively, a more sensitive readout is achieved via first enriching the electrophile-modified POI via Click-biotin pulldown, followed by western blot (**Extended Data Fig. 4B-C**).

**Figure 2.**
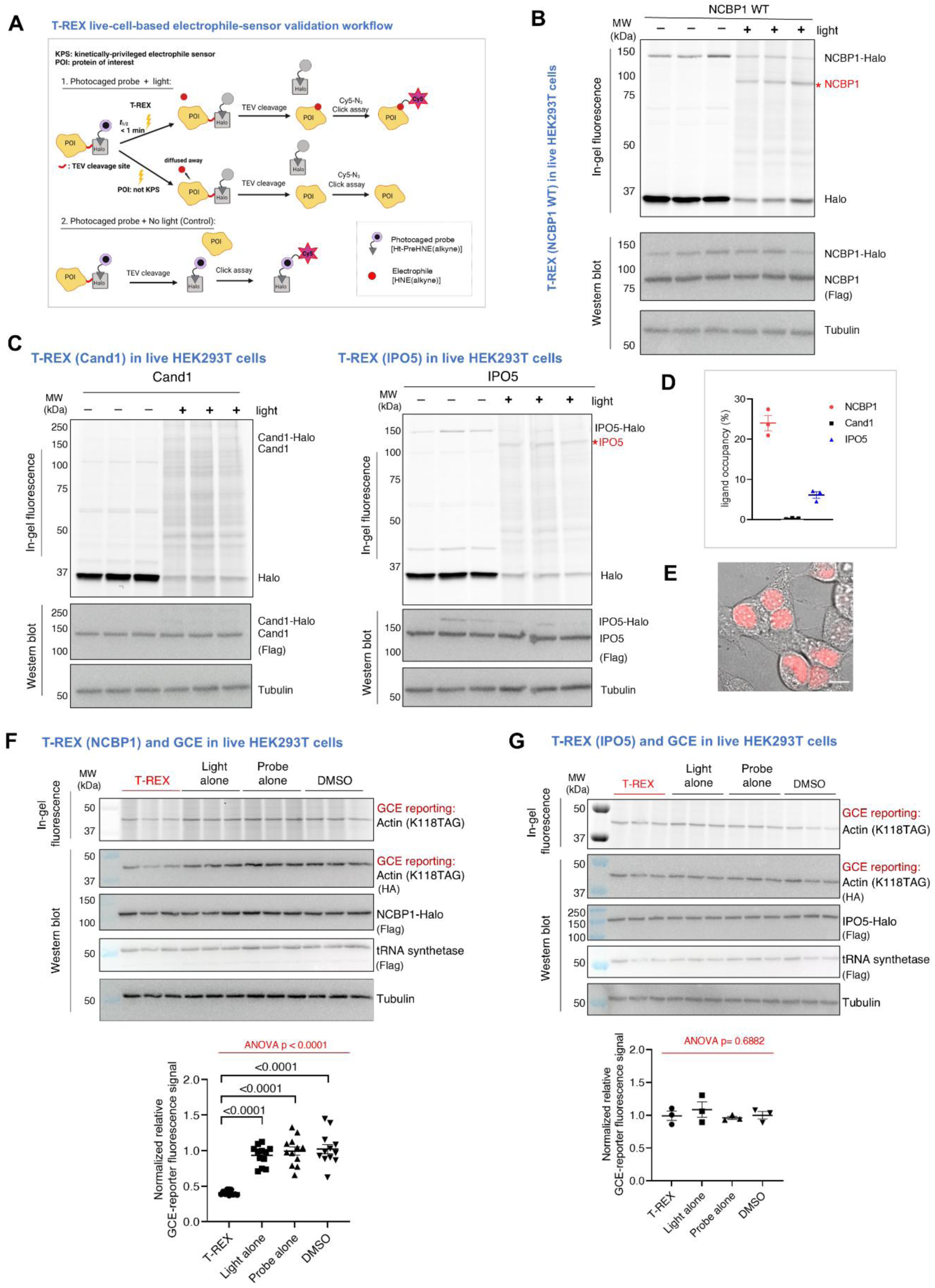
NCBP1 is a kinetically-privileged electrophile sensor, whose HNEylation downregulates translation. **A)** T-REX^16, 17^ workflow: cells expressing Halo-fused protein of interest (POI) are treated with photocaged-(alkyne-functionalized)HNE. Post washout, light triggers HNE-photouncaging (*t*_1/2_<1 min). Providing POI is a HNE(alkyne) kinetically-privileged sensor (KPS), it is HNEylated, prior to HNE diffusion. Unreacted HNE is captured by glutathione (5–10 mM), or enzymatically-degraded^4, 15^, but also elevates background by labeling other proteins. HNEylation can be quantified by in-gel fluorescence post cell lysis and Cy5-azide-Click coupling (**Methods** and **Supplementary Fig. 1D**). **Extended Data Fig. 4B,C** show a higher sensitivity readout. **B)** T-REX in HEK293T expressing NCBP1-Flag-TEV-Halo (‘NCBP1-Halo’) (**Fig. 2A**) validated NCBP1 is a KPS. Samples not exposed to light but treated, post-lysis, with TeV-protease that separates NCBP1 and Halo, are negative controls and define maximum HNE signal. *Indicates NCBP1 post TeV-cleavage (92 kDa). **C)** Similar to **Fig. 2B**. on IPO5 (124 kDa, post TeV-cleavage) and Cand1 (136 kDa, post TeV-cleavage). **D)** Quantification of HNE-ligand occupancy (LO) following T-REX on the indicated POIs, from **Fig. 2B-C**^18^. LO=(HNE(alkyne)-signal on POI band, post TeV-cleavage, under T-REX conditions)/(Ht-PreHNE(alkyne)-signal on Halo without light exposure). Data present mean±SEM (n=3). **E)** NCBP1-Flag-TeV-Halo localized to the nucleus in HEK293T treated with Halo-targetable TMR fluorescent-dye. Scale bar: 10 µm. n=3. See **Extended Data Fig. 4E,6A**. **F)** HEK293T were co-transfected with NCBP1-Flag-TEV-Halo, HA-Actin(K118TAG), and tRNA-synthease, MmPylRS (**Methods** and **Supplemental Fig. S1E**). T-REX was performed followed by BCNK incubation (50 µM, 4 h). Cell lysates were incubated with Tetrazine-Cy5 (1 µM, 30 min). In-gel Cy5-fluorescence and anti-HA blot showed translation suppression following NCBP1-HNEylation, but not IPO5-HNEylation. p values were calculated with Tukey’s multiple comparisons test. Data present mean±SEM (n=12). See **Supplementary Fig. 9A** for additional replicates. **G)** As in **Fig. 2F** except IPO5-Flag-TeV-Halo was used. See **Extended Data Fig. 4C**

All 3 Halo-fused POIs exhibited similar expression levels (**Extended Data Fig. 4D**). T-REX in live HEK293T cells revealed that among the three candidate-POIs, only NCBP1 and IPO5 were kinetically-privileged^15^ electrophile responders (**Fig. 2B-C**). Cand1, which houses 30 cysteines (**Supplmentary Fig. 8**) was not electrophile-sensitive. Notably, % LO(NCBP1) (**Fig. 2D**) was on par with the best electrophile-responder proteins we have examined^5–7, 15–20, 26–29^. Live imaging validated that NCBP1-Halo was chiefly nuclear-restricted (**Fig. 2E**), consistent with NCBP1 housing an intrinsic nuclear-localization signal. The % LO(IPO5) was more similar to weakly-reactive proteins such as HuR^16, 29, 30^. Such proteins have not always shown phenotypic outputs when HNEylated under T-REX. IPO5 localized primarily to the nuclear periphery (**Extended Data Fig. 4E**). On the other hand, the non-electrophile-sensitive Cand1 largely resided in cytoplasmic fraction (**Extended Data Fig. 4E**), providing interesting spatial contexts to protein hits from Localis-REX electrophile responder mapping. We also deployed the alternative biotin-pulldown post T-REX, yielding a more sensitive readout: a similar trend of % LO across NCBP1, CAND1, and IPO5 was observed (**Extended Data Fig. 4A-C**).

To evaluate the extent to which the two HNE-sensitive POIs regulate translation, we coupled T-REX to GCER (**Supplementary Fig. 1E**). This workflow unveiled that HNEylation of NCBP1 suppressed translation, whereas IPO5-HNEylation did not (**Fig. 2F-G** and **Supplementary Fig. 9A** and **9B**). Translation inhibition induced by NCBP1-HNEylation was verified using BONCAT (**Extended Data Fig. 5A-B**). These data indicated that NCBP1’s HNEylation was responsible for translation inhibition during nuclear Localis-REX.

### Of NCBP1’s 19 cysteines only C436-labeling halts translation

Although many amino acid side chains react with HNE, cysteine is the most intrinsically-reactive coded amino acid^4, 31^. Cysteine is most often identified as a kinetically-privileged residue^15^. As an essential large subunit of the nuclear cap-binding complex (CBC), NCBP1 is highly conserved from plants to man (**Supplementary Fig. 10A**). Human and mouse NCBP1 have 19 conserved cysteines (**Supplementary Fig. 6 and 10A-C**)^32^, several of which are solvent accessible^33^ (**Supplementary Fig. 10B**). Each cysteine was individually mutated to alanine. % LO(NCBP1) did not change across all single (Cys-to-Ala)-mutants, relative to wild-type (wt) (**Fig. 3A**, **Supplementary Fig. 11A**). Mutation of all 19-Cys’s to Ala ablated NCBP1 electrophile-sensing (**Fig. 3B**, **Supplementary Fig. 11B**). All mutants—including the ‘all-Cys-to-Ala’ mutant—were soluble and expressed to similar levels as Halo-fusions (**Supplementary Fig. 11A-C**). Although many proteins we and others have investigated have specific electrophile-sensing cysteines^2–4, 31, 34^, several well-known electrophile-responder proteins—including Keap1, the prototypical electrophile-sensor protein^35^—harbor multiple cysteines capable of electrophile sensing.

**Figure 3.**
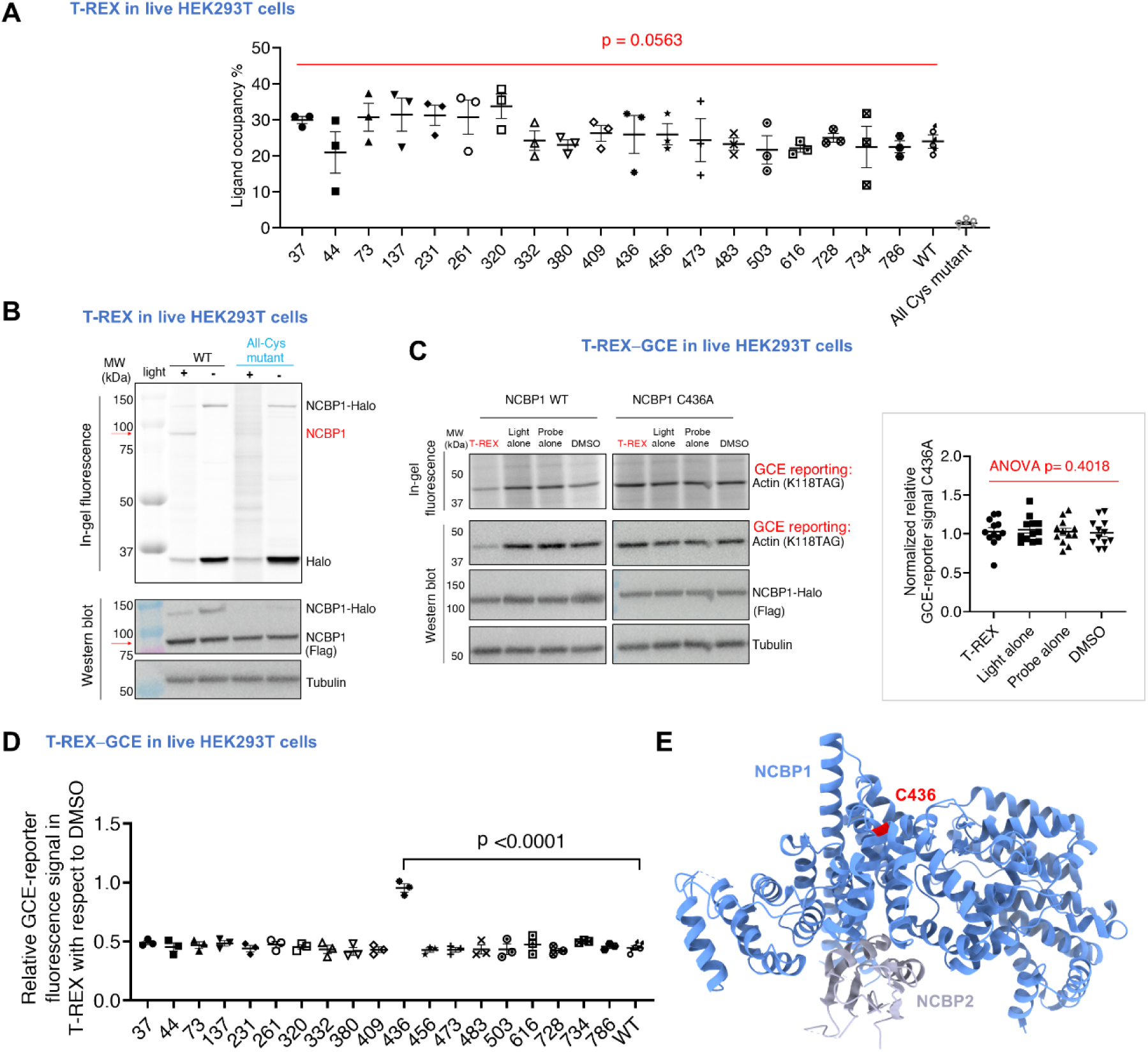
No single NCBP1-cysteine is responsible for sensing; only C436-specific HNEylation suppresses translation. **A)** HEK293T expressing NCBP1-Flag-TEV-Halo (wt, or Cys-to-Ala single-mutant), were subjected to T-REX (**Fig. 2A**) and HNE-LO was quantified as reported^16–18^. See **Supplementary Fig. 11A** for corresponding in-gel Cy5-fluorescence and anti-Flag and anti-tubulin western blots. p values were calculated with Tukey’s multiple comparisons test. Data present mean±SEM (n=3). **B)** Similar experiment to **Fig. 3A** was performed except that comparison between wt and all-Cys mutant (where all 19-NCBP1-Cys’s are mutated to Ala) was made. The bands corresponding to HNEylated-NCBP1 post TeV-cleavage (top fluorescence) and the corresponding protein (bottom western blot), were indicated with a red arrow. See **Extended Data Fig. 6C**. **C)** T-REX—GCER to assess the effect of NCBP1(wt or C436A)-targeted HNEylation on translation. *<u>Inset</u>*-quantification of the results on NCBP1(C436A) against all three T-REX technical negative controls. See relevant validations in **Supplementary Fig. 9,14** that ruled out additional potential interference from technical and biological components. *Inset*: the corresponding quantification for NCPB1(C436A). *p* values were calculated with Tukey’s multiple comparisons test. Data present mean±SEM (n=12). **D)** T-REX—GCER for all 18 single-mutants compared with wt. Results were quantified in each case for the GCE signal (i.e., translation efficiency) measured under T-REX divided by the DMSO control set. (See **Fig. 2F** and **3C**; **Supplementary Fig. 12,13,14** for corresponding datasets from which quantification was derived). p values were calculated with Tukey’s multiple comparisons test. Data present mean±SEM (n=3). **E)** Heterotrimeric complex of human NCBP1 and NCBP2 (PDB-1H6K) with C436 in red. (the 3^rd^ molecule of NCBP2 is hidden behind the pale magenta NCBP1 molecule). See **Supplementary Fig. 16**.

Having validated that NCBP1 senses electrophiles through several cysteines, we tested the hypothesis that NCBP1-HNEylation is responsible for translation suprression when HNE is released in the nucleus. We first ascertained that NCBP1-HNEylation did not affect mRNA-levels of the actin GCE-reporter construct^11, 12^ (**Extended Data Fig. 5C-E**). GCER post-T-REX-assisted NCBP1-HNEylation in cells showed largely similar levels of translation suppression to what we found upon Localis-REX with NLS-Halo (compare **Fig. 1C**, first plot, with **Fig. 2F**, inset). We repeated this experiment with each of the NCBP1 Cys-to-Ala single-point mutants (**Supplementary Fig. 12-13**). Intriguingly, NCBP1(C436A)-Halo HNEylation was unable to initiate translation suppression. However, translation suppression was observed in NCBP1(wt)-Halo, and all other single-Cys mutants (**Fig. 3C-E**, **Supplementary Fig. 14**). Similar results were obtained using BONCAT (**Extended Data Fig. 5F**). We further confirmed that in cells expressing either NCBP1(wt)-Halo or NCBP1(C436A)-Halo, basal translation efficiency remained the same (**Supplementary Fig. 15**). Thus, although several NCBP1-cysteines sensed electrophiles, only C436 was a functional first-responder residue regulating translation. We have reported similar situations for other electrophile-responsive proteins, such as CDK9^6^ and HSPB7^27^. Imaging data also showed that both the all-Cys-to-Ala mutant and C436 mutants were largely nuclear localised (**Extended Data Fig. 6A**).

To further reinforce our hypothesis that NCBP1-electrophile sensing in the nucleus is a key regulatory axis for protein-synthesis downregulation, we replicated Localis-REX in HEK293T cells co-expressing NLS-Halo and non-Halo-fused-NCBP1(C436A), and assayed for translational efficiency (**Extended Data Fig. 7**). We compared in parallel the extent of translation efficiency suppression following Localis-REX nuclear-targeted HNE generation in cells co-transfected with NLS-Halo and either non-Halo-fused-NCBP1(wt) or empty vector [that balanced the amount of plasmid encoding non-Halo-fused-NCBP1(wt) or -(C436A)]. Overexpression of NCBP1(C436A), strongly—albeit not fully—blocked the translation suppression observed in response to nucleus-specific HNE-buildup in empty-vector control or non-Halo-fused-NCBP1(wt)-coexpressing samples under otherwise identical conditions (**Extended Data Fig. 6B,C**). This outcome is consistent with the data above, which showed that NCBP1 is the main HNE-sensor in the nucleus, and that its HNEylation at C436 dampened translation. These results further indicated that other sensors, such as IPO5 (**Fig. 2C,D**), also exist.

All these data point to C436 as a bona fide HNE-sensing residue that functionally couples electrophile sensing to translation. To unambiguously test this hypothesis, we engineered NCBP1-Halo housing C436 as its only cysteine [termed ‘NCBP1(C436-only)’ hereafter]. NCBP1(C436-only) expression and solubility remained similar to wt and All-Cys-to-Ala mutant (**Extended Data Fig. 6B**). Following T-REX, % LO(NCBP1) also remained similar between NCBP1(C436-only) and NCBP1-wt (**Extended Data Fig. 6C**). T-REX coupled GCER demonstrated that NCBP1(C436-only) HNEylation induced the same extent of translation suppression as NCBP1(wt)-Halo HNEylation (**Extended Data Fig. 6E,F**). Post T-REX, we identified a C436-HNEylated peptide within NCBP1(C436-only)-Halo enriched from live HEK293T cells (**Extended Data Fig. 8**), following enrichment digest mass-spectrometry^5, 6^. Altogether these experiments strongly support a model in which translation inhibition caused by nuclear HNE-buildup is ascribable to NCBP1(C436)-HNEylation.

### Translation suppression was independent of NCBP2

NCBP1 partners with NCBP2 to form the CBC that performs numerous context-specific operations in eukaryotic mRNA-biogenesis, mRNA-homeostasis surveillance, and mRNA-translation initiation^36^. The available structure of NCBP1–NCBP2 heterodimer, with and without a bound 5′ 7-methylguanosine (m^7^G)-cap mimic^33, 37^, shows that C436 is solvent exposed and distal to the heterodimer interface (**Fig. 3E**, **Supplementary Figure 16A,B**). T-REX on NCBP2-Halo validated that NCBP2 is not an HNE-sensor (**Supplementary Fig. 16C**). Moreover, NCBP1-HNEylation suppressed translation as efficiently when NCBP2 was overexpressed, as in empty-vector-transfected cells (**Supplementary Fig. 16D**). NCBP1/NCBP2-association was also unperturbed by NCBP1-specific HNEylation (**Supplementary Fig. 17**).

### Survival and homeostasis-related expression were affected

NCBP1 is essential for CBC function^36, 38–42^. Given the CBC’s crucial roles in RNA maturation, we next investigated the effects of NCBP1-HNEylation on RNA levels using RNA-Seq at three different time-points (3, 6, and 12 h) post T-REX-directed NCBP1-specific HNEylation. We compared these changes against all three technical T-REX controls (light alone, REX-probe alone, and DMSO-vehicle) (**Extended Data Fig. 9A**, **Supplementary Table 4**). Pairwise differential analyses were applied to cells that had undergone NCBP1-specific HNEylation relative to each control (**Extended Data Fig. 9B,C,D,E and Supplementary Fig. 18,19**). Each set was performed in independent biological quadruplicates. The numbers of significantly differentially-expressed (SDE) genes—either up- or downregulated—resulting from each comparative set are illustrated in Venn diagrams, along with their identities and thresholds (*p*_adj_) implemented (**Supplementary Fig. 20**).

STRING analysis^43^ of up- and down-regulated SDEs at each time-point (**Supplementary Fig. 20**) showed that several genes lie outside of known interactions. The outputs from early time-points post NCBP1-specific HNEylation particularly stood out in this regard. Subjecting the same SDE-gene list to *g:Profiler* analysis^44^ showed significant enrichment of defined gene ontology (GO)-subterms under biological processes (BPs), and to some extent, molecular functions (MFs) (**Supplementary Fig. 21**) in all time-points. A closer assessment revealed that ~62% (53 out of 86 total) of these top-ranked SDEs are associated with the following (functions): **(i)** transcription factors/coregulators (e.g., *dach1*, *pou3f2*, xbp1, and 10 others); **(ii)** stress/cell-death response (e.g., *txnip*, *hrk*, *bnip3(l)*, *sesn3*, *egln3*, and 15 others); **(iii)** mRNA-biogenesis/translation (e.g., *snrpd3*, *qki*, *tincr*, *mars1*, *yars1*); and **(iv)** receptor–nucleus signal transduction (e.g., *bmp4*, *map2k6*, *cxcr4*, *igfbp5*, and 10 others) (**Supplementary Table 5**). These outputs demonstrate that electrophile modification of NCBP1 has broad-spectrum ramifications on cell homeostasis and survival.

### NCBP1(C436)-signaling modulated splicing of several genes

We were further intrigued by the fact that a significant number of our top-ranked SDEs (either up- and downregulated SDE-genes) (**Supplementary Table 5**) comprised disease-relevant genes that undergo alternative splicing (AS), yielding protein variants with potentially different functions. We first investigated one such gene, *dach1*, by qRT-PCR (**Supplementary Table 6**). *Dach1* is an essential transcription factor in developmental physiology and disease, and was downregulated upon NCBP1-HNEylation (**Fig. 4A**, **Extended Data Fig. 9E**). Following T-REX in HEK293T cells ectopically expressing NCBP1-Halo, and consistent with our RNA-Seq results above (**Fig. 4A**, **Extended Data Fig. 9B,E**), qRT-PCR data showed selective depletion of endogenous *dach1* mRNA-transcript, against all controls (**Fig. 4B**, **Supplementary Fig. 22, Supplementary Table 6**). By contrast, the corresponding abundance of pre-mRNA (assayed using qRT-PCR primers spanning the intronic region flanked by exon-2 and -3), increased upon NCBP1-HNEylation (**Fig. 4C**, **Supplementary Table 6**). These observations were not due to changes in mature *dach1* mRNA stability. As expected, the half-life of the corresponding pre-mRNA was short (~2-12 min, preventing precise calculation); yet this also seemed relatively unperturbed (**Supplementary Fig. 23**). These data indicate that *dach1* downregulation induced by NCBP1-HNEylation was likely due to inhibition of splicing.

**Figure 4.**
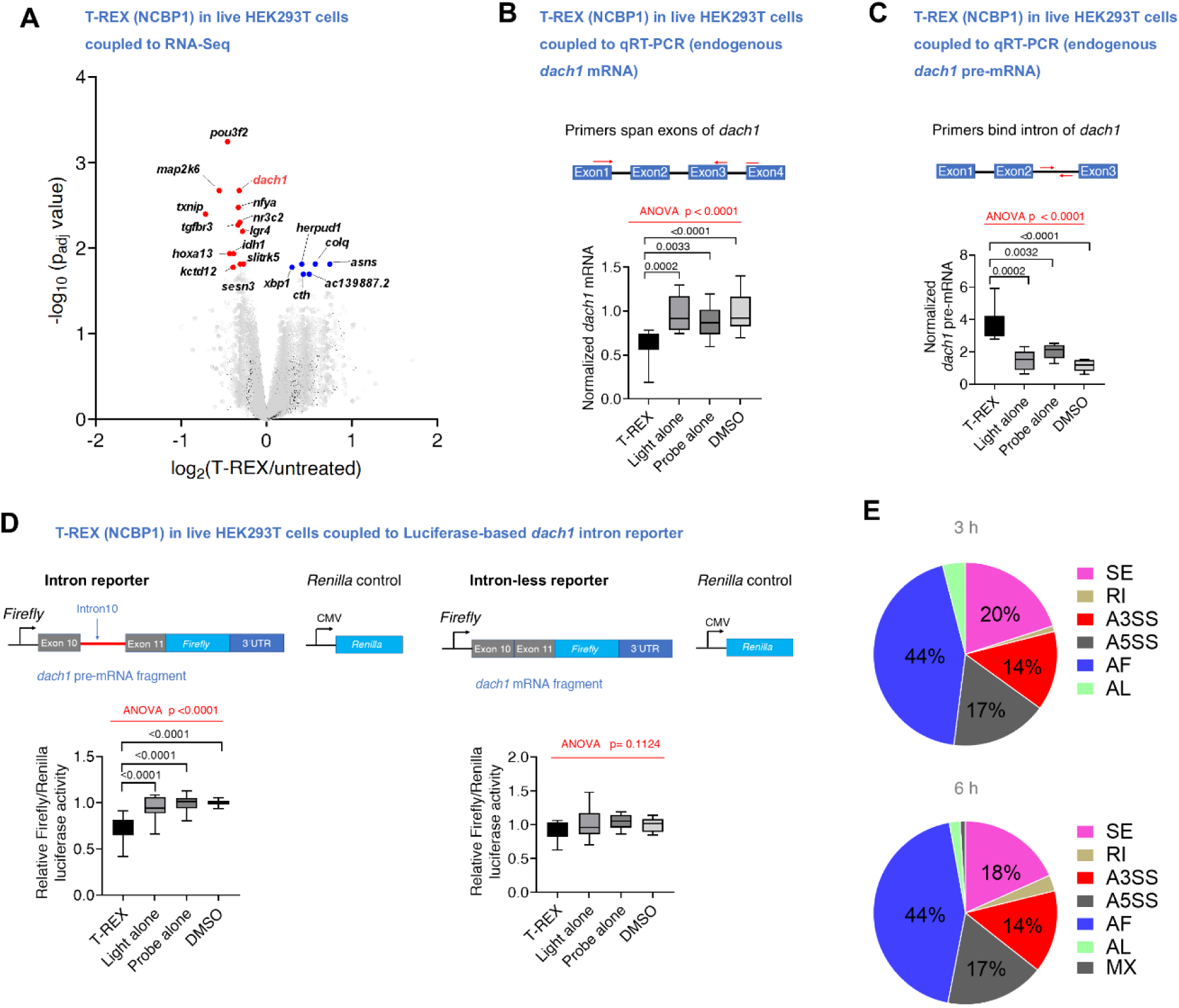
HNEylated NCBP1 modulates splicing. **A)** Differental transcript expression from HEK293T subjected to NCPB1-HNEylation by T-REX against non-treated cells. (3 h post T-REX shown; see **Extended Data Fig. 9** and **Supplementary Fig. 18,19** for workflow and additional time-points and controls). Statistically-significant differentially-expressed (SDE) genes are marked in **blue** (upregulated) and **red** (downregulated), respectively; non-SDE genes are shown in **grey** (see **Supplementary Table 4,5**). For the corresponding MA-scatter plot featuring fold change vs. mean expression, see **Extended Data Fig. 9E**. **B)** Exon-targeted qRT-PCR in HEK293T validated suppression of DACH1 mRNA following T-REX-assisted NCBP1-HNEylation, against negative controls. p values were calculated with Tukey’s multiple comparison test (n=12). **C)** Intron-targeted qRT-PCR in HEK293T revealed upregulation of DACH1 pre-mRNA following T-REX-assisted NCBP1-HNEylation, against negative controls. p values were calculated with Tukey’s multiple comparison test (n=12). **D)** Validation by orthogonal reporter assays. Reporters to assay splicing were built by installing the indicated exons from *dach*1, with/without the intron (left and right, respectively), upstream of a firefly luciferase reporter, with *dach*1’s 3′-UTR downstream. HEK293T expressing NCBP1(wt)-Halo, CMV-Renilla luciferase (internal control) and intron or intron-less reporter, were subjected to T-REX against indicated controls and Firefly and Renilla luminescence were measured (**Methods**). Replacing NCBP1(C436A)-Halo with NCBP1(wt)-Halo showed no change of normalized luciferase signal (see **Supplementary Fig. 24**) (n=18). Box and whisker plots: line-median; box-25-75 percentiles; whiskers-1-99 percentiles. **E)** Alternative splicing events affected by NCBP1-HNEylation. 1266 genes were analyzed across 3 time-points post T-REX-assisted NCBP1-HNEylation, against negative controls, with thresholds for significance defined in the Main Text and **Methods**. Data from 3 and 6 h time-points are shown. See **Supplementary Table 7**,**8**, SE-exon skipping; RI-retained intron; A5(3)SS-alternative 5’/(3’)-splice sites; AF-alternative first exon/alternative promoter usage; AL-alternative last exon splicing; MX-mutually exclusive exon splicing events.

To independently substantiate the above findings, we built ratiometric-readout-based *dach1*-targeted dual-luciferase reporter systems^29^, targeted at an intron of amenable size (intron 10, ~3 kb), flanked by exon-10 and -11. Thus, either an exon-exon sequence (for intron-less reporter), or exon-intron-exon sequence (for intron-10 reporter), from *dach1*-mRNA, was fused upstream of the *firefly* luciferase (ff-Luc) reporter construct, while the *dach1*-3’-UTR was maintained downstream of ff-Luc. Co-expressing this construct alongside a constitutive *Renilla* luciferase (R-Luc) reporter construct in HEK293T cells, we found that NCBP1-HNEylation selectively depleted the intron-10 reporter signal (normalized to R-Luc internal control), but had no effect on the intron-less reporter signal (**Fig. 4D**). These data further pinpoint NCBP1-HNEylation-driven splicing inhibition at intron-10. Importantly, when we deployed NCBP1(C436A)-Halo in place of the wt-protein, no change in the intron reporter signal was observed following T-REX (**Supplementary Fig. 24**).

Beyond its essential transcriptional roles, DACH1 has also been implicated in downregulating cytoplasmic translational programs^45^. We generated *crispr-cas9*-assisted *dach1*-silenced (KO) lines (>90% silencing efficiency in both KO lines, **Supplementary Fig. 25A**). Likely linked to DACH1’s essential role in cell growth, DACH1-KO lines grew significantly slower than the KO-control line, or native HEK293T (**Supplementary Fig. 25B**). T-REX–GCER in these lines showed the same extent of translation inhibition across KO-1, KO-2, KO-control, and native cells (**Supplementary Fig. 25C,D,E**). These data ruled out DACH1 as a player in NCBP1-HNEylation-induced protein-synthesis stall.

### NCBP1-HNEylation promoted alternative splicing

Although changes in *dach1* splicing emerged not to be involved in translation downregulation upon NCBP1-HNEylation, our investigations above indicated that NCBP1-HNEylation could affect splicing. The CBC indeed binds pre-mRNA-transcripts, post co-transcriptional 5’ m^7^G-capping^38, 39^. It plays several roles in nuclear RNA processing, including regulation of constitutive splicing obligatory for mRNA-maturation^36, 38, 39^. Structural studies (**Supplementary Fig. 16B**) hint that NCBP1’s association with NCBP2 is essential for enhancing the latter’s affinity to 5’-capped transcripts^33^. The CBC promotes the initial interaction between spliceosome factors and the 5’-end of the pre-mRNA transcript (at the first intron), augmenting spliceosome assembly and splicing^33, 38, 39^. To examine if the newly-discovered NCBP1-electrophile signaling affects alternative splicing events, we launched a comprehensive genome-wide differential splicing analysis of our RNA-Seq datasets. Following the established SUPPA2 pipeline^46^ (**Methods**), we assigned relative abundances of the splicing events or transcript isoforms [referred to as proportion spliced-in (PSI)^47^] for each transcript across all 3 time-points, under 4 different conditions: T-REX (NCBP1-HNEylation), and 3 controls (DMSO-, light-, or probe-alone treatment). The difference in the mean PSI (dPSI) between the two chosen conditions (T-REX vs. designated control) was subsequently derived for each transcript, for each type of alternative events scored, along with associated *p*_adj_. Events were grouped accordingly: skipping of exon (SE, including both exon inclusion/exclusion; the most common event in mammals^48^); retention of intron (RI); alternative 5’ and 3’ splice-site selection (A5SS; A3SS)’; alternative first exon (i.e., alternative promoter usage) (AF), and alternative last exon splicing (AL); and mutually exclusive exons (MX). Hits that fulfilled the following requirements: *p_adj_* ≤ 0.05 and appeared under at least 2 pairwise comparisons (i.e., T-REX vs. two different controls), out of the 3, were designated as ¢significant events¢ at each time-point (**Supplementary Table 7**). As summarized in **Supplementary Table 8**, changes in alternative splicing were not correlated with changes in its mRNA expression.

Using the above-mentioned thresholds for significance, our differential splicing analysis showed AF (44%), followed by SE (20%), A5SS (17%), and A3SS (14%), were the most-preferred alternative events triggered at 3 h post NCBP1-HNEylation (**Fig. 4E**). Interestingly, in a report where CBC influences AS in *Arabidopsis thaliana*^49^ (~29% sequence identity to mouse and human NCBP1), *Arabidopsis* CBC affects 101 AS events (*p* ≤ 0.1) from the 252 analyzed, with >50% of the AS events being AF^49^, likely consistent with the direct positioning of CBC at 5’ cap and the first splice site of the pre-mRNA transcript^38, 39^ and indeed broadly consistent with our own results. In our survey, 3 h post NCBP1-HNEylation, 200 splicing events, across 119 distinct transcripts, were significantly affected among the 1266 genes analyzed, where transcript counts were determined using Salmon^50^. At 6 h, the relative proportions of alternative splicing event types remained largely similar, with an additional contribution also from mutually exclusive exons (MX) (**Fig. 4E**).

For instance, 6 h following NCBP1-HNEylation, *thoc7* (encodes THO Complex subunit 7) underwent a statistically significant loss of alternative splicing event [skipping of exon 2 (SE)], leading to isoform-2 (**Extended Data Fig. 9F**). *Thoc7* encodes a functional spliceosome-associated protein integral to the THO complex that is involved in mRNA nuclear export. We validated this finding using qRT-PCR. Leveraging primers targeting *thoc7* isoform-2, normalized by results using primers targeting full-length (isoform-1) (**Supplementary Table 6**), the relative abundance of exon-2-lacking isoform-2 was significantly reduced following T-REX in HEK293T cells expressing NCBP1-Halo, compared to all 3 technical controls (**Extended Data Fig. 9G,H**). Replacing NCBP1-wt-Halo with NCBP1(C436)-Halo, ablated the observed reduced abundance of *thoc7* isoform-2 (**Extended Data Fig. 9I**). Similarly, *mybl2* (encodes *myb*-like protooncogene 2) underwent a statistically-significant gain of SE following NCBP1-HNEylation. This SE process produced isoform-2, that lacks exon-3 (**Supplementary Table 7,8**). We validated this SE-gain by qRT-PCR (**Extended Data Fig. 9J,K,L** and **Supplementary Table 6**). Notably, alternative splicing events of a significant number of genes associated with ribosomal translation were affected by NCBP1-HNEylation, including eukaryotic translation initiation factor 4E-binding protein 1 [4ebp1 (also known as elf4ebp1)]; several ribosomal-subunit proteins, e.g., rpl9, rpl32, and mitochondrial ribosomal-subunit proteins, e.g., mrpl13, mrpl23; and protein that binds hibernating ribosomes (ccdc124) (**Supplementary Table 7**).

### eIF2α and 4E-BP1 phosphorylation were unaffected

We also explored the extent to which key players driving translation initiation were involved in this process^36, 38, 39^. Following CBC-dependent translation initiation (CT) and NMD-associated quality control, steady-state translation cycles are driven by eIF4E-dependent 5’-cap-directed translation initiation (ET), as CBC is replaced by eIF4E. However, the functional boundaries between CT vs. ET are becoming increasingly blurred^36, 38, 39^. Particularly under stress, the CT-based initiation may be more operative^38, 39^. Nonetheless, both CT- and ET-supported initiation mechanisms require active eIF2α: both are suppressed by phosphorylation of eIF2α, a global stress-responsive means to rewire translational programs. However, additional stress-reponsive translational remodeling programs are emerging^36, 38, 39, 51–54^. A major mechanism includes hypophosphorylation of eIF4E-binding protein 1 (4E-BP1), which elicits ET inhibition with minimal effect on CT^38^. This mechanism likely explains heat-shock, late-stage hypoxia, and serum starvation-dependent translational responses, where CT is better able to support the cell translational capacity^36, 38, 39, 51–54^. We thus assessed the phosphorylation of 4E-BP1 and eIF2α post NCBP1-HNEylation. Neither was changed significantly (**Supplementary Fig. 26**).

### NCBP1-HNEylation promoted *s6k1* alternative splicing

These data led us to zoom into specialized factors unique to CBC-bound mRNA translation. Upregulation of CBC-dependent mRNA-translation is subsequently driven by a series of signaling actions involving activation of mTORC1 that phosphorylates ribosomal protein S6 kinase 1 (S6K1; also known as p70S6K), promoting the resultant activated p-S6K1’s recruitment and phosphorylation of SKAR (a component of EJC)^55^. Beyond translation initiation and quality control, CBC is emerging to function also in the steady-state translation, particularly during stress^38, 39, 51–53, 56^. Interestingly, our RNA-Seq data revealed that *rps6kb1* transcript (encoding protein S6K1) was outstandingly affected following NCBP1-HNEylation: Sashimi plots^57^ showed significantly-reduced intronic reads spanning exons 13 and 14; and also exons 12 and 13; but not for the intronic reads spanning upstream exons, such as exons 5 and 6; and exons 6 and 7 (**Fig. 5A**).

**Figure 5.**
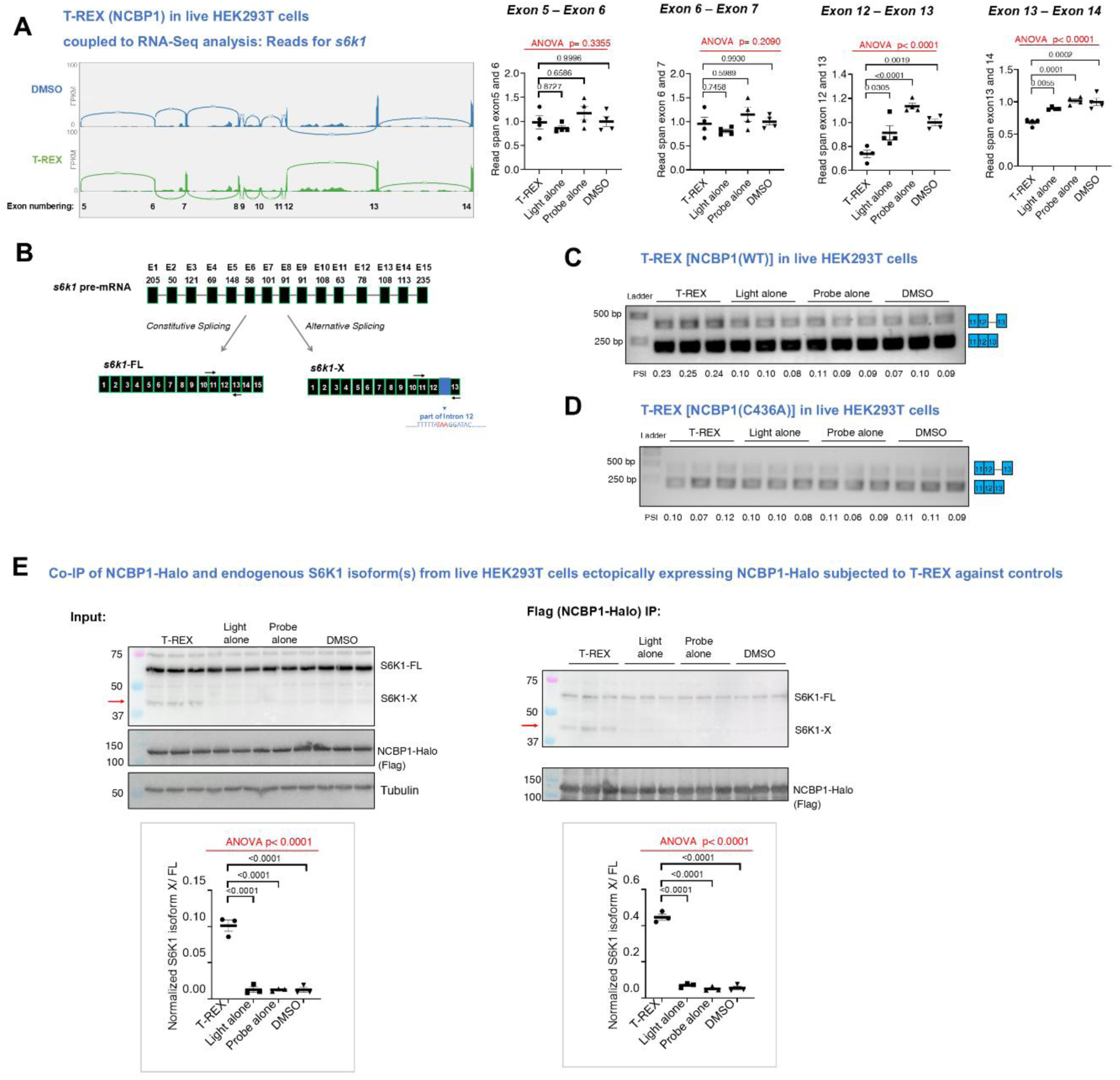
HNEylated NCBP1 generates an alternatively-spliced S6K1 protein dominantly inhibits translation. **A)** Representative Sashimi plots^57^ (*top*) and quantification (*bottom*), illustrating the reduction in reads for S6K1: exons 12 and 13; exons 13 and 14; following T-REX-assisted NCPB1-HNEylation against DMSO. No decrease was observed in preceding exons. p values were calculated with Tukey’s multiple comparison test. All data present mean±SEM (n=4). Genomic coordinates on Sashimi plot x-axes are omitted for clarity. Values within arcs depict number of junction reads. Sashimi plots are shown for only one out of 4 biological replicates (T-REX vs. DMSO). See plots below for quantification of read spans across indicated exons, across all datasets, and under T-REX vs. technical-controls. FPKM-fragments/kilobase transcript/million fragments mapped. **B)** Exon structure of S6K1 pre-mRNA (*top*-E1-E15, E indicates exon, and numbers above exons correspond to the no. base-pairs), and spliced forms: *bottom left*-S6K1-FL mRNA arising from constitutive splicing; *bottom right*-S6K1-X, an alternative-spliced isoform, formed upon NCBP1-HNEylation (this work). S6K1-X lacks E14 and E15, and partially lacks ‘E13’ due to alternative splicing at intron 12, which is partly retained. The pair of arrows shown for -FL and -X corresponds to data in **Fig. 5C,D**. See **Supplementary Fig. 27**. **C)** Following T-REX-enabled NCBP1-HNEylation, amplicon featuring Exon11-Exon12-Exon13 (*bottom band*-215 bp) and Exon11-Exon12-Intron12-Exon13 (*top band*-412 bp) were analyzed using primers in **Fig. 5B** (see **Supplementary Table 6**). PSI-proportion spliced in. Each band was PCR-amplified and validated by sequencing. n=3 each set. See **Supplementary Fig. 28**. **D)** Similar to **Fig. 5C** except HEK293T expressing NCBP1(C436A)-Halo. N=3 each set. **E)** *<u>Left panel</u>*<u>-</u>in HEK293T, NCBP1-HNEylation upregulated S6K1-X (45 kDa, red arrow). *<u>Right panel</u>*<u>-</u>results from Flag-immunoprecipitation (**Methods**). See **Extended Data Fig. 10A,B**. Ratio of S6K1-X:S6K1-FL in lysate versus post Flag-IP, normalized by NCBP1-Halo expression was calculated. p values were calculated with Tukey’s multiple comparisons test. All data present mean±SEM (n=3).

S6K1 has several splice variants, including a kinase-dead, full length (FL) variant, and alternative shorter isoforms arising from alternative events between Exons 6 and 7 (thereby resulting in the loss of Exons 8-15, along with modified/alternative Exon 7)^58^. The short isoforms from both mouse and human—identical to S6K1-FL up to Exon 6 but missing more than half of the conserved kinase domain—are considered catalytically inactive. Interestingly, the short isoforms manifest context-specific oncogenic behaviors, whereas the kinase-dead FL variant is tumor-suppressive^58, 59^. Our analysis (**Fig. 5A**) showed that NCBP1-HNEylation promoted an alternative exon-skipping event, distinct from those reported (**Fig. 5B**, **Supplementary Fig. 27**). Specifically, NCBP1-HNEylation produced S6K1-X that lacks exons 14 and 15; and has an alternatively-spliced intron 12 (**Supplementary Fig. 27**). qPCR analysis of endogenous S6K1, following NCBP1-HNEylation in HEK293T cells, confirmed selective build-up of S6K1-X-associated amplicon that includes part of intron 12 (264 bp), under T-REX, compared to all 3 controls (**Fig. 5C**). A significantly-increased ‘proportion spliced-in (PSI)^47^’ index was found compared to the amplicon lacking the above-mentioned intron-12 constituent. Both amplicons were further validated by sequencing (**Supplementary Fig. 28**). Notably, when NCBP1(C436A) replaced NCBP1(wt) in the same qPCR assay post T-REX under otherwise identical conditions, HNE-sensing-active but HNE-signaling-deficient NCBP1(C436A) did not produce S6K1-X (**Fig. 5D**). We proceeded to investigate the functional ramifications of the resulting S6K1-X-protein.

### NCBP1-HNEylation produced a truncated S6K1-protein, S6K1-X

We examined how NCBP1-HNEylation modulates S6K1-FL and S6K1-X as a function of time, and how HNEylated NCBP1 associates with the two S6K1 variants by co-IP. S6K1-X was detected by western blot analysis as early as 2 hours post T-REX-mediated NCBP1-HNEylation. S6K1-X was detected at ~45 kDa by an antibody that binds in the N-terminal region (**Extended Data Fig. 10A,B,C**). This band was absent in all controls: an antibody that binds S6K1 in the C-terminal region could not detect S6K1-X, although it could detect S6K1-FL. S6K1-X associated with HNEylated NCBP1 more effectively than S6K1-FL (**Fig. 5E**). However, this association was sensitive to RNAse (**Supplementary Fig. 29**), indicating that S6K1-X associates with RNA (or another RNA-binding protein). Imaging of HEK293T cells ectopically expressing either S6K1-X or S6K1-FL showed largely cytoplasmic localisation (**Extended Data Fig. 10D**).

### S6K1-X inhibited translation

Given that the level of S6K1-X produced upon NCBP1-HNEylation was modest (**Fig. 5E** and **Extended Data Fig. 10A,B**), we posited that S6K1-X exerted dominant-negative effects on translation. To test this hypothesis, we compared effects of either S6K1-X, or S6K1-FL overexpression in HEK293T cells, using cells transfected with an empty vector (EV) as control. Consistent with our hypothesis, overexpression of S6K1-X inhibited translation, whereas S6K1-FL had no effect compared to EV control (**Fig. 6A**, **Extended Data Fig. 10C**). Should the effect of NCBP1 HNEylation be mediated through S6K1-X, increased S6K1-X production by T-REX to NCBP1 in cells already expressing this inhibitory splice variant should have minimal effect on translation. In cells expressing S6K1-FL or EV, T-REX-assisted NCBP1-HNEylation caused translation inhibition, as previously observed. However, in cells expressing S6K1-X, NCBP1 HNEylation had no effect (**Supplementary Fig. 30**). These outcomes are consistent with S6K1-X eliciting a dominant-negative effect on translation, mediated by NCBP1-HNEylation.

**Figure 6.**
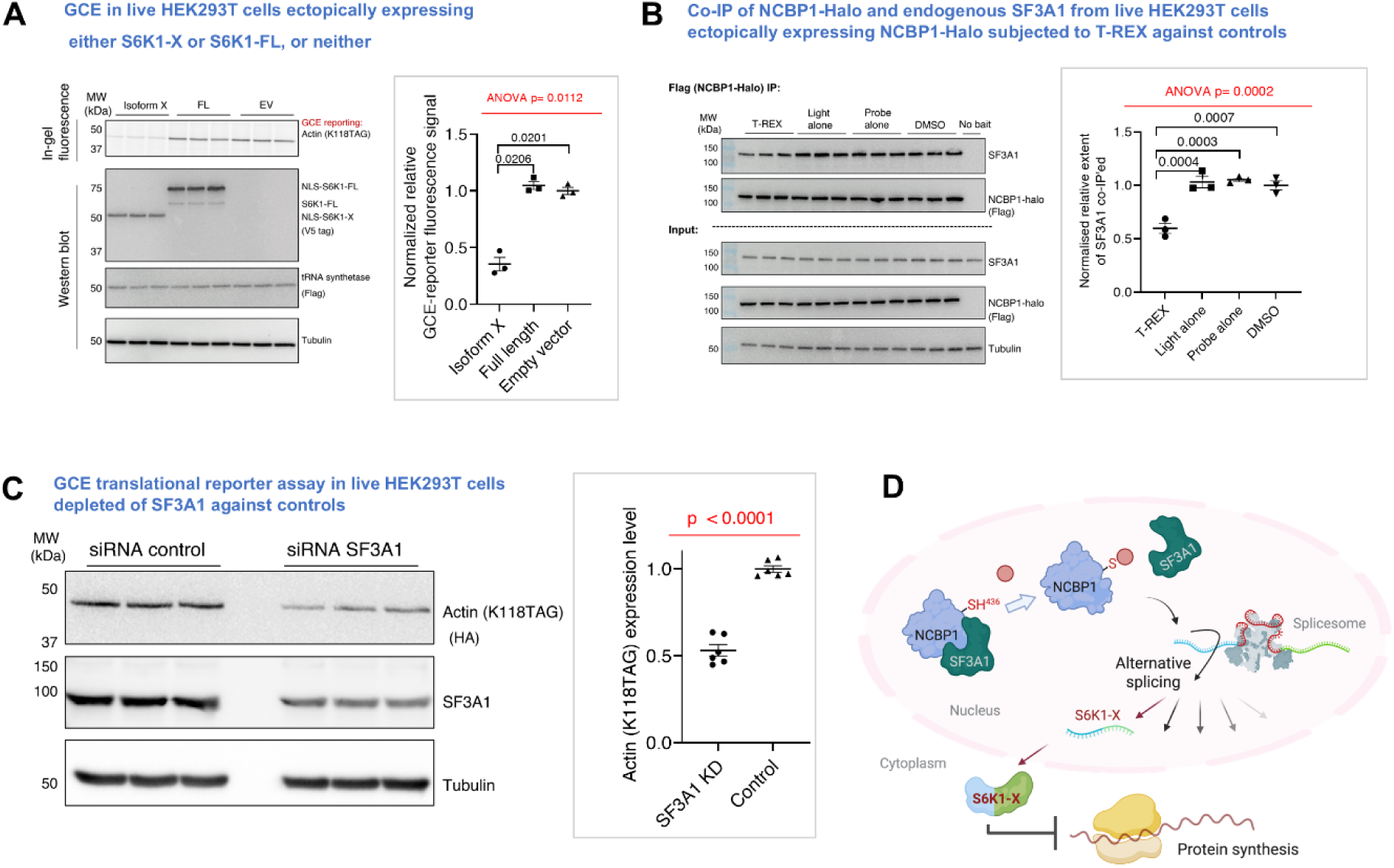
S6K1-X overexpression stalls translation; SF3A1 is necessary for NCBP1(C436)-HNEylation-dependent S6K1-X formation. **A)** S6K1 -X dominantly suppresses translation. HEK293T overexpressing S6K1-FL, or S6K1-isoform-X or empty vector (EV), replete with GCER constructs, described elsewhere, were treated with BCNK (50 µM, 4 h). Post lysis, and coupling with tetrazine-Cy5, samples were analyzed by in-gel Cy5-fluorescence reporting protein translation. Anti-V5-tag (detecting S6KL), anti-Flag (tRNA synthetase PylRS), and anti-tubulin (loading control) blots were also performed. Ectopic S6K1 and S6K1-X each have two forms, one with and one without nulcear-localization sequence (NLS). Thus the blot shows: NLS-S6K1-FL (74 kDa, also called p85), S6K1-FL (59 kDa), NLS-S6K1-X (51 kDa), S6K1-X (45 kDa). See **Extended Data Fig. 10C**. *<u>Inset</u>*-quantification. *p* values were calculated with Tukey’s multiple comparison test. Data present mean±SEM (n=3). See **Supplementary Fig. 30**. **B)** In HEK293T, NCBP1 was HNEylated and then anti-Flag pulldown was performed with results compared to T-REX controls (**Methods**). Input and elution samples were analyzed by western blot as indicated. *<u>Right panel</u>*-quantification of eluted binding partner protein indicated, normalized by NCBP1-Halo expression and tubulin. p values were calculated with Tukey’s multiple comparison test. All data present mean ± SEM. See Methods for sample size information. **C)** HEK293T were co-transfected with GCE(actin)-reporter and SF3A1 siRNA (or control siRNA). 36 h later, cells were treated with BCNK (50 µM, 4 h). Post cell lysis and tetrazine-Cy5-Click, samples were analyzed using in-gel fluorescence reporting protein translation efficiency. Western blot was performed using anti-HA [reporting Actin (K118TAG)], anti-SF3A1, and anti-tubulin (loading control). p values were calculated by an unpaired, two-tailed t-test. Data present mean±SEM (n=6). **D)** NCBP1(C436)-HNEylation induced significant alternative splicing (AS) across >250 genes (see also **Supplementary Table 8** and **Fig. 5E**). One such eventformed S6K1-x. S6K1-x dominantly inhibited translation. AS occurs as a result of NCBP1-HNEylation reducing association with SF3A1, a key component of splicesosome. All full-view blots and gels are shown in **Source Data Fig. 1-5**.

### SF3A1 mediated S6K1-X formation

All these data pointed to a mechanistic model where NCBP1(C436)-specific HNEylation in the nucleus triggers a broad-spectrum perturbation of alternative splicing (AS) processes, including the formation of S6K1-X, which inhibits translation. S6K1-X’s primarily cytoplasmic location (**Extended Data Fig. 10D**) and its ability to associate with mRNA (**Supplementary Fig. 29**) are also consistent with this model. We next explored upstream factor(s) regulating splicing or transcription-export modulated by NCBP1-HNEylation. We focused on four known or putative NCBP1 interactors having these functions, namely, ALYREF, PHAX, PRPF4 and SF3A1. The first and second proteins are key transcription-export factors for mRNAs and snRNA/snoRNAs; the third and fourth are essential splicesome components. T-REX coupled with co-IP assays revealed that NCBP1-HNEylation caused no perturbation of the association between NCBP1 and ALYREF, PHAX, or PRPF4 (**Supplementary Fig. 31**). However, the same process selectively impaired the SF3A1—NCBP1 interaction (**Fig. 6B**). This result is remarkable because unlike ALYREF, PRPF4, and PHAX, no direct evidence of NCBP1’s association with SF3A1 has been demonstrated.

We proposed that NCBP1-HNEylation would reduce interaction with SF3A1, impacting splicing and inhibiting translation. Consistent with this postulate, NCBP1-HNEylation-driven translation suppression was ablated in SF3A1-depleted cells compared to KD-control (**Supplementary Fig. 32**). In SF3A1-deficient HEK293T cells, translation efficiency was reduced compared to KD-control (**Fig. 6C**), a result implying that SF3A1 promotes translation, beyond being important in splicing. SF3A1 silencing is thus epistatic with NCBP1-HNEylation in terms of translation suppression.

### S6K1-X is observed in Huntington’s diseased proteomes

To assess the extent to which these findings extend to disease states, we examined models of neurodegeneration, that often show elevated levels of protein HNEylation^8, 60^. In lines overexpressing a Huntington (Htt) disease-causative constructs, namely, Htt Q109 harboring extended polygutamine (poly-Q) mutations, significant levels of endogenous S6K1-X was observed, compared to non-diseased control cells overexpressing Htt Q15. As expected, the former diseased cells further manifested enhanced HNEylated proteomes (**Extended Data Fig. 10E**).

## Discussion

This study presents a hitherto unrecognized mechanism to stall protein synthesis during reactive small-molecule stress in the nucleus. Our data indicate that endogenous electrophilic stress accumulation in the nucleus is predominantly sensed and managed by the large subunit of the CBC, NCBP1, resulting in rapid suppression of translation ~2 h following nuclear-localized electrophile build-up. Such a mechanism provides a new means for stress upregulation in the nucleus to downregulate translation^36, 61^. Such a regulatory process is arguably needed because translation is thought to be largely uncoupled from events in the nucleus. We thus establish a clear, coordinated response to nuclear electrophilic stress, distinct from the cytosolic stress response, that martials upregulation of transcription of specific cytoprotective genes. As the nucleus is relatively protected from external reactive small-molecule agents, it is likely that this translation suppression response represents the fact that the build-up of reactive chemicals in the nucleus is mutagenic and potentially highly damaging.

The parallels between this mechanism and translation inhibition enacted by ER stress (usually referred to as the unfolded protein response) are not lost on us. In general ER stress responses lead to a general decrease in translation to allow coordinated response to stress. Both also use an unconventional splicing event that helps coordinate the response (S6K1 in the case of nuclear specific stress sensing; XBP1 during ER-stress^62^). However, several factors differentiate our mechanism from traditional stress-response pathways, and highlight the key factors of how substoichiometric electrophile signaling differs from conventional signaling pathways. In our case, NCBP1 is both the initial sensor of electrophile stress and the mediator of S6K1 differential splicing. This mode of the sensor/writer also being the reader is common for reactive small-molecule signaling. In ER stress response, a wealth of protein factors martials translational suppression responses. In the case of NCBP1, its electrophile-labeling leads to a change in splicing, mediated through association changes with SF3A1, producing a dominant-negative suppressor of translation, S6K1-X.

Although the response mode is particularly intriguing and sets new paradigms on how to martial stress, it shoud be noted that our identified electrophile-responder protein is itself enigmatic. Indeed, NCBP1 emerged to be a sensor of electrophilic stress on par with the best sensors we have characterized. What we did not expect was that none of its cysteines were essential for electrophile sensing (engagement of the electrophile with NCBP1 cysteines); however, only one of its cysteines (C436) was responsible for signaling (changing flux through specific pathways, ushering a phenotype). Given that aberrant translation can play important roles in disease^63–65^, and that we have identified a new electrophile-responsive player linked to this process, these data altogether introduce a new privileged sensor ideal for drug target mining^19, 66^. Future work will address this aspect, as well as how the other proteins identified in our screen fit into the nuclear electrophile response.

## Acknowledgements

Drs. Krishna N. Tripathi, Amogh Kulkarni, Kevin T. Fridianto, and Chaosheng Luo (Aye Lab) for the syntheses of Ht-PreHNE, Ht-PreHNE(alkyne), and HNE(alkyne). Mr. Surased Suraritdechachai (Uttamapinant lab), Riyaz Khan (NCCR master thesis student, Aye Lab), and Mr. Tony Hu (ThinkSwiss Scholar, Aye Lab) for assistance with gene cloning. Dr. Chloé Rogg (Aye Lab) for initial training of M.A. and assistance during exploratory investigations. Prof. Frances Platt (Head of the Department of Pharmacology, Oxford University) for provision of an interim tissue-culture room. Drs. Romain Hamelin and Maria Pavlou (EPFL proteomics core facility) for assistance with LFQ-DDA differential protecomics data processing and analysis. Drs. Viviane Praz, Hannes Richter, and Julien Marquis (University of Lausanne Genomic Technologies Facility) for assistance with RNA-Seq data processing and analysis. Dr. Ashley Sawle (Principal Bioinformatics Analyst, Cancer Research UK Cambridge Institute) for differential splicing analysis of RNA-Seq data. Dr. Marjorie Fournier and Vaishnavi Ravikumar (Advanced Proteomics Facility, Oxford Biochemistry) for PTM-mapping analysis. <u>Research</u> <u>support:</u> Swiss National Science Foundation (SNSF) SPIRIT (grant no. 193915) [to Y.A. (PI) and C.U. (co-PI)]; SNSF R-Equip Funding for proteomics research instrumentation (grant no. 316030_213435) [to Y.A. (PI) and Dr. Maria Pavlou (co-PI)]; and Academy of Medical Sciences Professorship (National Academy of Medical Sciences UK) and Department for Science, Innovation & Technology UK (DSIT) (grant no. APR11\1013) (to Y.A.). Novartis Foundation for Medical-Biological Research Postdoctoral Fellowship (to M.J.C.L.).

## Author Contributions

D.C.: performed all experiments, data compilation, analysis, and writing (figure legends and supporting information); M.A.: optimizations toward successful establishment of REX—GCE dual-technology integration; C.S.: collaborated with D.C. in independent validations of REX—GCE dual-technology; K.S.: collaborated with D.C. during initial developments of REX—GCE dual-technology; C.U., SNSF:SPIRIT project grant co-PI: funding acquisition; grant administration; GCE-technology technique transfer, experimental design, and consultation; data analysis; M.J.C.L.: experimental design and data analysis, writing (manuscript critical review and editing); Y.A.: conceptualization, experimental design and data analysis, writing (manuscript draft, critical review, and editing). All authors assisted with final proofing of the manuscript.

## Online Methods Section

### Plasmid cloning

Ligase-free cloning^16, 20^ was used unless specified otherwise. **Supplementary Tables 1** and **2** list primers, and sequences of localization signals used. All constructs were validated by sequencing.

pPB-MmPylRS-AF-4xPylT_CUA_ and pPB-HA-actin(K118TAG)-4xPylT_CUA_ were gifts from Jason Chin (MRC LMB)^12^. psGG_Nrf2-intron^29^ was digested with AflII (NEB) to insert exon10-intron10-exon11, or exon10-exon11, then digested with Xhol (NEB) to insert DACH1 3’UTR. pCS2+8-NCBP2-HA, pCS2+8-S6K1-FL-V5, and pCS2+8-S6K1-X-V5, were constructed by amplification of NCBP2, S6K1-FL (EPFL Gene Expression Core Facility), and S6K1-X from cDNA after reverse transcription of mRNA isolated from HEK293T using primer oligo(dT)20. All primers were from IDT DNA.

lentiCRISPRv2 gRNA-1 and -2 for DACH1 knockout line generation were constructed by insertion of DACH1(gRNA) into lentiCRISPRv2 (Addgene-#52961). DACH1(gRNA1)-AACCTGGCGGCCGCGAGCAA; DACH1(gRNA2)-GCCACAGTCACCTCTACCGG; Control(gRNA)-CTTCGAAATGTCCGTTCGGT. SF3A1(siRNA) & control were from Dharmacon.

### SDS-PAGE and western blot analysis

SDS-PAGE/blotting were performed as described^20^. For primary and secondary antibodies, see **Supplementary Table 9**.

### Cell culture

HEK293T were cultured in MEM+10% FBS, penicillin/streptomycin, sodium pyruvate, and non-essential amino acids. N2A were cultured in DMEM plus the same additives at 37°C in a humidified 5% CO_2_ atmosphere. N2A were differentiated with retinoic acid (20 µM, 24 h) in DMEM +2% FBS, sodium pyruvate, non-essential amino acids, and penicillin/streptomycin. Both cell lines were from ATCC and were validated mycoplasma -free using the mycoplasma PCR detection kit (Thermo).

### Knockout

HEK293T in a 10 cm dish at ~90% confluence were transfected with 7.5 μg pCMV-R8.74psPAX2, 750 ng pCMV-VSV-G and 7.5 μg plentiCRISPRv2 using TransIT-LT1 (Mirus Bio) following the manufacturer’s protocol. 48 h post transfection, 10 mL supernatant was collected and filtered (0.45-micron). Then 5.8 mL PBS, 325 μL 5M NaCl, and 2 mL PEG-8000 (50% in PBS) were added, and incubated overnight (4°C) with end-over-end rotation. The supernatant was discarded post-centrifugation (1,500g,1 h, 4°C). Pellet was resuspended in 100 μL PBS.

HEK293T in a 6-well plate ~90% confluence were treated with 25 μL lentiviral particles in 2.5 mL media, containing 8 μg/mL polybrene. After 24 h, media were changed, and cells were incubated for 24 h. Cells were then incubated in media containing 2 μg/mL puromycin.

### Localis-REX–GCE in differentiated N2A cells

N2A were seeded in a 12-well plate for in-gel fluorescence, and transfected with pCS2+8-NLS-Halo, pCS2+8-NES-Halo, pCS2+8-MOMLS-Halo, or pCS2+8-ERt-Halo, Actin(K118TAG), MmPylRS RNA synthesase (plasmid raito-1:10.2) and Lipofectamine3000 reagent as per the manufacturer’s recommendation. For whole proteomic labeling with BCNK, Val-MmPylRS RNA synthease was used instead of Actin(K118TAG) and MmPylRS RNA synthease (plasmid ratio of Halo:Val-MmPylRS is 1:0.2). 12 h later retinoic acid (20 µM) was added (24 h) to differentiate N2A. Cells were incubated with Ht-PreHNE (10 µM, 2 h), and rinsed thrice. Localis-REX: cells were exposed to light (2 min, 5 mM/cm^2^, 365 nm); light-alone: cells incubated with DMSO. were exposed to light (2 min, 5 mM/cm^2^, 365 nm); Pre: cells were incubated with Ht-PreHNE (10 µM, 2 h); DMSO control: cells were incubated with DMSO. 5 min post light exposure (if used), all cells were incubated with BCNK (60 µM, 4 h), washed with cold DPBS twice, then frozen in liquid nitrogen.

Localis-REX and T-REX controls are defined as follows:

|  | Ht-PreHNE | DMSO | Light |
| --- | --- | --- | --- |
| Light alone | - | + | + |
| Probe alone | + | + | - |
| DMSO | - | + | - |

### Localis-REX in differentiated N2A cells for function-guided spatial target-ID proteomics analysis

N2A were seeded in a 60-mm dish. 24 hrs later ~80% confluent cells were co-trasfected with pCS2+8-NLS-Halo and Lipofectamine3000 as per the manufacturer’s recommendation. 12 h later, cells were treated with retinoic acid (20 µM, 24h) to differentiate N2A. Cells were incubated with Ht-PreHNE(alkyne-functionalized) (15 µM, 2 h), then rinsed thrice. Cells were exposed to light (5 min, 5 mM/cm^2^, 365 nm). Control experiments were treated identically except Ht-PreHNE(non-alkyne-functionalized) replaced Ht-PreHNE(alkyne-functionalized). Cells were washed with cold DPBS twice, then frozen in liquid nitrogen.

### BONCAT experiment

HEK293T were seeded in 12-well plates (for Cy5-Click assay) or 60 mm dishes (for biotin-Click pulldown assay). After 24 hours, cells at ~80% confluence were transfected with either pCS2+8-NLS-Halo, pCS2+8-NCBP1-Halo, or pCS2+8-NCBP1(C436A)-Halo as indicated in corresponding figure legends. TransIT2020 was used as per the manufacturer’s recommendation. After 36 h, cells were incubated with Ht-PreHNE(no alkyne) (15 µM, 2 h), and rinsed thrice. During the second wash, the culture medium was changed to DMEM without methionine (ThermoFisher, 21013024) and incubated (30 min). At the indicated time post light exposure, all experimental groups were incubated with L-Azido-homoalanine (AHA, 50 µM, 4 h) in DMEM without methionine. Cells were washed with cold DPBS twice, and flash-frozen in liquid nitrogen.

### T-REX–GCE in HEK293T cells

HEK293T were seeded in 12-well plates, 24 h later cells at ~80% confluence were co-transfected with pCS2+8-NCBP1-Halo, pCS2+8-Cand1-Halo or pCS2+8-IPO5-Halo, Actin(K118TAG), MmPylRS RNA synthesase (raito: 1:1; TransIT2020). 36 h later, cells were incubated with Ht-PreHNE(no alkyne) (15 µM, 2 h), then rinsed thrice. Post light exposure, all cells were incubated with BCNK (60 µM, 4 h). Cells were washed twice with cold DPBS, then frozen in liquid nitrogen.

### T-REX in HEK293T cells

HEK293T were seeded in either 12-well plates (for in-gel fluorescence/western blot), 6-well plates (for RNA-Seq), or 60 mm dishes (for Flag-pull down). 24 h later, cells at ~80% confluence were co-transfected with pCS2+8-NCBP1-Halo, pCS2+8-Cand1-Halo or pCS2+8-IPO5-Halo, and TransIT2020. 36 h later, cells were incubated with Ht-PreHNE(alkyne) (15 µM, 2 h), and rinsed thrice, then exposed to light. At indicated times post light exposure, cells were washed with cold DPBS twice, then frozen in liquid nitrogen.

### Biotin-Click and streptavidin pull-down

Cells were harvested following the steps in **Localis-REX in differentiated N2A cells or T-REX in HEK293T cells**, and lysed (50 mM HEPES, 150 mM NaCl, pH 7.6, 1% Nonidet P-40 and 1X Roche cOmplete, mini, EDTA-free protease inhibitor cocktail by freeze-thaw thrice). Lysate was centrifuged (20,000 g,10 min, 4 °C). Protein concentration was normalized (Bradford assay; BSA-standard) to 1 mg/mL using 50 mM HEPES (pH 7.6) and 0.3 mM TCEP. In T-REX workflow, the samples were subjected to TeV protease (0.2 mg/mL, 30 min, 37°C) as indicated in figure legends. TeV cleavage was not applicable to Localis-REX. In both workflows, samples were then subjected to Click reaction/streptavidin pull-down as described previously^6^.

### Immunofluorescence

N2A cells were seeded in 35-mm glass-bottomed dishes. 24 h later cells at ~40% conflurence, were trasfected with pCS2+8-NLS-Halo, pCS2+8-NES-Halo pCS2+8-MOMLS-Halo, or pCS2+8-ERt-Halo, and Lipofectamine 3000. After 5 h, cells were incbuated with retionic acid (20 µM) in DMEM supplemented with 2% FBS (24 h). Then, cells were treated with Ht-PreHNE (alkyne) (15 µM, 2 h), then rinsed thrice. Cells were treated with MitoTracker-Red (100 nM, 30 min), then rinsed twice. Cells were stained as previously described. Images were taken using a Nikon spining-disk confocal microscope equipped with a 100X objective.

### Mass spectrometry analysis

#### (i)#Sample preparation for LC-MS/MS

Samples obtained through ‘biotin-Click, streptavidin pull down’ were separated by SDS-PAGE on a 12% polyacrylamide gel and Coomassie blue stained. Each gel lane was sliced, and proteins were digested. The gel pieces were washed twice with 50% ethanol in 50 mM ammonium bicarbonate (AB, Sigma-Aldrich, 20 min) and dried by vacuum centrifugation. Proteins were reduced (10 mM dithioerythritol, Merck-Millipore, 1 h, 56°C) followed by a washing-drying step as described above. Proteins were alkylated (55 mM iodoacetamide, Sigma-Aldrich, 45 min, 37°C), followed by a washing-drying as described above. Proteins were digested overnight at 37°C using MS grade Trypsin Gold (Promega, 12.5 ng/µl in 50 mM AB supplemented with 10 mM CaCl_2_). Peptides were extracted (70% ethanol, 5% formic acid (Merck-Millipore)) twice (20 min) then dried by vacuum centrifugation. Resulting peptides were desalted on StageTips^67^ and dried under a vacuum concentrator.

#### (ii)#Mass Spectrometry (MS)

Samples were resuspended in 2% acetonitrile (Biosolve), 0.1% FA and nano-flow separations were performed on a Dionex Ultimate 3000 RSLC nano UPLC system (Thermo Fischer Scientific) on-line connected with an Exploris 480 Orbitrap Mass Spectrometer or a Lumos Orbitrap Mass Spectrometer (Thermo Fischer Scientific). A capillary precolumn (Acclaim Pepmap C18, 3 μm-100Å, 2 cmx75 μm ID) was used for sample trapping and cleaning. A 50cm long capillary column (75μm ID; in-house packed using ReproSil-Pur C18-AQ 1.9 μm silica beads) was then used for analytical separations at 250 nl/min over 150 min biphasic gradients. Acquisitions were performed through Top Speed Data-Dependent acquisition. The most intense parent ions were selected and fragmented by High-energy Collision Dissociation (HCD) with 30% Normalized Collision Energy (NCE). Selected ions were then excluded for the following 20 s.

#### (iii)#Label-Free Quantification (LFQ)

Raw data were processed using MaxQuant 1.6.10.43 4^68^ against the *Mus Musculus* Reference proteome database (55341-protein sequences Release2021_02), Halo Protein sequence, or the *Homo Sapiens* Reference proteome database (77027-protein sequences Release2021_01). Carbamidomethylation was a fixed modification, whereas methionine oxidation; serine, threonine, tyrosine phosphorylation; N-termal acetylation; and glutamine to pyroglutamate were considered variable modifications. <3 missed cleavages were allowed, and the “Match between runs” option was enabled. A minimum of 2 peptides was required for protein identification and false discovery rate (FDR) cutoff was 0.01 for both peptides and proteins. Label-free quantification and normalisation were performed by Maxquant using the MaxLFQ algorithm, with standard settings^69^.

Statistical analyses of the label-free data were performed using Perseus v1.6.15.0 [doi: 10.1038/nmeth.3901.] (MaxQuant suite). Reverse proteins, potential contaminants and proteins only identified by sites were filtered out. Protein groups containing at least 2 valid values in at least one group were conserved for further analysis. Empty values were imputed with random numbers from a normal distribution (Width: 0.3 and Down shift: 1.8). A two-sample t-test with permutation-based FDR statistics (250 permutations, FDR = 0.01) was performed to determine significant differentially abundant candidates. Localis-REX-derived LFQ-MS time-resolved proteomics datasets are summarized in **Supplementary Table 3**.

### Anti-Flag pulldown

HEK293T were seeded in a 60 mm dish; the rest of the steps followed the protocol, **T-REX in HEK293T**. At indicated times post light exposure, cells were washed with cold DPBS twice, then frozen in liquid nitrogen. For immunoprecipitation of p-s6k1 bound by NCBP1, 1 h post T-REX, cells were treated with insulin (100 nM, 1 h) before harvesting. Cells were lyzed with NET-2 buffer (50 mM Tris-Cl, 100 mM NaCl, 0.1% NP-40, pH 7.4) with 1 mM PMSF, 1mM Na_3_VO_4_, 10 mM NaF, 25 nM Calyculin A, 1X Roche Protease inhibitor tablet, and 100 U/mL RNAse OUT inhibitor) followed by rapid freeze-thaw cycles (x3). Flag-pulldown with anti-Flag M2 affinity agarose gel (A2220, Sigma) was performed as described^6^.

### In-gel fluorescence assay

#### (i)#For GCE

Cells from one well of a 12-well plate were lysed in 30 µL buffer containing 50 mM HEPES (pH 7.6), 150 mM NaCl, 1% Nonidet P–40, 1X Roche cOmplete, mini, EDTA-free protease inhibitor cocktail, and 0.3 mM TCEP by freeze-thaw (x3). Cells debris was removed by centrifugation (18,000g, 8-10 min, 4 °C). Lysate concentration was determined using Bradford assay (standard-BSA) and diluted to 1 mg/mL, and incubated with Tetrazine-Cy5 (1 µM, 30 min, 37°C), then quenched with 5 µL 4X Laemmeli dye containing 6% βME. After 5-min incubation at 37 °C, the lysate was subjected to SDS-PAGE. After electrophoresis, the gel was rinsed 3X with ddH_2_O (5-min each) and imaged on a Biorad Chemidoc-MP Imager. Then the gel was transferred to a PVDF membrane for western blot analysis.

#### (ii)#For T-REX

All steps in this section were performed under red light unless otherwise stated. In -gel fluorescence assay for T-REX was performed as described^6^.

#### (iii)#For gel-based BONCAT analysis

Cells from one well of a 12 well plate were lysed using freeze-thaw (x3), in 30 µL 50 mM HEPES (pH 7.6), 150 mM NaCl, 1% Nonidet P–40, 1X Roche cOmplete, mini, EDTA-free protease inhibitor cocktail, and 0.3 mM TCEP. Cell debris was removed by centrifugation (18,000g, 8-10 min, 4°C). Lysate concentration was determined using Bradford assay (BSA-standard) and diluted to 1 mg/mL then subjected to biotin-Click reaction (final reaction mix-1.7 mM TCEP, 5% *t*-BuOH, 1% SDS, 1 mM CuSO_4_, 0.1 mM Cu(TBTA), 10 µM Cy5-alkyne). The samples were incubated (37 °C, 30 min) then quenched with 4X Laemmli dye, 6% βME. Post 5-min incubation (37 °C) the lysate was subjected to SDS-PAGE, washed with ddH_2_O (3x) then imaged on a Biorad Chemidoc-MP Imager. Gel was transferred to a PVDF membrane for western blot analysis.

### Luciferase reporter assays

HEK293T cells were transfected with NCBP1-Halo or NCBP1 C436A-Halo, intron- or intron-less firefly luciferase reporter, and internal control CMV-Renilla luciferase (in plasmid ratios: 1:1:0.025). Upon T-REX (as shown above), Luciferase reporter Assays were carried out as previously described^29^.

### RNA sequencing

#### Experimental setup and RNA purification

HEK293T cells were seeded in 6-well plates for 24 h. Cells at ~80% confluence were co-transfected with pCS2+8-NCBP1-Halo, pCS2+8-Cand1-Halo or pCS2+8-IPO5-Halo, using TransIT 2020 reagent per the manufacturer’s recommendation. 36 h later, T-REX was performed, followed by RNA purification^29^.

#### RNA-Seq analysis

Sample preparation and analyses were performed by The Genomic Technologies Facility (GTF) at the University of Lausanne. Libraries were prepared from 250 ng total RNA with TruSeq Stranded mRNA reagents (Illumina). The polyA selection step was replaced by an rRNA depletion step with the QIAseq FastSelect-rRNA HMR kit (Qiagen). Library preparation was performed on a Sciclone liquid handling robot (PerkinElmer; Waltham, Massachusetts, USA) with a PerkinElmer-developed automated script. Unique dual indexes were used for the barcoding of the libraries. Libraries were quantified (QubIT, Life Technologies), and quality was assessed on a Fragment Analyzer (Agilent Technologies). Sequencing was performed on an Illumina NovaSeq 6000 for 300 cycles (paired-end 150 nt reads). Sequencing data were demultiplexed using bcl2fastq2 Conversion Software (version 2.20, Illumina). T-REX(NCBP1)-mediated RNA-seq datasets are summarized in **Supplementary Table 4**.

### Differential splicing analysis

Alternative splicing analysis was performed by CRUK Cambridge Institute (Dr. Ashley Sawle), using SUPPA2 (https://github.com/comprna/SUPPA)^70^. The transcript counts from the raw datasets from RNA-Seq analyses (above) were estimated using Salmon^71^, resulting in 1266 genes to be further analyzed. Counts were normalized to transcripts per million (TPM), prior to further analysis with SUPPA2. Transcript usage and alternative splicing events were measured as “proportion spliced in” (PSI), i.e., for a given event, the total estimated counts for transcripts including the event were divided by the total estimated counts for all transcripts that could include the event. The list of alternative splicing events with full information (GeneID, PSI of each transcript under each condition, dPSI of difference in the mean PSI between two conditions, each Alternative splicing event and its detail, adj *p*-value) were summarized in **Supplementary Table 7** and **8**. ‘Loss’ and ‘gain’ respectively indicate splicing events that are less common / lost or more common / gained under the T-REX condition, with respect to indicated control.

### Quantitative real-time PCR (RT-qPCR)

RT-qPCR^29^ and analysis^12^ were carried out as previously described.

### RNA Half-life

HEK293T expressing NCBP1-Halo were conducted with T-REX and all other three controls, then performed as described. Data were fit to one-phase exponential decay: y=(1-Plateau)*exp(-*k*X)+Plateau; y-RNA transcript levels, *k*-decay constant, and x-time, (Prism v9.4.0).

### Mass spectrometry analysis of HNEylation site

Following the steps described in **T-REX in HEK293T cells**, cell lysis (3x freeze-thaw) was performed in 500 μL (50 mM HEPES (pH 7.6), 100 mM NaCl, 1% Nonidet P-40, 10 mM Imidazole, 5 mM βME and Roche protease inhibitor). Lysate was clarified by centrifugation (20,000g, 10 min, 4 °C). Protein concentration was determined by Bradford assay(BSA-standard) and diluted to 1.5 mg/mL in lysis buffer and incubated with 50 μL TALON resin (pre-equilibrated with lysis buffer, 1 h, 4°C). Resin was washed thrice (5 min, 500 μL wash buffer-50 mM HEPES (pH 7.6), 100 mM NaCl, 0.5% Nonidet P-40, 20 mM imidazole and 5 mM βME), then proteins were eluted with 30 μL elution buffer (50 mM HEPES, 100 mM NaCl, 200 mM Imidazole, 5 mM βME). Samples were subjected to SDS-PAGE and stained with colloidal blue. The band of NCBP1-Halo was cut and into cubes of ~ 1×1 mm and washed (2x) in Milli-Q water, destained in 1:1 v/v 0.1 M ammonium bicarbonate (AmBic):acetonitrile (ACN) (30 min, 24 °C). Destained gel pieces were dehydrated with ACN, then reduced in 10 mM TCEP (Tris-(2-carboxyethyl)phosphine)/0.1 M AmBic (30 min, 24 °C) and alkylated in 50 mM 2-chloroacetamide/0.1 M AmBic (30 min, 24 °C). The gel pieces were dehydrated and then digested (16-18 h, 37 °C) with endoprotease Trypsin (Promega) [10 ng/µL trypsin/10 mM AmBic/10 % ACN]. Digested peptides were extracted twice from the gel (5 % formic acid/ACN (1:2 v/v), 15 min, 37 °C) and dried by vacuum centrifugation.

Peptides were separated by Ultimate RSLC 3000 nano liquid chromatography (Thermo Scientific), coupled in line to a Q Exactive mass spectrometer equipped with an Easy-Spray source (Thermo Scientific). Peptides were trapped onto a C18 PepMap™ 100 precolumn (300 µm i.d. x 5 mm, 100 Å, Thermo Scientific) using Solvent A (0.1 % formic acid in HPLC grade water). Peptides were further separated onto an Easy-Spray RSLC C18 column (75 µm i.d., 50 cm length, Thermo Scientific) using a 40 min linear gradient (15% to 38% solvent B (0.1 % formic acid in ACN)), flow rate 200 nL/min. Raw data were acquired in data-dependent acquisition mode (DDA). Full-scan MS spectra were acquired in the Orbitrap (Scan range 350-1500 m/z, resolution 70,000; AGC target, 3e6, maximum injection time, 50 ms). The 5 most intense peaks were selected for higher-energy collision dissociation (HCD) fragmentation at 30% of normalized collision energy. HCD spectra were acquired in the Orbitrap at resolution 17,500, AGC target 5e4, maximum injection time 120 ms with fixed mass at 180 m/z. Charge exclusion was selected for unassigned and 1+ ions. The dynamic exclusion was set to 5 s.

Acquired tandem mass (MS/MS) spectra were searched using Sequest HT in Proteome Discoverer (v1.4), against a custom *Homo sapiens* database (Proteome ID UP000005640) downloaded from UniProt (2025_04_29), containing 20,406 protein entries, including 292 common laboratory contaminants and NCBP1-Halo-His sequence. The following variable modifications were considered – Carbamidomethylation and HNE modifications (+134.073 Da, 152.084 Da, 154.099 Da) on cysteine residues, in addition to Oxidation (M), Acetylation (N-ter). Two missed cleavages were permitted. Peptide mass tolerance was set at 20 ppm on the precursor and 0.6 Da on the fragment ions. Data was filtered at FDR below 1% at PSM level. Trypsin was set as the endoprotease. All identified HNE modified MS/MS spectra were manually inspected and validated.

## Data availability

Proteomics data that support the findings of this study (**Fig. 1D, 2B** and **2C**, **Supplementary Table 3**) have been deposited to the ProteomeXchange Consortium via the PRIDE partner repository with the dataset identifier **PXD:051483**. The RNA-seq raw data from this study used to generate **Fig. 4A, 4E** and **5A**, **Extended Data Fig. 9, Supplementary Fig. 18-21, and Supplementary Table 4-8** have been deposited in the Gene Expression Omnibus (GEO) database under accession code (**GSE:265951**).

---

**Extended Data Fig 1.**
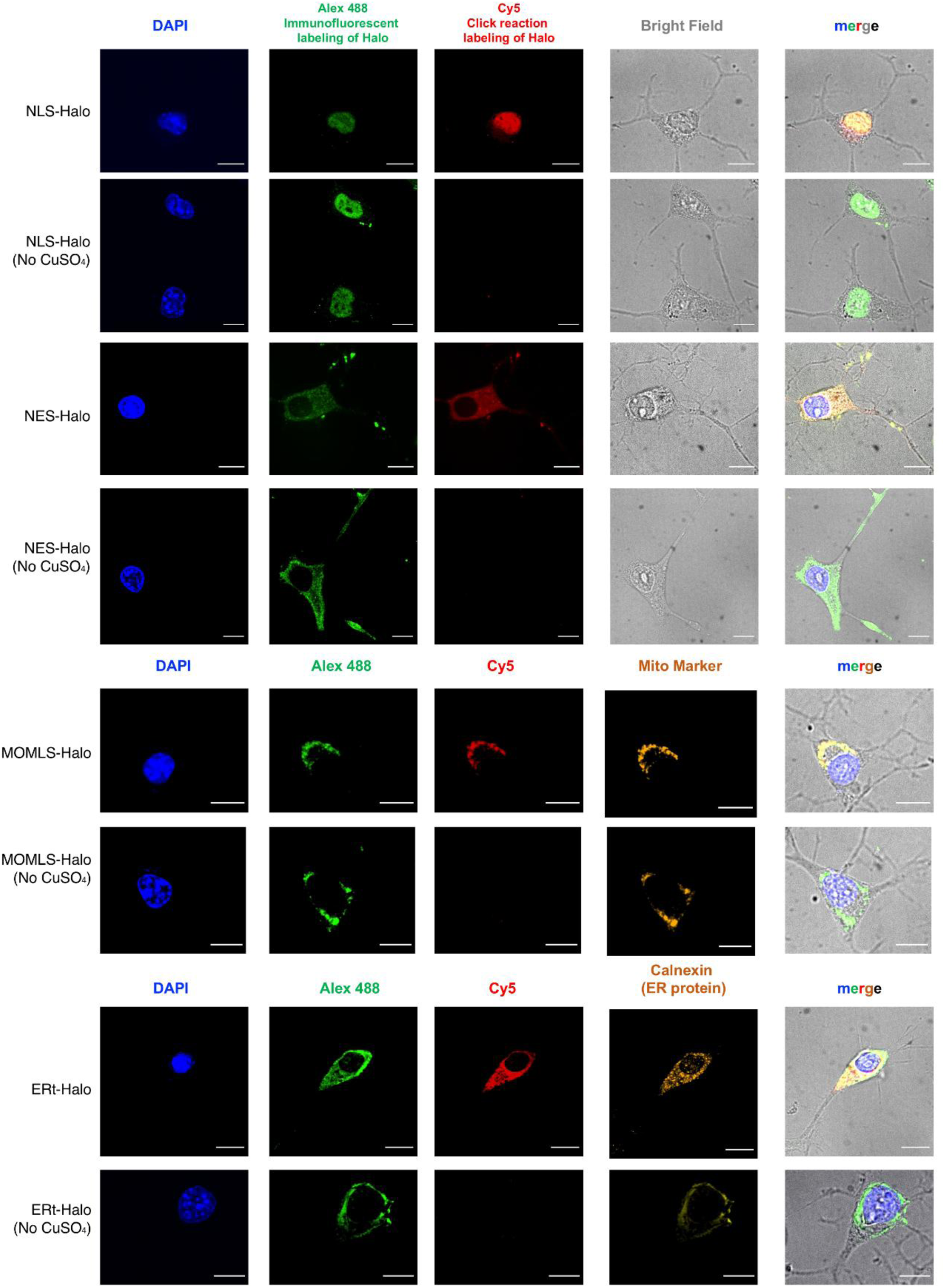
Immunofluorescence imaging validates expected localization of Ht-PreHNE(alkyne) covalently bound to Halo expressed at designated subcellular locales. Differentiated N2A cells ectopically expressing either NLS-Halo, NELS-Halo, MOMLS-Halo, or ERt-Halo were incubated with Ht-PreHNE(alkyne) (10 µM) for 2 h, followed by rinsing, permeabilization and fixing. Localization of Ht-PreHNE(alkyne)-bound Halo was analyzed by Click coupling to Cy5-azide, using anti-Halo antibody and anti-mouse Alex 488 as primary and second antibodies, respectively. DAPI and MitoTracker Red (CaM7512, Thermo Fisher) were used as a nuclear and mitochondria marker, respectively. ER-resident Calnexin (sc-23954, Santa Cruz) was used as an ER marker. Conditions without Cu (hence, no Click coupling) were used as control for background labeling with Cy5-azide. Imaging was performed using a Nikon spinning disk confocal microscope equipped with a 100X objective. Scale bar, 10 µm.

**Extended Data Fig 2.**
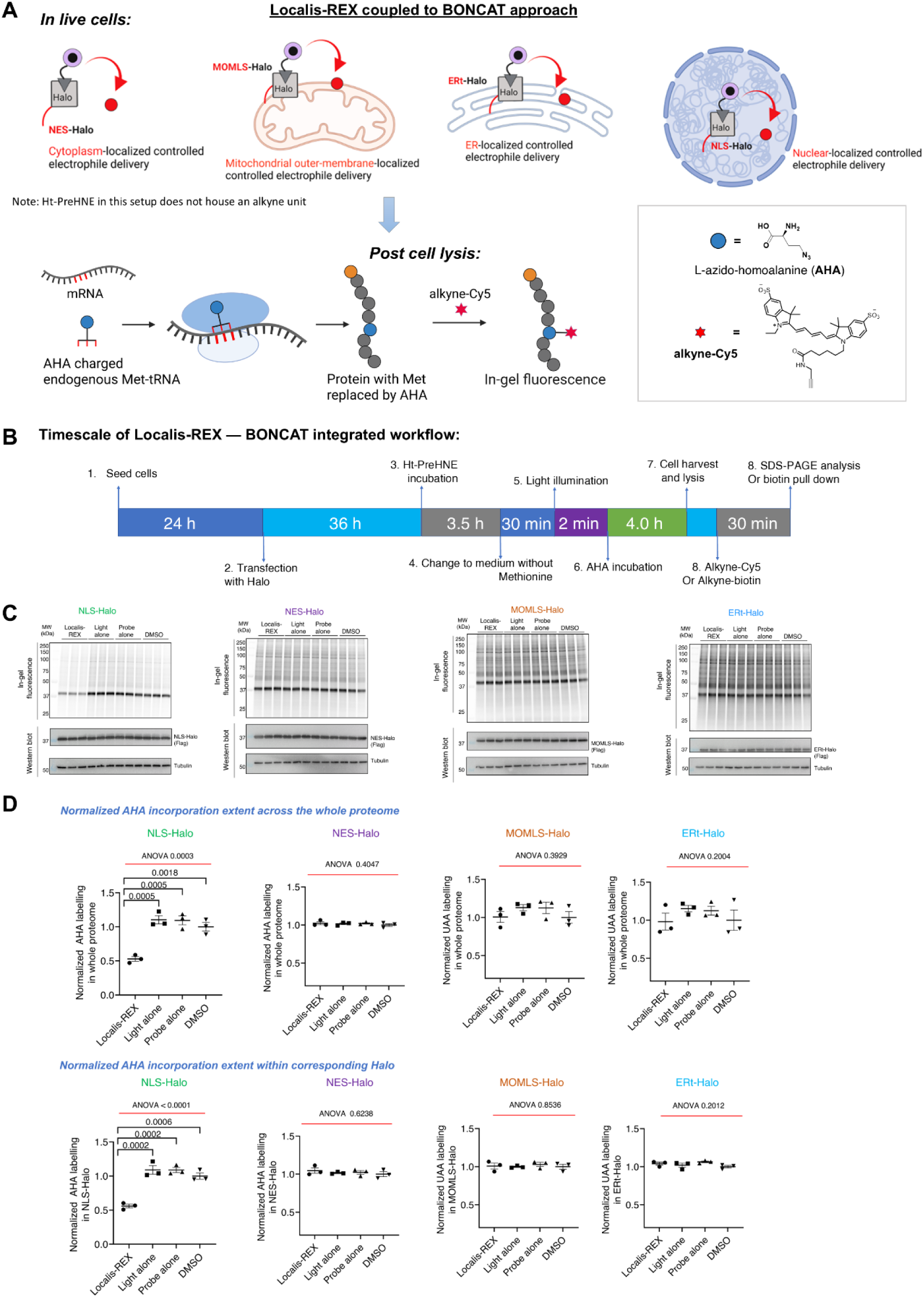
Localis-REX coupled with Click-biotin pulldown and LFQ-based quantitative proteomics, identifies nuclear-specific kinetically-privileged electrophile-sensor proteins. See also Fig. 1D and Supplemental Table S3. **A)** Chemical structures of REX-probes used in the assay. **B)** Representative datasets. Briefly, post Localis-REX execution in differentiated N2A cells expressing NLS-Halo, cells were lysed and subjected to Click-biotin-pulldown (see **Methods**). Both elution and input samples were resolved by SDS-PAGE and stained with colloidal Coomassie to detect proteins. ‘Control’ indicates an identical experiment except that no-alkyne-functionalized version of REX-probe was used. **C)** Results from 3 independent biological replicates stemming from Localis-REX coupled to differential enrichment-based proteomics target-ID involving label-free quantitation (LFQ) (see **Supplemental Table S3**), were subjected to Gaussian fit analysis examining LFQ ratios distribution as indicated. The x-axis indicates log_2_ [alkyne : no-alkyne control] of each hit detected in each replicate, and the y-axis indicates the number of proteins. Proteins of log_2_ [alkyne : no-alkyne control] more than two σ of Gaussian filter were considered as significant hits in each replicate. ‘alkyne’ and ‘no-alkyne control’ indicate the two REX-probes used for differential enrichment proteomics workflow. **D)** Correlation plots of the proteomics datasets: comparisons were performed between indicated independent replicates, with the resultant R^2^ greater than or equal to 0.96 in all cases. Heat map representation of R^2^ was shown on the *left*.

**Extended Data Fig 3.**
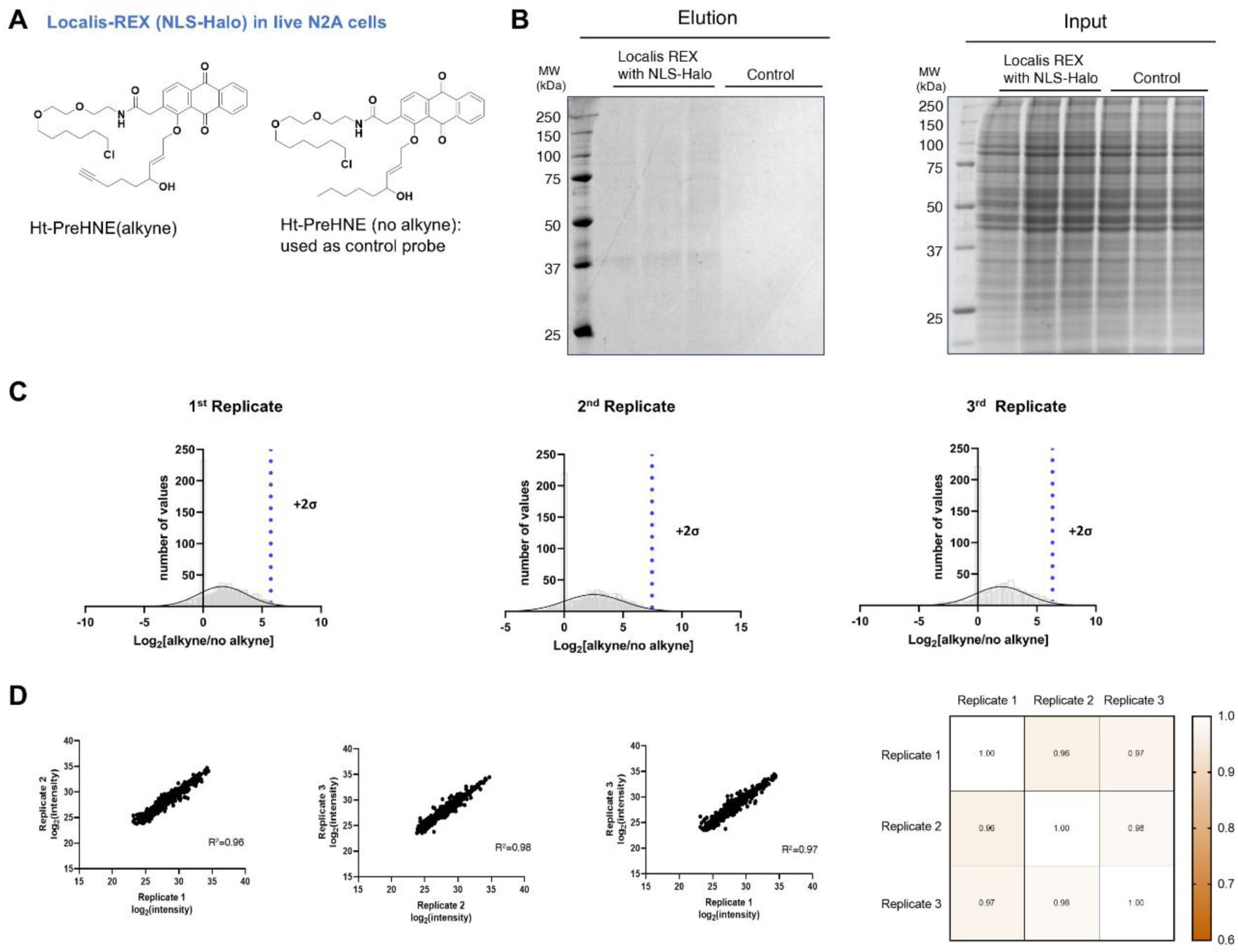
BONCAT experiment coupled with Localis-REX at different subcellular locales, shows reduction of nascent proteome synthesis in response to nuclear targeted electrophile delivery. **A)** and **B)** Schematic diagram and workflow of Localis-REX coupled to BONCAT assay. **C)** Localis-REX was performed similarly to **Supplemental Fig. S2**, except that the cells were incubated in media without Methionine for 30 min prior to light illumination (2 min), triggering localised electrophile generation. The cells were next subjected to media containing L-Azido-homoalanine (AHA, 50 µM, 4 h), to allow AHA incorporation into nascent proteome (see **Methods**). Post cell lysis, Cy5-Click coupling allowed quantification of AHA incorporation, correlating to translation efficiency. **D)** ‘Normalized AHA labelling in whole proteome’ (y-axis for plots in upper panel) corresponds to relative values derived from the in-gel fluorescence signal of Cy5, normalized by that of anti-tubulin; ‘Normalized AHA labelling in corresponding Halo’ (Y-axis for plots in lower panel) corresponds to relative values derived from the in-gel fluorescence signal of Cy5 under corresponding halo band, normalized by that of anti-tubulin; *p* values were calculated with Tukey’s multiple comparisons test. All data present mean±SEM (*n*=3).

**Extended Data Fig 4.**
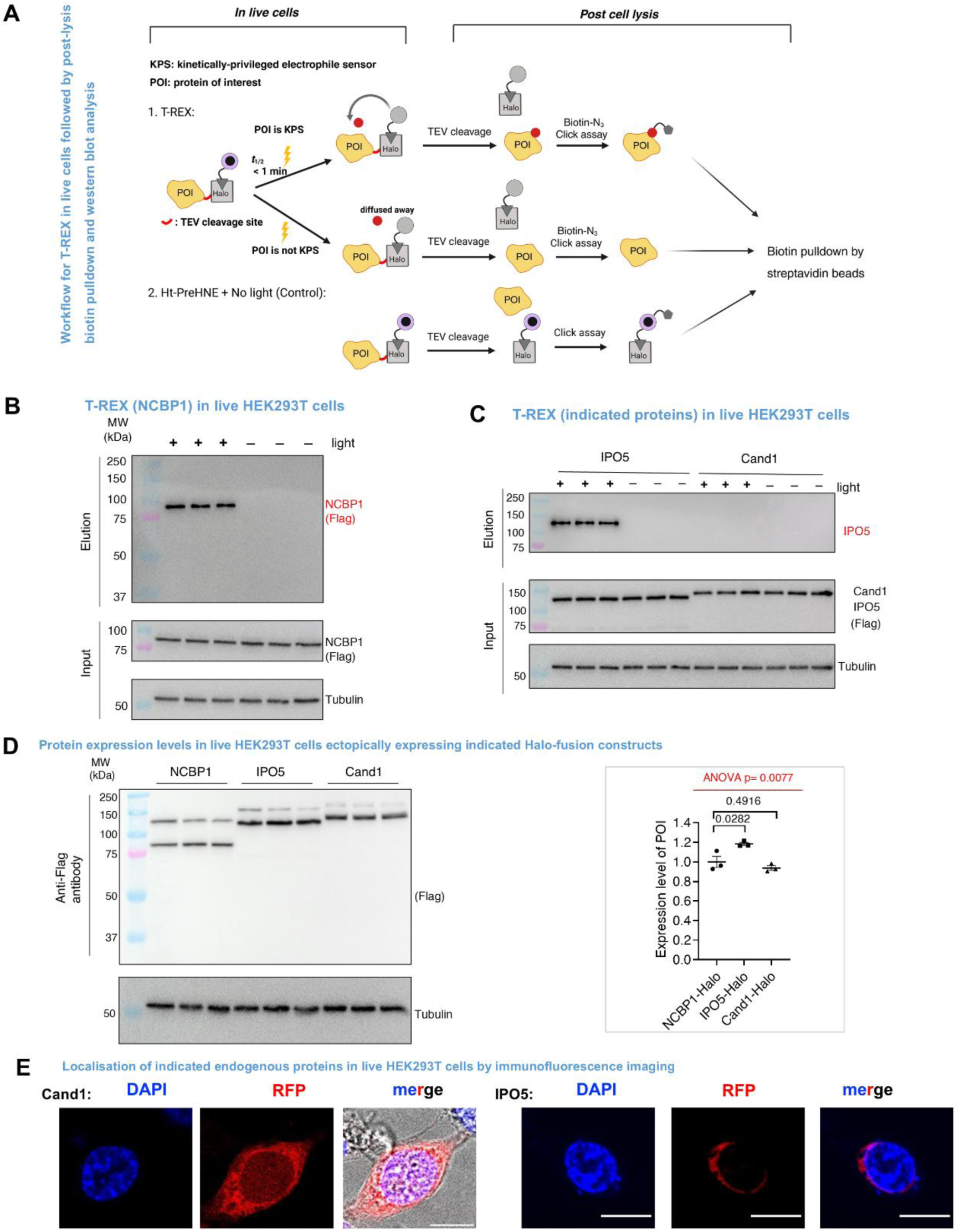
T-REX coupled with Click biotin pulldown shows only NCBP1 and IPO5, but not Cand1, manifest HNE responsivity following localised electrophile delivery. **A)** Schematic of T-REX coupled with Click biotin pulldown to validate electrophile-responsive proteins. **B)** and **C)** T-REX was performed in HEK293T expressing NCBP1-Flag(Tev-site)-Halo, IPO5-Flag(Tev-site)-Halo, or Cand1-Flag(Tev-site)-Halo (**Methods**). Post lysis, all samples (except Input samples), were subjected to TeV-cleavage [separating Halo and flag-tagged protein of interest (POI)]. All samples, including Input samples, were next subjected to Click biotin-azide coupling, and streptavidin-bead enrichment (**Methods**). Both Input and Elution samples were resolved by SDS-PAGE, blotted and analyzed using indicated antibodies. The band from Anti-Flag blot in the Elution represents HNE(alkyne)-modified POI (post TeV cleavage). This band was observed only in light-exposed samples. Conversely, the band in Input represents Ht-PreHNE(alkyne) photocaged probe-bound Halo-POI in no light-exposed controls, or in light-exposed samples, residual photocaged probe-bound Halo-POI, or HNE(alkyne)-modified POI fused to Halo with residual photocaged probe on Halo. **D)** Relative expression of NCBP1-Flag(Tev-site)-Halo, Cand1-Flag(Tev-site)-Halo and IPO5-Flag(Tev-site)-Halo was assessed by anti-Flag blot, normalized by anti-tubulin. Samples were derived from HEK293T cells transfected with the corresponding constructs: POI-Flag(Tev-site)-Halo, followed by TeV protease treatment in cell lysate—performed to replicate the TeV-treatment step deployed in **Fig. 2B-C**. Consistent with Fig. 2B, TeV-cleavage was significantly limited for POI-Flag(TeV-site)-Halo when POI was NCBP1. Quantification is shown in the *Right Panel*. *p* values were calculated using Tukey’s multiple comparison test. All data present mean±SEM (*n*=3). **E)** Immunofluorescence was performed in HEK293T (**Methods**). Primary antibodies: anti-Cand1 (mAb, Santa Cruz, sc137055, 1:100) and anti-IPO5 (pAb, Abcam# ab187175, 1:100). Secondry antibodies: anti-ms IgG Alex Fluor 568 and anti-rabbit IgG Alex Fluor 568 1:2000. DAPI was used for nuclear staining. Images were taken using a Nikon spining-disk confocal microscope equipped with a 100X objective. Scale bar, 10 µm.

**Extended Data Fig 5.**
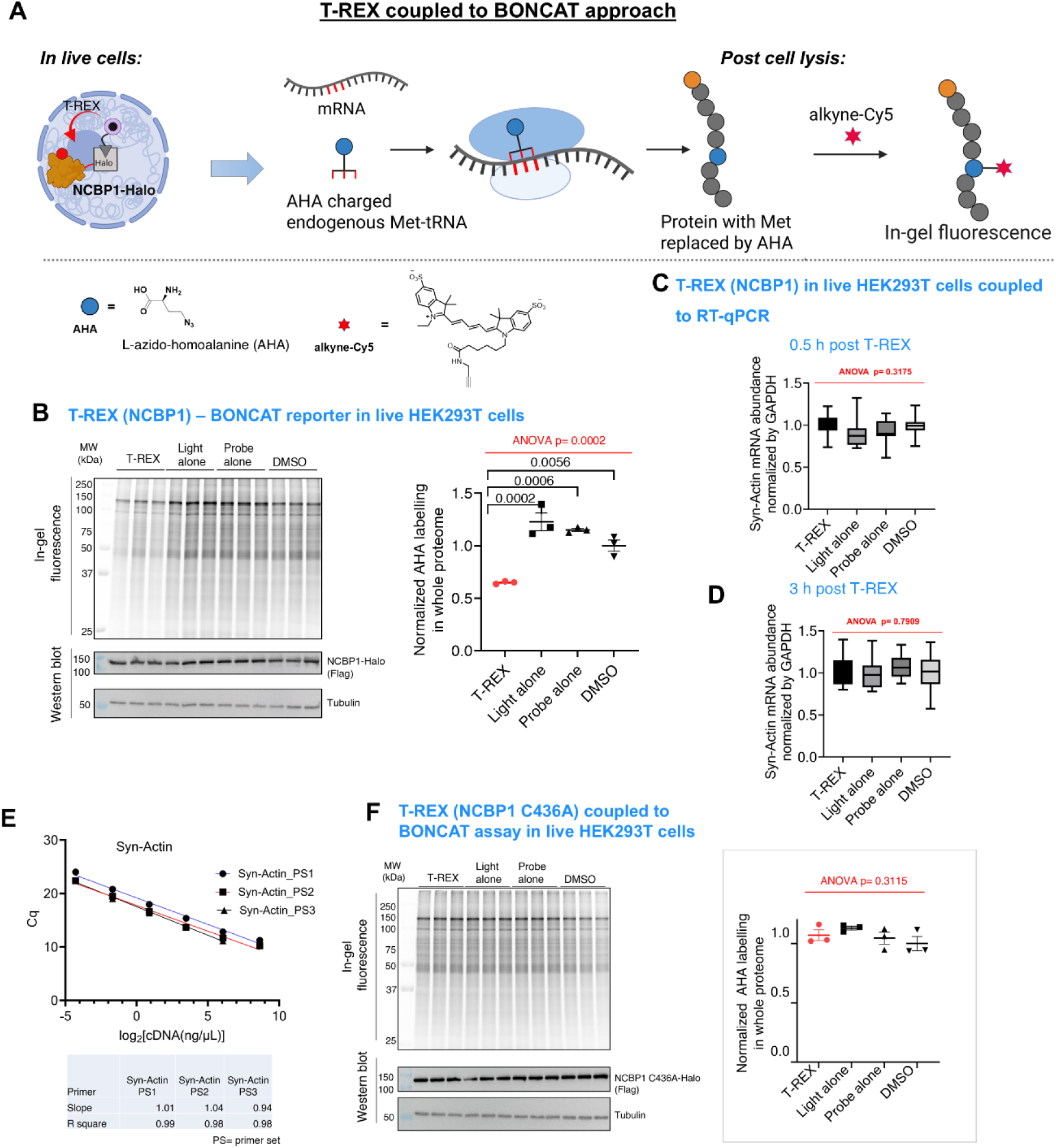
T-REX coupled to BONCAT, shows reduction of nascent proteome synthesis in response to NCBP1-HNEylation, but not NCBP1(C436A)-HNEylation. **A)** Schematic of T-REX coupled with BONCAT, followed by in-gel fluoresccence. **B)** T-REX was performed similarly to **Fig. 2B**, except cells were incubated in media without Methionine for 30 min prior to 2-min light exposure that triggers localised HNE (no-alkyne) generation within the vicinity of NCBP1(wt)-Halo. Post light exposure, cells were subjected to media containing L-Azido-homoalanine (AHA, 50 µM, 4 h) allowing AHA incorporation into nascent proteome (see **Methods**). Post lysis, samples were subjected to Cy5-alkyne Click-coupling reaction. Inset shows quantification. ‘Normalized AHA labelling in whole proteome’ (y-axis) shows relative values derived from the in-gel fluorescence signal of Cy5, normalized by anti-tubulin blot. *p* values were calculated with Tukey’s multiple comparisons test. Data present mean±SEM (*n*=3). C) qRT-PCR analysis (**Methods** and **Supplemental Table S6**) validated that NCBP1-HNEylation does not affect synthetic actin (Actin (K118TAG)) mRNA, compared to controls; ruling out that Syn-Actin suppression (e.g., **Fig. 2D**) is an off-target consequence of the experimental setup deployed. T-REX was performed in HEK293T expressing NCBP1-Halo, Actin(K118TAG), and tRNA synthetase, and RNA isolation post T-REX-assisted NCBP1-specific HNEylation (see **Methods**) was carried out at: **C)** 0.5 h and **D)** 3 h. *P* values were calculated with Tukey’s multiple comparison test. All data present mean±SEM (*n*=12). **E)** Primer efficiency tests (**Methods**) [Data present mean±SEM (n=6; 3 biological replicates, 2 technical replicates each). **F)** T-REX-BONCAT experiments were peformed as in **Extended Data Fig 5A**, except cells expressed NCBP1(C436A)-Halo over NCBP1-wt-Halo. Inset shows quantification. ‘Normalized AHA labelling in whole proteome’ (y-axis) corresponds to relative values derived from Cy5 in-gel fluorescence, normalized to anti-tubulin signal. *p* values were calculated with Tukey’s multiple comparisons test. Data present mean±SEM (*n*=3).

**Extended Data Fig 6.**
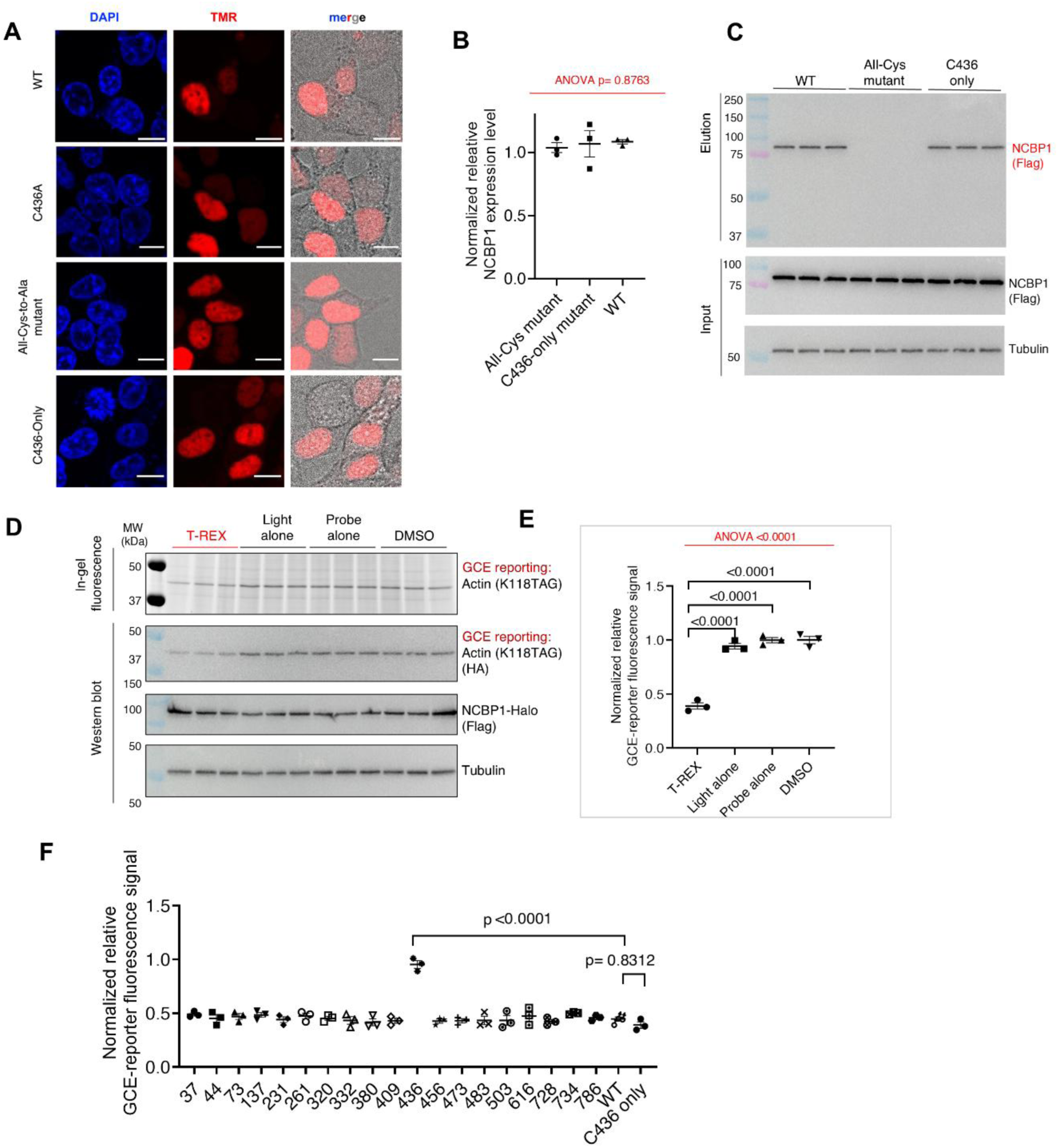
C436 as the sole functional site on NCBP1 that can propagate electrophile signal to protein translation inhibition. **A)** Live imaging shows nuclear localization of NCBP1-wt and mutants. HEK293T expressing NCBP1(wt)-Halo or indicated mutants were subjected to live imaging assay as described in **Fig. 2E**, using cell-permeable Halo-targetable TMR dye. Images were taken using a Nikon spining-disk confocal microscope equipped with a 100X objective. Scale bar, 10 µm. **B)** Validation of protein expression levels of indicated NCBP1 constructs expressed in HEK293T by western blot. p values were calculated with Tukey’s multiple comparisons test. All data present mean±SEM. **C)** T-REX in HEK293T coupled to post-lysis Click biotin pulldown assay was performed as in **Extended Data F4B**, except that cells were transiently transfected with indicated NCBP1(wt or mutant)-Halo constructs. Briefly, signal in elution blot indicates HNEylated NCBP1 (wt or mutant, post TeV-cleavage that separates Halo and the protein of interest) following T-REX. **D)** and **E)** HEK293T co-transfected with NCBP1(C436-only)-Halo and GCE(actin) reporter were subjected to similar experiments as **Fig. 3C** (**Methods**). *p* values were calculated with Tukey’s multiple comparisons test. All data present mean±SEM. **F)** Quantification of relative translation efficency as reported by GCE. The numbers within x-axis show Cys to Ala single-mutation sites. For **B,E,F**, *n*=3 independent expreiments.

**Extended Data Fig 7.**
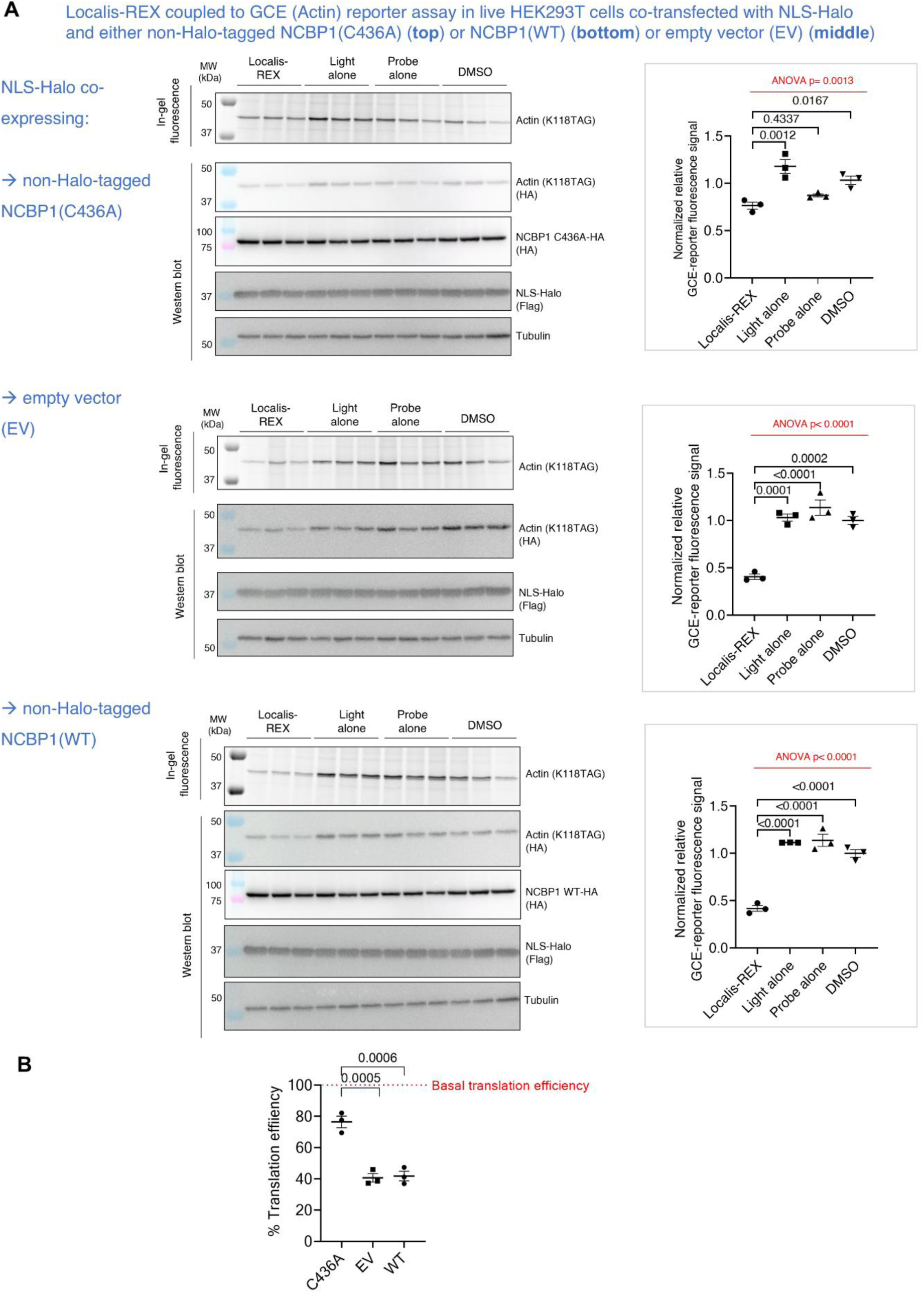
The extent of protein translation suppression induced by Localis-REX-assisted nucleus-targeted HNE release was attenuated in cells overexpressing (non-Halo-tagged) NCBP1(C463A) compared to empty vector or (non-Halo-tagged) NCBP1(wt) overexpressing cells, under otherwise identical conditions. **A)** Top panel: Localis-REX—GCE assay (see **Fig. 1A**, and **Methods**) was performed in HEK293T cells ectopically co-overexpressing NLS-Halo and (no-Halo-tagged) NCBP1(C436A)-HA. The data were analyzed using in-gel fluorescence and western blot analyses. Middle and lower panels: the above experiment was replicated using either Empty vector (EV) or NCBP1(wt)-HA in place of NCBP1(C436A). Insets on right show corresponding quantification in each case. p values were calculated with Tukey’s multiple comparisons test (*n*=3). Data present mean±SEM. **B)** Relative comparison of the 3 datasets in (A) as fold-change suppression of translation efficiency in cells transfected with either NCBP1(C436A), empty vector (EV), or NCBP1 (WT). Values in each case were obtained by GCE-reporter signals measured under T-REX normalized to that of samples under DMSO control. 100% on the y-axis designates the basel translation efficieny. 59 (±5)% translation suppression from basal translation efficiency was measured in EV-transfected cells compared to 24 (±6)% in cells overexpressing NCBP1(C436A). *p* values were calculated with two-tailed, unpaired Student’s *t*-test (*n*=3).

**Extended Data Fig 8.**
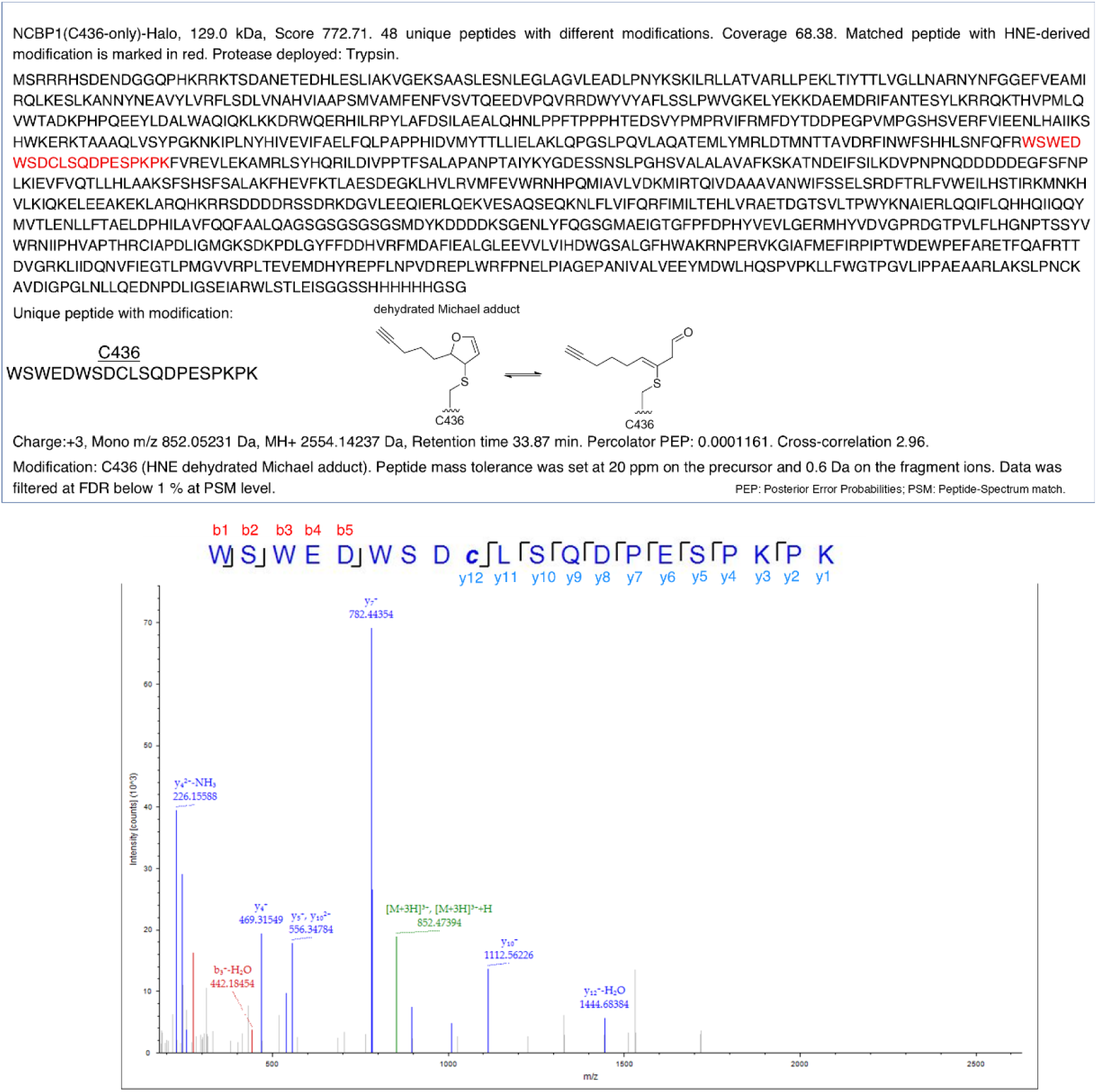
Digest mass spectrometry analysis validated C436 as functional HNEylation site consistent with all other biochemical and cell-based datasets. The experimental workflow and data analysis were described in ‘Mass spectrometry analysis of HNEylation site following T-REX in live HEK293T cells expressing the indicated NCBP1 construct’ (**Methods**).

**Extended Data Fig 9.**
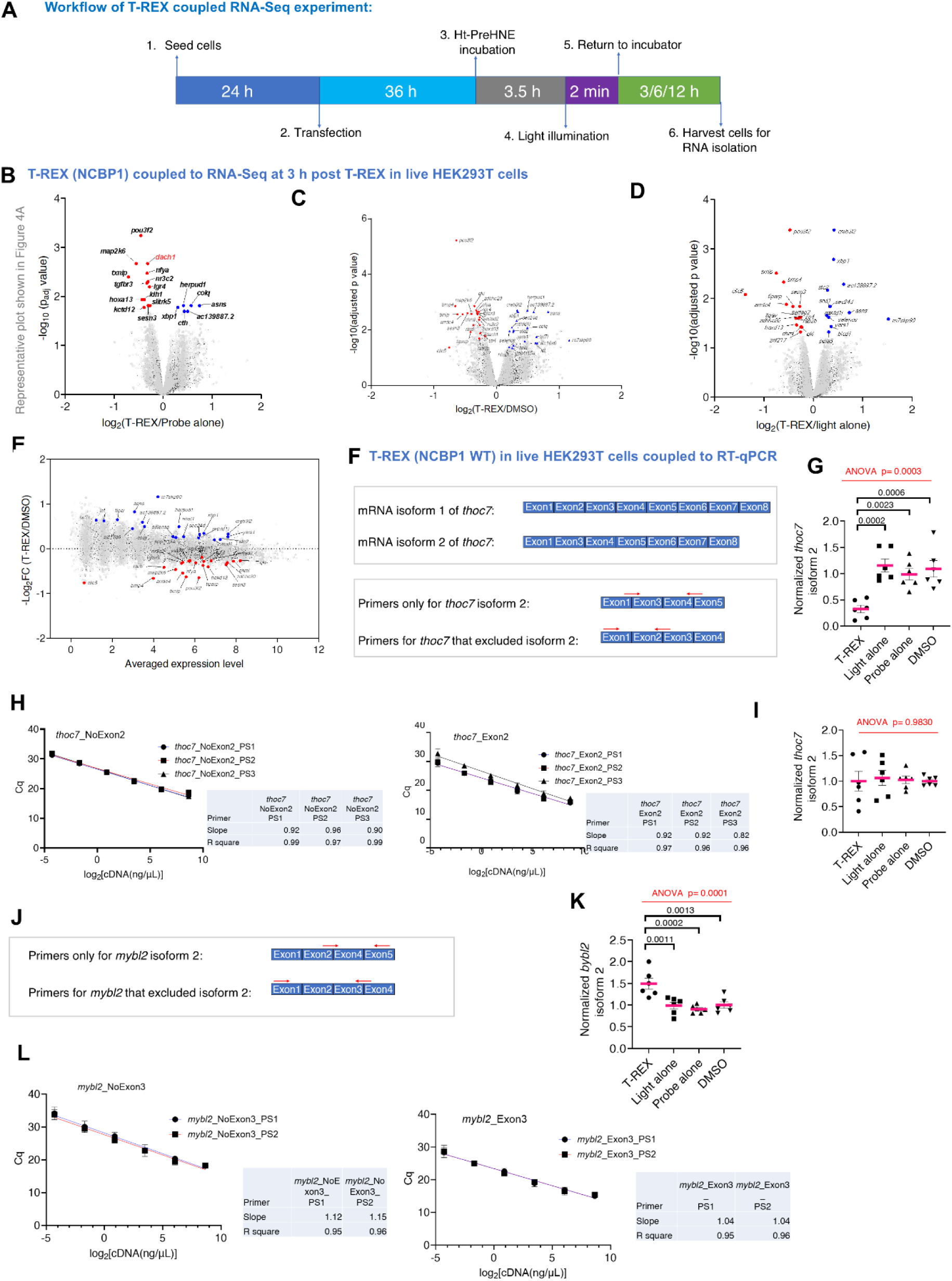
HNEylation of NCBP1 affects mRNA splicing. **A)** Workflow for T-REX-coupled RNA-Seq experiments. **B) C)** and **D)** Volcano and E) MA plots derived from RNA-seq analysis showing statistically-significant differentially-expressed genes (SDEs) for T-REX versus indicated controls (*3 h post T-REX*). For volcano plots (*top 3 plots*): x-axis-log_2_(Fold change (FC) each gene between T-REX and one technical control (probe alone; DMSO; or light alone)); y-axis-adjusted p value [-Log10(padj)]. (top-left plot is identical to Figure 4A). Red dot-genes significantly downregulated by T-REX; blue dots-same but upregulated. Names are shown for arbitrarily-selected SDEs. See **Supplemental Table S4**. In MA scatter plot (*see bottom plot*the x-axis-average expression level of each gene; y-axis-log_2_(FC each gene between T-REX and DMSO). Only hits with p<0.05 for T-REX versus at least two out of 3 technical controls were considered. The names of hits were marked in the MA plot; red dots-downregulated SDEs, blue dots-upregulated SDEs. **F)** Schematic of relevant *THOC7* mRNA isoforms (*top*) and qRT-PCR primer-annealing sites (*bottom*). **G)** qRT-PCR results reporting the relative abundance of isoform 2 normalized by abundance of isoform-1. The plot shows quantification of mRNA of THOC7 isoform-2 across different conditions. *p* values were calculated with Tukey’s multiple comparison test. Data present mean±SEM (n=6). **H)** Primer efficiency test (n=6; 3 biological replicates, 2 technical replicates each). **I)** Similar setup/quantification as **G**, except HEK293T were transfected with NCBP1(C436A)-Halo over NCBP1(wt)-Halo. *p* values were calculated with Tukey’s multiple comparison test. Data present mean ± SEM (n=6). **J)** Schematic of relevant *MYBL2* mRNA isoforms and illustration of qRT-PCR primer-annealing sites. **K)** qRT-PCR results on isoform-2 abundance (generated through skipping of exon 2) normalized to other isoforms. Data present mean±SEM (n=6). **L)** Two primer sets were designed to quantify mRNA of MYBL2 isoform-2 by RT-qPCR analysis Data present mean± SEM (n=12). See **Supplemental Table S6** for primers.

**Extended Data Fig 10.**
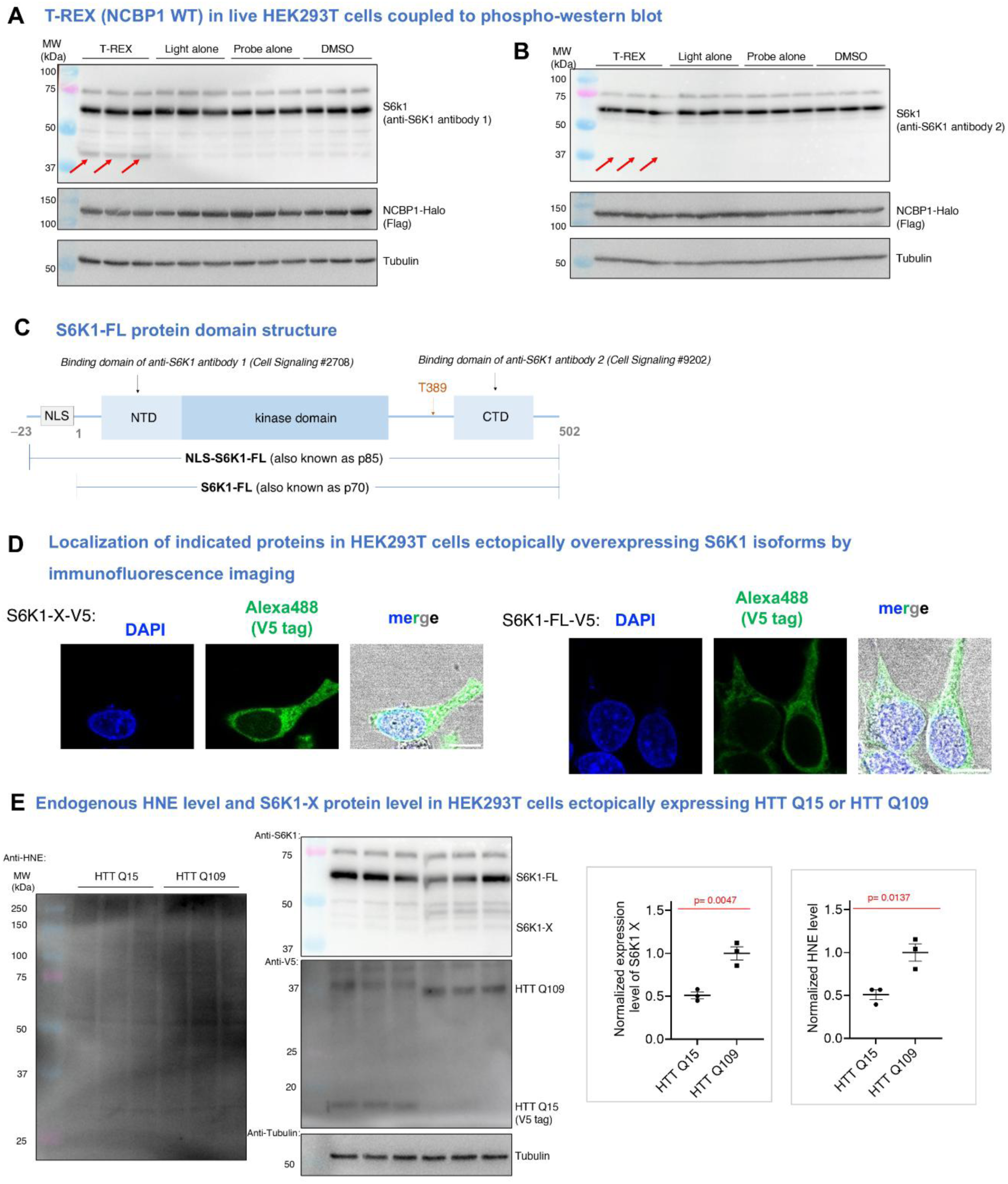
S6K1-X is a suppressor of protein translation. **A)** S6K1 full length and isoform-X are primarily localized to the cytoplasm. HEK293T were transfected with S6K1-X-V5 or S6K1-FL-V5. 36 h later, the cells were fixed and incubated with blocking buffer. Anti-V5 (Invitrgen R96025, 1:200) incubation (2 h) was followed by washing with DPBS three times and anti-ms IgG Alex Fluor 488 (1:2000) secondry antibody treatment. Cells were again rinsed three times with DBPS followed by treatment with DAPI (2 µg/mL in DPBS, 5 min), and final rinsing with DPBS. Images were taken using a Nikon spining-disk confocal microscope equipped with a 100X objective. Scale bar, 10 µm. **B)** and **C)** Expression of S6K1 isoform-X, following NCBP1-HNEylation, was examined using two different S6K1 antibodies. See also Fig. 5E. 2 h post T-REX in HEK293T expressing NCBP1-Halo, cells were lysed and analyzed by western blot using indicated antibodies. **D)** Domain structure of S6K1-FL highlighting double (or multiple) bands typically observed by western blot, corresponding to two key forms: S6K1-FL (p70) and NLS-S6K1-FL (p85). S6K1-FL lacks the nuclear localization sequence (NLS) encoding 23 amino acids at the N-terminal end. **E)** S6K1-X expression was upregulated in a cell-based Huntingtin disease model. HEK293T ectopically expressing either HTT-Q15 (non-disease control) or HTT-Q109 (disease-inducing construct) were analysed by western blot using the indicated antibodies, reporting the extent of proteome HNEylation and endogenous S6K1 and S6K1-X expression levels. Quantification plots are shown on the right. *p* values were calculated using unpaired, two-tailed Student’s *t*-test. All data present mean± SEM (n=3).

