## Supplementary information for "NCBP1 stress signaling drives alternative S6K1 splicing inhibiting translation"

### Table of Content

Reagents………………………………………………………………………….. p. 1

Supplementary Tables 1-8………………………………………………. p.2- 21

Supplementary Figures 1-36……………………………………………..p. 22-54

Supplemental References…………………………………………………p. 55

**Reagents**.

HaloTag-targetable photocaged precursor to Hydroxynonenal (Ht-PreHNE (no alkyne)) and Ht-PreHNE (alkyne) were synthesized as described previously[1]. Tetrazine-Cy5 (CLK-015-05) was from Jena Bioscience, Bicylco[6.1.0] nonyne-Lysine (BCNK) (sc-8016) was from Sichem, Sulfo-Cy5-azide (B3330), CuTBTA (21050), L-Azidohmoalanine (3799), Cyanine 5 alkyne (B30B0), and Biotin alkyne (C37B0) were from Lumiprobe. Biotin-dEG-azide (PEG4340.0100) was from Iris. Pierce high-capacity streptavidin agarose beads (20361) were from Thermo Fisher. Anti-Flag M2 affinity gel (A2220), Bovine Serum Albumin (BSA) power, and retinoic acid (R2625) were from Sigma-Aldrich. Dithiothreitol (DTT), and tris(2-carboxyethyl)phosphine (TCEP)-HCl were from Goldbio Biotechnology. Phusion HotStart II polymerase was from Thermo Scientific. All restriction enzymes were from NEB. The plasmid for recombinant expression of TeV protease (Prk793, Addgene 8827) was from Addgene. N2A and HEK293T cells were from the American Type Culture Collection (ATCC). Protease inhibitor cocktail cOmpleteTM EDTA-free (11873580001) was from Roche. Minimal Essential Media (41090028), Dulbecco's Modified Eagle Media (12491015), Opti-MEM (51985026), Dulbecco's PBS (DPBS, 14190169), TrypLE Express Enzyme (12605028), 100X pyruvate (100 mM, 11360039), 100X non-essential amino acids (11140035), 100X penicillin-streptomycin (15140122), NP-40 (85124) were from Life Technologies. Fetal bovine serum (FBS) was from Sigma Aldrich (F2442). Light exposure was performed using Spectroline and Camag ultraviolet lamps (ENF-240C [Spectroline] and lamp 4 [Camag]. The lamps were positioned above a confluent monolayer of cells with power ~5mW/cm$^2$ on samples. TransIT-2020 (MIR 5406) was from Mirus. PEI (23966-1) was from Polysciences. Actinomycin D (A9415) was from Sigma-Aldrich. All sterile cell culture plasticware was from TPP or CellTreat. TRIzol reagent (15596018) was fromAmbion. Passive lysis buffer for luciferase reporter assay was from Promega. Direct-zol RNA Kits was from Zymo (R2050) Superscript III (18080085) and RNaseOUT (10777019) enzymes for qPCR experiment were from Thermo Fisher. SYBR Green Master Mix reagent for RT-qPCR was from Bio-Rad. Sources of antibodies are listed in **Supplementary Table 4**.

### Supplementary Tables

**Supplementary Table 1. Primers used for cloning.**

| plasmid | Primer | Sequence (5'→3') |
|---|---|---|
| pCS2+8 -NLS-Halo | Forward primer1 | ATGCCAAAAAAGAAGCGCAAGGTATCGGGTCCTAAGAAGAAGCGTAAAGTGTCAGG |
| | Reverse primer1 | GGTTGATTATCGATAAGCTTGATATCGAATTCTTAGCCGGAAATCTCGAGCGTC |
| | Forward primer2 | TTTCTTCCATTTCAGGTGTCGTGAGGATCCGCCACCATGCCAAAAAAGAAGCGCAAGG |
| | Reverse primer2 | GGTTGATTATCGATAAGCTTGATATCGAATTCT |
| pCS2+8 -NES-Halo | Forward primer1 | GAGCTTGCCCTAAAGCTAGCAGGCCTCGATCTATCTGGAATGGCAGAAATCGGTACTGGC |
| | Forward primer2 | TCGCACTCAAACTCGCCGGGCTAGACCTGTCAGGGGAAGAGCTTGCCCTAAAGCTAGCAG |
| | Forward primer3 | TCCATTTCAGGTGTCGTGAGGATCCGCCACCATGGAGGAACTCGCACTCAAACTCGCCG |
| | Forward primer4 | TTCTTCCATTTCAGGTGTCGTGAGGATC |
| | Reverse primer1 | GGTTGATTATCGATAAGCTTGATATCGAATTCTTAGCCGGAAATCTCGAGCGTC |
| | Reverse primer2 | GGTTGATTATCGATAAGCTTGATATCGAATTCT |
| pCS2+8 -MOMLS-Halo | Forward primer1 | TTTCTTCCATTTCAGGTGTCGTGAGGATCCGCCACCATGAAGAGCTTCATTACAAGG |
| | Reverse primer1 | GGTTGATTATCGATAAGCTTGATATCGAATTCTTAGCCGGAAATCTCGAGCGTC |
| | Forward primer2 | TTTCTTCCATTTCAGGTGTCGTGAGGATCCGCCACCATGCCAAAAAAGAAGCGCAAGG |
| | Reverse primer2 | GGTTGATTATCGATAAGCTTGATATCGAATTCT |
| pCS2+8 -ERt-Halo | Forward primer1 | AAGCTGATCATCGATCAGAACGTTTTTATC |
| | Reverse primer1 | GGTTGATTATCGATAAGCTTGATATCGAATTCTTATCACTCCTGCGCGATGCT |
| | Forward primer2 | TTTCTTCCATTTCAGGTGTCGTGAGGATCCGCCACCATGCCAAAAAAGAAGCGCAAGG |
| | Reverse primer2 | GGTTGATTATCGATAAGCTTGATATCGAATTCT |
| | Forward primer1 | TAATTAAAGGCCGGCCAGCGATCGCCGGACCCACCATGTCGCGGCGGCGG |

| | | |
|---|---|---|
| pCS2+8-NCBP1-Flag-TEV-Halo | Reverse primer1 | CCTGACTTGTCGTCATCGTCTTTATAATCCATGGCCTGCAGGGCACAGAA |
| | Forward primer2 | ACTTGTTCTTTTTGCAGGATCCACTAGTGGCGCGCCATTAATTAAAGGCCGGCCAGCGA |
| | Reverse primer2 | TCCCTGAGCCCTGGAAATACAAGTTTTCCCCTGACTTGTCGTCATCGTCTTTATAATCC |
| pCS2+8-Cand1-Flag-TEV-Halo | Forward primer1 | TAATTAAAGGCCGGCCAGCGATCGCCGGACCCACCATGGCGAGCGCCTCGTAC |
| | Reverse primer1 | CTGACTTGTCGTCATCGTCTTTATAATCCATACTAGTGTCCATTGATTCCAAGTTAGTAG |
| | Forward primer2 | Same with Forward primer2 of pCS2+8-NCBP1-Flag-TEV-Halo |
| | Reverse primer2 | Same with Reverse primer2 of pCS2+8-NCBP1-Flag-TEV-Halo |
| pCS2+8-IPO5-Flag-TEV-Halo | Forward primer1 | TTAAAGGCCGGCCAGCGATCGCCGGACCCACCATGGCGGCGG |
| | Reverse primer1 | CCTGATCCTGACCCGCTTCCGGATCCCGCAGAGTTCAGGAGCTCCT |
| | Forward primer2 | Same with Forward primer2 of pCS2+8-NCBP1-Flag-TEV-Halo |
| | Reverse primer2 | Same with Reverse primer2 of pCS2+8-NCBP1-Flag-TEV-Halo |
| pCS2+8-NCBP1 C380A-Flag-TEV-Halo | Forward primer | AGAGCCAGGTTGAAGTTTGGCCAGTTCAATGAGGAGTGTT |
| | Reverse primer | AACACTCCTCATTGAACTGGCCAAACTTCAACCTGGCTCT |
| pCS2+8-NCBP1 C332A-Flag-TEV-Halo | Forward primer | AACTGTGCAGCAGCAGTCTTCCTTTCCTTCCAGTGGG |
| | Reverse primer | CCCACTGGAAGGAAAGGAAGACTGCTGCTGCACAGTT |
| pCS2+8-NCBP1 C456A-Flag-TEV-Halo | Forward primer | GGTAAGACAACCTCATAGCTTTTTCTAGAACTTCTCTTACAAACTTCGG |
| | Reverse primer | CCGAAGTTTGTAAGAGAAGTTCTAGAAAAAGCTATGAGGTTGTCTTACC |
| pCS2+8-NCBP1 C616A-Flag-TEV-Halo | Forward primer | ATTTGCTACGGCAGCAGCATCAACTATTTGTGTACGAATCATCTTATCC |
| | Reverse primer | GGATAAGATGATTCGTACACAAATAGTTGATGCTGCTGCCGTAGCAAAT |
| pCS2+8-NCBP1 C728A-Flag-TEV-Halo | Forward primer | GTCCCATCAGTTTCGGCTCGTACTAGGTGCTCGGTC |
| | Reverse primer | GACCGAGCACCTAGTACGAGCCGAAACTGATGGGAC |
| pCS2+8-NCBP1 C786A-Flag-TEV-Halo | Forward primer | CGGCCTGCAGGGCAGCGAACTGCTGGAACA |
| | Reverse primer | TGTTCCAGCAGTTCGCTGCCCTGCAGGCCG |
| | Forward primer | AGGTTGCTCTCCAAAGAGGCGGCACTCTTTTCTCCTAC |

| | | |
|---|---|---|
| pCS2+8-NCBP1 C44A-Flag-TEV-Halo | Reverse primer | GTAGGAGAAAAGAGTGCCGCCTCTTTGGAGAGCAACCT |
| pCS2+8-NCBP1 C232A-Flag-TEV-Halo | Forward primer | ATCTGGGCCCACAGGGCATCTAAATACTCTTCTTGTGGATGT |
| | Reverse primer | ACATCCACAAGAAGAGTATTTAGATGCCCTGTGGGCCCAGAT |
| pCS2+8-NCBP1 C261A-Flag-TEV-Halo | Forward primer | TGTGCTGCAGTGCTTCAGCCAGGATGCTGTCAAAGG |
| | Reverse primer | CCTTTGACAGCATCCTGGCTGAAGCACTGCAGCACA |
| pCS2+8-NCBP1 C320A-Flag-TEV-Halo | Forward primer | CTTCCAGTGGGACTTAATGATGGCGTGAAGATTCTCTTCTATTACA |
| | Reverse primer | TGTAATAGAAGAGAATCTTCACGCCATCATTAAGTCCCACTGGAAG |
| pCS2+8-NCBP1 C483A-Flag-TEV-Halo | Forward primer | ATCTCCATACTTGTAAATGGCGGTTGGGTTTGCAGGACAC |
| | Reverse primer | GTGTCCTGCAAACCCAACCGCCATTTACAAGTATGGAGAT |
| pCS2+8-NCBP1 C503A-Flag-TEV-Halo | Forward primer | TTTTAAAGGCAACAGCTAAAGCGAGGGCAACAGAATGTCCAG |
| | Reverse primer | CTGGACATTCTGTTGCCCTCGCTTTAGCTGTTGCCTTTAAAA |
| pCS2+8-NCBP1 C36A-Flag-TEV-Halo | Forward primer | GCACTCTTTTCTCCTACTTTAGCTATTAAAGATTCCAAATGATCTTCAGTTTCAT |
| | Reverse primer | ATGAAACTGAAGATCATTTGGAATCTTTAATAGCTAAAGTAGGAGAAAAGAGTGC |
| pCS2+8-NCBP1 C73A-Flag-TEV-Halo | Forward primer | AGGCGTGCAACTGTAGCAAGAAGCCTTAAGATCTTGCTCT |
| | Reverse primer | AGAGCAAGATCTTAAGGCTTCTTGCTACAGTTGCACGCCT |
| pCS2+8-NCBP1 C137A-Flag-TEV-Halo | Forward primer | GCGGCAATCACATGAGCATTCACAAGATCAGATAAAAAACGGACCA |
| | Reverse primer | TGGTCCGTTTTTTATCTGATCTTGTGAATGCTCATGTGATTGCCGC |
| pCS2+8-NCBP1 C409A-Flag-TEV-Halo | Forward primer | ACCAATTAATAAACCTGTCTACAGCGGTAGTGTTCATTGTGTCCAAAC |
| | Reverse primer | GTTTGGACACAATGAACACTACCGCTGTAGACAGGTTTATTAATTGGT |
| pCS2+8-NCBP1 C436A-Flag-TEV-Halo | Forward primer | CAGGATCTTGACTAAGAGCATCTGACCAATCTTCCCAGCT |
| | Reverse primer | AGCTGGGAAGATTGGTCAGATGCTCTTAGTCAAGATCCTG |
| | Forward primer | CTGCTGCAGCCTCTCTATAGCGTTCTTATACCATGGTGTT |

| | | |
|---|---|---|
| pCS2+8-NCBP1 C743A-Flag-TEV-Halo | Reverse primer | AACACCATGGTATAAGAACGCTATAGAGAGGCTGCAGCAG |
| pCS2+8-NCBP1 C477A-Flag-TEV-Halo | Forward primer | AGGTTGGGTTTGCAGGAGCCAGAGCTGAGAAGGTAG |
| | Reverse primer | CTACCTTCTCAGCTCTGGCTCCTGCAAACCCAACCT |
| psGG_10-11-DACH1-3UTR and psGG_10-11No intron-DACH1-3UTR | Forward primer1 | CTGTTGGTAAAGCCACCATGATAGTTCAAAAGAGGCTAAA |
| | Reverse primer1 | TTTAGCCTCTTTTGAACTATCATGGTGGCTTTACCAACAG |
| | Forward primer2 | TTGGCATCTTCCATTTGTATTGTCCTTTCAGCATCTGTTC |
| | Reverse primer2 | GAACAGATGCTGAAAGGACAATACAAATGGAAGATGCCAA |
| | 3'UTR_Forward primer1 | TCAACGTCTAAGGCCGCGACTCTAGAATCTTTCCTGTTGAAGAAATCCATGTTATAGA |
| | 3'UTR_Reverse primer1 | TCTGCTCGAAGCGGCCGGCCCTCGAGGAGATTTCAGACAGTTTTATTAGGAGATTGTTG |
| | 3'UTR_Forward primer2 | GGCATGGATAGACACCCTGCTGCTTGCGCCAGCGCCAGGATCAACGTCTAAGGCCGCGA |
| | 3'UTR_Reverse primer2 | TAGTTGTGGTTTGTCCAAACTCATCAATGTATCTTATCATGTCTGCTCGAAGCGGCCG |
| pCS2+8-NCBP2-HA | Forward primer1 | ATGTCGGGTGGCCTCCTG |
| | Forward primer2 | TGACACTATAGAATACAAGCTACTTGTTCTTTTTGCCACCATGTCGGGTGGCCTCCTG |
| | Forward primer3 | CGTCGGAGCAAGCTTGATTTAGGTGACACTATAGAATACAAGCTACTTGTTCTTTTTG |
| | Reverse primer1 | CTGGTTCTGTGCCAGTTTTCCATAG |
| | Reverse primer2 | AGGCTCGAGAGGCCTTGAATTCGATTATTACTGGTTCTGTGCCAGTTTTCCATAG |
| | Reverse primer3 | GTCTGGATCTACGTAATACGACTCACTATAGTTCTAGAGGCTCGAGAGGCCTTGAATT |
| | Forward primer1 | GGCGTTCCATTGACGTAAATGGG |
| | Reverse primer1 | CATGCTCCCTGATCCTGACCCGCTTCCGGATCCCTGGTTCTGTGCCAGTTTTCCATAG |

| | | |
|---|---|---|
| pCS2+8-NCBP2-Flag-TEV-Halo | | |
| | Reverse primer2 | TTTTCCCCTGACTTGTCGTCATCGTCTTTATAATCCATGCTCCCTGATCCTGACCC |
| pCS2+8-S6K1-FL-(V5) | Forward primer1 | CCTATCCGTACGACGTACCAGACTACGCAGGATCCGGAAGCATGAGGCGACGAAGGAGGC |
| | Forward primer2 | ATTAATTAAAGGCCGGCCAGCGATCGCCGGACCCACCTATCCGTACGACGTACCAGACT |
| | Forward primer3 | ACTTGTTCTTTTTGCAGGATCCACTAGTGGCGCGCCATTAATTAAAGGCCGGCCAGCG |
| | Reverse primer1 | TAGATTCATACGCAGGTGCTCTGG |
| | Reverse primer2 | TCTAGAGGCTCGAGAGGCCTTGAATTCGATTATTATAGATTCATACGCAGGTGCTCTGG |
| | Reverse primer3 | CTTATCATGTCTGGATCTACGTAATACGACTCACTATAGTTCTAGAGGCTCGAGAGGCC |
| pCS2+8-S6K1-X-(V5) | Forward primer1 | Same as Forward primer1 of pCS2+8-S6K1-FL-(V5) |
| | Forward primer2 | Same as Forward primer2 of pCS2+8-S6K1-FL-(V5) |
| | Forward primer3 | Same as Forward primer3 of pCS2+8-S6K1-FL-(V5) |
| | Reverse primer1 | GATTGACAGGTAGAACACCATGTGA |
| | Reverse primer2 | CTAGAGGCTCGAGAGGCCTTGAATTCGATTATTAGATTGACAGGTAGAACACCATGTGA |
| | Reverse primer3 | TTATCATGTCTGGATCTACGTAATACGACTCACTATAGTTCTAGAGGCTCGAGAGGCCT |
| pCS2+8-NCBP1 C436A-HA | Forward primer1 | TAATTAAAGGCCGGCCAGCGATCGCCGGACCCACCATGTCGCGGCGGCGG |
| | Reverse primer1 | TATGCGTAGTCTGGTACGTCGTACGGATAGCTTCCGGATCCGGCCTGCAGGGCACAGAA |
| | Forward primer2 | ACTTGTTCTTTTTGCAGGATCCACTAGTGGCGCGCCATTAATTAAAGGCCGGCCAGCGA |
| | Reverse primer2 | TCTAGAGGCTCGAGAGGCCTTGAATTCGATTATTATGCGTAGTCTGGTACGTCGTACG |

| | | |
|---|---|---|
| pCS2+8-HTT Q15-HA and pCS2+8-HTT Q109-HA | Forward primer1 | AAGGCCGGCCAGCGATCGCCGGACCCACCATGGCGACCCTGGAAAAGCT |
| | Reverse primer1 | CGTAGAATCGAGACCGAGGAGAGGGTTAGGGATAGGCTTACCTCGGTGCAGCGGCTCC |
| | Forward primer2 | GTTCTTTTTGCAGGATCCACTAGTGGCGCGCCATTAATTAAAGGCCGGCCAGCGA |
| | Reverse primer2 | TCTAGAGGCTCGAGAGGCCTTGAATTCGATTATTACGTAGAATCGAGACCGAGGAGAG |
| pCS2+8-XPO1-Flag-TEV-Halo | Forward primer1 | AAAGGCCGGCCAGCGATCGCCGGACCCACCATGCCAGCAATTATGACAATGTTAGCA |
| | Reverse primer1 | CCCTGATCCTGACCCGCTTCCGGATCCATCACACATTTCTTCTGGAATCTCATGTG |
| | Forward primer2 | GCAGGATCCACTAGTGGCGCGCCATTAATTAAAGGCCGGCCAGCGAT |
| | Reverse primer2 | CCTGACTTGTCGTCATCGTCTTTATAATCCATGCTCCCTGATCCTGACCCGCTTC |
| pCS2+8-RAN-Flag-TEV-Halo | Forward primer1 | AAAGGCCGGCCAGCGATCGCCGGACCCACCATGGCTGCGCAGGGAGAG |
| | Reverse primer1 | CCCTGATCCTGACCCGCTTCCGGATCCCAGGTCATCATCCTCATCCGGG |
| | Forward primer2 | GCAGGATCCACTAGTGGCGCGCCATTAATTAAAGGCCGGCCAGCGAT |
| | Reverse primer2 | CCTGACTTGTCGTCATCGTCTTTATAATCCATGCTCCCTGATCCTGACCCGCTTC |
| pPB-MmPylRS-AF-4xPylT$_{CAC}$ | PT Overlap R | GTCCGTTCGATCTACATGATCAGGTTTC |
| | PT Overlap CAC Val F | ATCATGTAGATCGAACGGACTCACAATCCGTTCAGCCG |
| | PT1 FW | AATAAGGTCTCAGCATCTAGGGCAGGAAGAGGGCCTATTTC |
| | PT1 RV | AATAAGGTCTCACCATTATGGGCAGGAAGAGGGCCTATTTC |
| | PT2 FW | AATAAGGTCTCACCATTATGGGCAGGAAGAGGGCCTATTTC |
| | PT2 RV | AATAAGGTCTCAACTAGAGAAAAACCGCACTTGTCCGGAAA |
| | PT3 FW | AATAAGGTCTCATAGTTGCGGGCAGGAAGAGGGCCTATTTC |
| | PT3 RV | AATAAGGTCTCAGATCCATAAAAACCGCACTTGTCCGGAAA |
| | PT4 FW | AATAAGGTCTCAGATCATGGGGCAGGAAGAGGGCCTATTTC |
| | PT4 RV | AATAAGGTCTCAATTGATTAAAAACCGCACTTGTCCGGAAA |

**Supplementary Table 2. Localization sequences used for HaloTag locale in Localis-REX.**

| | |
|---|---|
| Flag-NLS (at N terminal of Halo) | GATTACAAGGATGACGACGATAAGATGGAGGAACTCGCACTCAAACTCGCCGGGCTAGACCTGTCAGGGGAAGAGCTTGCCCTAAAGCTAGCAGGCCTCGATCTATCTGGA |
| Flag-NES (at N terminal of Halo) | GATTACAAGGATGACGACGATAAGATGGAGGAACTCGCACTCAAACTCGCCGGGCTAGACCTGTCAGGGGAAGAGCTTGCCCTAAAGCTAGCAGGCCTCGATCTATCTGGA |
| MOMLS-Flag (at N terminal of Halo) | ATGAAGAGCTTCATTACAAGGAACAAGACAGCCATTTTGGCAACCGTTGCTGCTACAGGTACTGCCATCGGTGCCTACTATTATTACAACCAATTGCAACAGGATCCACCGGTCGCCACCGATTACAAGGATGACGACGATAAGATG |
| ERt-V5 tag-Flag (at C terminal of Halo) | GATTATAAAGATGATGATGATAAAGGTAAGCCTATCCCTAACCCTCTCCTCGGTCTCGATTCTACGAGATCTCTCCTGGGTGGCGCCCTGGCGAACTTGTTTGTGATAGTTGGGTTTGCAGCCTTTGCTTACACGGTCAAGTACGTGCTGAGGAGCATCGCGCAGGAG |

**Supplementary Table 3. LFQ-MS hits from Localis-REX leveraging nuclear-targeted Halo.**

Please see the enclosed excel file labeled '**Supplemental Table S3 (Localis-REX_proteomics-hits)'**.

[Note: The 6 protein hits shown in Main Figure 1D, constitute top-ranked hits consistently captured across 3 independent biological replicate-runs, whereas entries in Supplemental Table S3 are ranked in the order of highest-to-lowest 'average' LFQ scores. Thus, for instance, protein such as Tln1 does not appear within the list in Figure 1D, since its LFQ score in the second replicate experiment is only 6.72, although the average score (7.68) is high].

**Supplementary Table 4. Time-course RNA-Seq datasets at 3, 6, and 12 h time-points, following T-REX-assisted NCBP1-HNEylation, against all relevant T-REX controls (light alone, probe alone, and DMSO).**

Please see the enclosed excel file labeled '**Supplemental Table S4 (T-REX_NCBP1_RNASeq-hits)'**.

[Note: Within each tab of this excel file, hit transcripts with adj $p$ values < 0.05, are highlighted in red].

**Supplementary Table 5. The input list of top-ranked candidate SDEs and their respective biological functions.**

Please see the enclosed excel file labeled '**Supplemental Table S5 (SDEs_STRING_gProfiler)'**.

**Supplementary Table 6. RT-qPCR primers**

| gene | Primer set | Sequence (5'→3') |
|---|---|---|
| **DACH1 mRNA** | Forward primer | CAATGACTGCACCAACGCAA |
| | Reverse primer | GGAAGTTCCAGTCCGACACT |
| **DACH1 Pre-mRNA** | Forward primer | GTCACCATTTTAAACTTTATTAGGGTTCAGC |
| | Reverse primer | ATATGTGGCATCAAAGAGAGGAAATTTATATCA |
| **Actin (K118TAG)** | Forward primer | CCAGACTACGCAGGCTTCGA |
| | Reverse primer | AATCCTTCTGACCCATGCCCA |
| **POUSF2** | Forward primer | GCTCTTGGCCACTCTCCATT |
| | Reverse primer | TTGCCTTCGATAAAGCGGGT |
| **SNRPD3** | Forward primer | GTAGGCCAGAGCCGAACTC |
| | Reverse primer | TCCCCCGATATACCTCACCG |
| **S6K1** | Forward primer | CTGGAGCACCCCCATTCACT |
| | Reverse primer_E11_12 | GATGAGCTTGAACTTCTCCAGCG |
| | Reverse primer_E13_14 | CCACATATGTAAAACCCAGAAAGACCTG |
| **THOC2 isoform2** | Forward primer | GTGACTGACGATATAGCCAGTACCA |
| | Reverse primer | GCCAAAGCATCATATTCTTGGCGA |
| **THOC7 Isoform2 excluded** | Forward primer | TGGGAGCCGTGACTGACG |
| | Reverse primer | GCTCCAGCTATGCTACATTCTATTTCC |
| **MYBL2 Isoform2** | Forward primer | CCATGAGGAGAACCGCACTGA |
| | Reverse primer | GTGCCAGCGTTCACGGC |
| **MYBL2 Isoform2 excluded** | Forward primer | CCAGCCACTTCCCTAACCG |
| | Reverse primer | CAGTGTCCACTGCTTTGTGC |
| **GAPDH** | Forward primer | GACAGTCAGCCGCATCTTCT |
| | Reverse primer | AAATGAGCCCCAGCCTTCTC |

**Supplementary Table 7. Time-course differential splicing analysis: datasets at 3, 6, and 12 h time-points, following T-REX-assisted NCBP1-HNEylation, against all relevant T-REX controls (light alone, probe alone, and DMSO).**

Please see the enclosed excel file labeled '**Supplemental Table S7 (differential splicing analysis)'**. Events at each time point were deemed significant if they appear at least twice (against 3 independent biological replicates) across the 3 comparisons, namely, T-REX vs. light alone control, T-REX vs. probe alone control, and T-REX vs. DMSO vehicle control. The tables shown are already filtered for adjusted p-value <0.05 (adjusted using the Benjamin-Hochberg method).

**Supplementary Table 8. Schematic list of genes that altogether undergo statistically-significant alternative splicing (AS) events at either 3 or 6 h following NCBP1-HNEylation (data from Supp. Table S7: differential splicing analysis), against their respective statistically significant (SS) differential expression (DE) changes following NCBP1-HNEylation (data from Supp. Table S4: T-REX_NCBP1_RNA-Seq-hits)**

| **Gene name** | **AS event type** | **AS direction post T-REX** (against corresponding controls indicated in Supp. Table S7) | **Time (h) post T-REX** | **Do these genes show differential expression (DE)?** (note: unless otherwise indicated, all SS DE genes below were found only in the 12 h post T-REX against DMSO control set) |
|---|---|---|---|---|
| CCDC124 | AF | Loss | 3 | N |
| ABCA13 | A5 | Loss | 3 | N |
| ABCA13 | SE | Loss | 3 | N |
| ACAT1 | AF | Gain | 3 | N |
| ACAT1 | AF | Gain | 3 | N |
| ACAT1 | AF | Gain | 3 | N |
| ACAT1 | SE | Loss | 3 | N |
| ANKRD16 | SE | Gain | 3 | N |
| ARRDC2 | AF | Gain | 3 | N |
| ATF4 | A3 | Gain | 3 | Upregulation, (12h, T-REX against Light alone control), Upregulation, (12h, T-REX against DMSO |
| ATF4 | A3 | Gain | 3 | Upregulation, (12h, T-REX against Light alone control), Upregulation, (12h, T-REX against DMSO |
| ATF4 | A5 | Loss | 3 | Upregulation, (12h, T-REX against Light alone control), Upregulation, (12h, T-REX against DMSO |
| ATF4 | SE | Loss | 3 | Upregulation, (12h, T-REX against Light alone control), Upregulation, (12h, T-REX against DMSO |
| ATRAID | A5 | Gain | 3 | N |
| BMPR1A | A3 | Gain | 3 | N |
| BOLA1 | A3 | Loss | 3 | N |
| C1orf122 | A5 | Gain | 3 | N |
| C20orf27 | SE | Gain | 3 | N |
| CCDC124 | AF | Loss | 3 | N |
| CCT4 | SE | Gain | 3 | Y (Downregulation) |
| CD83 | AF | Loss | 3 | N |
| CDKN2A | AF | Loss | 3 | Y (Downregulation) |

| | | | | |
|---|---|---|---|---|
| CHCHD3 | AF | Gain | 3 | N |
| CKB | RI | Loss | 3 | N |
| COPB1 | AF | Loss | 3 | N |
| COPB1 | AL | Gain | 3 | N |
| COPB1 | AL | Gain | 3 | N |
| COPB1 | SE | Loss | 3 | N |
| COTL1 | SE | Gain | 3 | Y (Downregulation) |
| COTL1 | SE | Gain | 3 | Y (Downregulation) |
| COX6A1 | SE | Loss | 3 | N |
| CRIPT | AF | Loss | 3 | N |
| DDX5 | A5 | Gain | 3 | N |
| DDX5 | AF | Gain | 3 | N |
| DDX5 | AF | Gain | 3 | N |
| DDX5 | AF | Gain | 3 | N |
| DDX5 | AF | Gain | 3 | N |
| DDX5 | AF | Gain | 3 | N |
| DDX5 | AF | Gain | 3 | N |
| DDX5 | SE | Gain | 3 | N |
| DIDO1 | A5 | Loss | 3 | N |
| DIDO1 | AF | Loss | 3 | N |
| DNTTIP1 | SE | Gain | 3 | N |
| E2F3 | A3 | Gain | 3 | Y (Downregulation) |
| E2F3 | AF | Gain | 3 | Y (Downregulation) |
| EIF4EBP1 | AF | Loss | 3 | Upregulation, (12h, T-REX against Light alone control), Upregulation, (12h, T-REX against DMSO |
| EIF4EBP1 | AF | Gain | 3 | Upregulation, (12h, T-REX against Light alone control), Upregulation, (12h, T-REX against DMSO |
| EMC6 | A5 | Loss | 3 | Y (Downregulation) |
| FAM168B | A5 | Gain | 3 | N |
| FAM3C | A5 | Loss | 3 | N |
| FLYWCH2 | A5 | Loss | 3 | N |
| FOXO3 | AF | Loss | 3 | N |
| GAR1 | AF | Loss | 3 | Y (Downregulation) |
| GAR1 | AF | Loss | 3 | Y (Downregulation) |
| GTF2E2 | AF | Gain | 3 | N |
| HCFC1 | A3 | Loss | 3 | N |
| HCFC1 | A3 | Loss | 3 | N |
| HLA-C | A3 | Loss | 3 | N |
| INPPL1 | A3 | Gain | 3 | N |
| INPPL1 | SE | Loss | 3 | N |
| ISCA2 | A3 | Gain | 3 | N |
| ISCA2 | SE | Gain | 3 | N |
| KEAP1 | AF | Loss | 3 | N |
| KEAP1 | AF | Loss | 3 | N |
| LIMK1 | A3 | Gain | 3 | N |
| LIN7C | SE | Gain | 3 | Y (Downregulation) |

| | | | | |
|---|---|---|---|---|
| LRP3 | AF | Gain | 3 | N |
| MAP4K5 | A5 | Gain | 3 | N |
| MAP4K5 | A3 | Gain | 3 | N |
| MARCHF6 | AF | Loss | 3 | N |
| MARCHF6 | AF | Loss | 3 | N |
| MARCHF6 | AF | Loss | 3 | N |
| MARCHF6 | AF | Gain | 3 | N |
| MARCHF6 | AF | Gain | 3 | N |
| MBD3 | A3 | Gain | 3 | N |
| MEA1 | AF | Loss | 3 | Downregulation, (12h, T-REX against probe alone control), Downregulation, (12h, T-REX against DMSO) |
| MEA1 | A5 | Loss | 3 | Downregulation, (12h, T-REX against probe alone control), Downregulation, (12h, T-REX against DMSO) |
| MEA1 | AF | Gain | 3 | Downregulation, (12h, T-REX against probe alone control), Downregulation, (12h, T-REX against DMSO) |
| MEX3C | AF | Loss | 3 | N |
| MGARP | A5 | Gain | 3 | N |
| MGARP | AF | Gain | 3 | N |
| MGARP | SE | Gain | 3 | N |
| MGAT4C | A3 | Gain | 3 | N |
| MGAT4C | A3 | Gain | 3 | N |
| MIEN1 | A5 | Loss | 3 | N |
| MRPL13 | A5 | Loss | 3 | N |
| MRPL23 | AF | Loss | 3 | N |
| MRPL23 | AF | Loss | 3 | N |
| MRPL23 | RI | Gain | 3 | N |
| MRPL3 | A5 | Gain | 3 | Y (Downregulation) |
| MRPL3 | A5 | Loss | 3 | Y (Downregulation) |
| MYBL2 | SE | Gain | 3 | N |
| NA | A5 | Gain | 3 | N |
| NDUFAF3 | AF | Gain | 3 | N |
| NDUFAF3 | AF | Loss | 3 | N |
| NDUFAF3 | AF | Loss | 3 | N |
| NKAIN3 | SE | Gain | 3 | N |
| NKAIN3 | SE | Loss | 3 | N |
| PANK1 | AF | Loss | 3 | N |
| PAPPA | AL | Loss | 3 | N |
| PAPPA | SE | Gain | 3 | N |
| PCOTH | AF | Loss | 3 | N |
| PKDCC | A3 | Loss | 3 | N |
| PKMYT1 | AL | Loss | 3 | N |
| PKMYT1 | A3 | Gain | 3 | N |
| PKN1 | AF | Gain | 3 | N |
| PLEKHG4B | SE | Loss | 3 | N |
| PRKCSH | A5 | Loss | 3 | N |

| | | | | |
|---|---|---|---|---|
| PRKCSH | A5 | Loss | 3 | N |
| PRKCSH | AL | Loss | 3 | N |
| PRNP | A3 | Gain | 3 | N |
| PSMD3 | SE | Loss | 3 | N |
| PTMS | AF | Loss | 3 | N |
| PTMS | AF | Loss | 3 | N |
| PTMS | AF | Loss | 3 | N |
| PTP4A1 | AF | Loss | 3 | N |
| PTP4A1 | AF | Loss | 3 | N |
| PTP4A1 | AF | Loss | 3 | N |
| PTTG1IP | SE | Gain | 3 | Y (Downregulation) |
| RAB4A | A3 | Gain | 3 | N |
| RFX7 | SE | Gain | 3 | N |
| RNPS1 | AF | Loss | 3 | N |
| RNPS1 | AF | Loss | 3 | N |
| RNPS1 | AF | Loss | 3 | N |
| RPA2 | A5 | Gain | 3 | N |
| RPA2 | AF | Gain | 3 | N |
| RPA2 | AF | Gain | 3 | N |
| RPL29 | A5 | Gain | 3 | N |
| RPL29 | A5 | Gain | 3 | N |
| RPL29 | A5 | Gain | 3 | N |
| RPL29 | A5 | Gain | 3 | N |
| RPL29 | AF | Loss | 3 | N |
| RPL29 | AF | Loss | 3 | N |
| RPL29 | AF | Gain | 3 | N |
| RPL32 | A3 | Loss | 3 | N |
| RPL4 | AL | Gain | 3 | N |
| RPL9 | A3 | Gain | 3 | N |
| RPL9 | A5 | Gain | 3 | N |
| RPL9 | AF | Gain | 3 | N |
| RPL9 | SE | Gain | 3 | N |
| RPL9 | SE | Gain | 3 | N |
| RPL9 | SE | Gain | 3 | N |
| RPS6 | A5 | Loss | 3 | N |
| RSPH6A | A3 | Gain | 3 | N |
| SF3B2 | SE | Loss | 3 | Y (Upregulation) |
| SIGIRR | A5 | Gain | 3 | N |
| SLC16A1 | AF | Gain | 3 | Y (Downregulation) |
| SLC16A1 | AF | Gain | 3 | Y (Downregulation) |
| SLC25A3 | A5 | Loss | 3 | Y (Downregulation) |
| SLC7A3 | A3 | Gain | 3 | N |
| SLC7A5 | AF | Gain | 3 | Y (Upregulation) |
| SLC7A5 | AF | Loss | 3 | Y (Upregulation) |
| SMNDC1 | AF | Gain | 3 | Y (Downregulation) |
| SMNDC1 | AF | Gain | 3 | Y (Downregulation)) |

| | | | | |
|---|---|---|---|---|
| SMNDC1 | AF | Gain | 3 | Y (Downregulation) |
| SNRPD1 | A3 | Loss | 3 | N |
| STK17B | AF | Loss | 3 | N |
| STK17B | AF | Gain | 3 | N |
| STPG4 | AF | Gain | 3 | N |
| STPG4 | AF | Gain | 3 | N |
| STPG4 | AF | Gain | 3 | N |
| STPG4 | AF | Gain | 3 | N |
| TARS2 | AF | Loss | 3 | N |
| TARS2 | SE | Gain | 3 | N |
| TCEAL8 | SE | Gain | 3 | N |
| THOC7 | SE | Gain | 3 | N |
| TIMM13 | A5 | Loss | 3 | N |
| TIMP1 | SE | Gain | 3 | N |
| TMED4 | AL | Gain | 3 | N |
| TMEM250 | AF | Gain | 3 | N |
| TMEM44-AS1 | A3 | Gain | 3 | N |
| TMUB1 | AF | Gain | 3 | N |
| TNR | SE | Gain | 3 | N |
| TOMM40 | A3 | Gain | 3 | N |
| TOP2A/Y_RNA | AF | Loss | 3 | N |
| TST | AF | Gain | 3 | N |
| TTN | SE | Gain | 3 | N |
| TTN | SE | Gain | 3 | N |
| TTN | SE | Loss | 3 | N |
| TXLNG | A5 | Loss | 3 | Y (Downregulation) |
| UBN2 | AF | Gain | 3 | N |
| UFM1 | A5 | Loss | 3 | N |
| UQCC2 | SE | Gain | 3 | N |
| VAMP8 | AF | Loss | 3 | N |
| VPS37C | AF | Loss | 3 | N |
| VPS37C | AF | Loss | 3 | N |
| VPS37C | AF | Loss | 3 | N |
| XACT | AF | Gain | 3 | N |
| YDJC | AL | Loss | 3 | N |
| YDJC | A5 | Loss | 3 | N |
| YDJC | SE | Loss | 3 | N |
| YIPF5 | SE | Gain | 3 | N |
| ZFR | SE | Loss | 3 | N |
| ZNF319 | A5 | Loss | 3 | N |
| ZNF444 | A3 | Loss | 3 | N |
| ZNF460 | AF | Loss | 3 | Y (Upregulation) |
| ZNF460 | AF | Loss | 3 | Y (Upregulation) |
| ZNF460 | AF | Loss | 3 | Y (Upregulation) |
| ZNF764 | A3 | Loss | 3 | N |
| ZNF768 | AF | Gain | 3 | N |

| | | | | |
|---|---|---|---|---|
| ACO1 | A5 | Gain | 6 | N |
| ACO1 | SE | Loss | 6 | N |
| ACO1 | SE | Gain | 6 | N |
| ARHGAP5 | AF | Loss | 6 | N |
| ARHGAP5 | AF | Loss | 6 | N |
| ARPP19 | SE | Gain | 6 | Y (Downregulation) |
| B4GALT2 | A5 | Loss | 6 | N |
| B4GALT2 | AF | Loss | 6 | N |
| BATF3 | AF | Loss | 6 | N |
| BEND4 | A5 | Loss | 6 | Y (Downregulation) |
| BOLA1 | A3 | Gain | 6 | N |
| C19orf53 | AF | Gain | 6 | Y (Downregulation) |
| C19orf53 | AF | Loss | 6 | Y (Downregulation) |
| C19orf53 | A5 | Loss | 6 | Y (Downregulation) |
| C20orf27 | SE | Loss | 6 | N |
| CBL | SE | Gain | 6 | N |
| CBX6 | A3 | Gain | 6 | Y (upregulation) |
| CCNT2 | RI | Loss | 6 | N |
| CCZ1 | A3 | Gain | 6 | Y (Downregulation) |
| CCZ1 | SE | Gain | 6 | Y (Downregulation) |
| CD83 | AF | Gain | 6 | N |
| CDKN2A | AF | Gain | 6 | Y (Downregulation) |
| COX20 | A3 | Loss | 6 | N |
| CSTF1 | AF | Loss | 6 | N |
| DAD1 | A5 | Gain | 6 | N |
| DAZAP2 | SE | Loss | 6 | Y (Downregulation) |
| DEGS1 | AF | Loss | 6 | N |
| DIRAS1 | AF | Loss | 6 | N |
| ECE1 | AF | Loss | 6 | N |
| EEF1A2 | RI | Gain | 6 | N |
| EIF1 | SE | Gain | 6 | Upregulation, (12h, T-REX against probe alone control), Upregulation, (12h, T-REX against DMSO |
| EIF3E | AF | Loss | 6 | Downregulation, (3h, T-REX against DMSO control), Downregulation, (12h, T-REX against DMSO |
| EIF3E | A5 | Gain | 6 | Downregulation, (3h, T-REX against DMSO control), Downregulation, (12h, T-REX against DMSO |
| EIF4EBP1 | AF | Gain | 6 | Upregulation, (12h, T-REX against Light alone control), Upregulation, (12h, T-REX against DMSO |
| EMC10 | A5 | Loss | 6 | N |
| EMC10 | SE | Gain | 6 | N |
| EMC4 | A3 | Gain | 6 | N |
| EMC4 | SE | Loss | 6 | N |
| EMC6 | A5 | Loss | 6 | Y (Downregulation) |
| ERBB3 | AF | Gain | 6 | N |
| ETS2 | AF | Gain | 6 | N |

| | | | | |
|---|---|---|---|---|
| ETS2 | AF | Loss | 6 | N |
| FAM168B | A5 | Loss | 6 | N |
| FLNC | SE | Loss | 6 | N |
| GNB2 | AF | Gain | 6 | N |
| GNB2 | AF | Loss | 6 | N |
| GNB2 | A3 | Gain | 6 | N |
| GPT2 | A5 | Gain | 6 | N |
| HES6 | A3 | Loss | 6 | N |
| HLTF | A3 | Loss | 6 | N |
| IPO13 | A3 | Gain | 6 | N |
| IRX3 | SE | Gain | 6 | N |
| ISCA2 | A3 | Gain | 6 | N |
| ISCA2 | SE | Gain | 6 | N |
| IVNS1ABP | SE | Loss | 6 | Y (Downregulation) |
| KLC2 | A3 | Loss | 6 | N |
| KPNA1 | A5 | Gain | 6 | N |
| KPNA1 | A5 | Loss | 6 | N |
| KPNA1 | AL | Gain | 6 | N |
| LANCL2 | SE | Gain | 6 | N |
| LBX2-AS1 | A3 | Loss | 6 | N |
| LIN7C | SE | Gain | 6 | Y (Downregulation) |
| LINC02631 | A5 | Gain | 6 | N |
| LMBRD2 | A3 | Loss | 6 | N |
| MAP7D1 | RI | Loss | 6 | N |
| MAPK1 | SE | Gain | 6 | N |
| MAPRE2 | A5 | Loss | 6 | Y (Downregulation) |
| MEA1 | AF | Loss | 6 | Downregulation, (12h, T-REX against probe alone control), Downregulation, (12h, T-REX against DMSO) |
| MEA1 | AF | Loss | 6 | Downregulation, (12h, T-REX against probe alone control), Downregulation, (12h, T-REX against DMSO) |
| MEA1 | A5 | Gain | 6 | Downregulation, (12h, T-REX against probe alone control), Downregulation, (12h, T-REX against DMSO) |
| MEX3C | AF | Gain | 6 | N |
| MGARP | AF | Gain | 6 | N |
| MID1IP1 | A5 | Gain | 6 | N |
| MID1IP1 | SE | Gain | 6 | N |
| MRPL13 | A5 | Loss | 6 | N |
| MRPL36 | AF | Gain | 6 | N |
| MRPL51 | A5 | Loss | 6 | N |
| MRPL51 | A5 | Gain | 6 | N |
| MRPL51 | AF | Loss | 6 | N |
| MYBL2 | SE | Gain | 6 | N |
| NAA15 | AF | Gain | 6 | Y (Downregulation) |
| NACA | AF | Gain | 6 | N |
| NDUFAF3 | AF | Gain | 6 | N |

| | | | | |
|---|---|---|---|---|
| NDUFAF3 | AF | Loss | 6 | N |
| NKD2 | SE | Loss | 6 | N |
| NR6A1 | A3 | Gain | 6 | N |
| NR6A1 | A3 | Loss | 6 | N |
| NRBF2 | SE | Gain | 6 | N |
| NRGN | A3 | Gain | 6 | N |
| ODC1 | A5 | Gain | 6 | N |
| ODC1 | AF | Loss | 6 | N |
| PFDN1 | AL | Loss | 6 | N |
| PFDN1 | SE | Gain | 6 | N |
| PHF10 | AF | Gain | 6 | N |
| PHYH | AF | Loss | 6 | N |
| PKN1 | AF | Loss | 6 | N |
| PLXNB2 | A5 | Gain | 6 | N |
| POLR2G | AF | Loss | 6 | N |
| PPP1R10 | AF | Gain | 6 | N |
| PRKAB1 | AF | Loss | 6 | N |
| PTGES3 | AF | Gain | 6 | Y (Downregulation) |
| PTGES3 | MX | Loss | 6 | Y (Downregulation) |
| PTGES3 | SE | Gain | 6 | Y (Downregulation) |
| PTMA | AF | Gain | 6 | Y (Downregulation) |
| PTMA | AF | Gain | 6 | Y (Downregulation) |
| PTMA/U4 | AF | Gain | 6 | N |
| R3HDM4 | A3 | Gain | 6 | N |
| RACK1 | AF | Loss | 6 | N |
| RACK1 | RI | Loss | 6 | N |
| RAN | A5 | Gain | 6 | Y (Downregulation) |
| RAN | SE | Gain | 6 | Y (Downregulation) |
| RAN | A5 | Gain | 6 | Y (Downregulation) |
| RASSF8 | AF | Gain | 6 | N |
| RGS19 | AF | Loss | 6 | N |
| RNF139 | AF | Gain | 6 | N |
| RPL32 | A3 | Gain | 6 | N |
| RPL32 | A3 | Loss | 6 | N |
| RPL32 | A3 | Loss | 6 | N |
| RPL9 | A3 | Gain | 6 | N |
| RPL9 | A5 | Gain | 6 | N |
| RPL9 | AF | Gain | 6 | N |
| RPL9 | SE | Gain | 6 | N |
| RPL9 | SE | Gain | 6 | N |
| RPS14 | AF | Loss | 6 | N |
| RPS2 | AF | Gain | 6 | N |
| RPS2 | MX | Loss | 6 | N |
| RPS2 | RI | Gain | 6 | N |
| RPS6 | A5 | Loss | 6 | N |
| SBNO2 | A3 | Gain | 6 | N |

| | | | | |
|---|---|---|---|---|
| SBNO2 | A3 | Gain | 6 | N |
| SETD1B | A5 | Loss | 6 | N |
| SLC7A5 | AF | Gain | 6 | Y (upregulation) |
| SNU13 | AF | Gain | 6 | Y (Downregulation) |
| SNU13 | AF | Gain | 6 | Y (Downregulation) |
| SNU13 | SE | Gain | 6 | Y (Downregulation) |
| SRCAP | A5 | Loss | 6 | N |
| SRCAP | AF | Gain | 6 | N |
| STPG4 | AF | Gain | 6 | N |
| TARS2/MIR6878 | AF | Loss | 6 | N |
| TARS2/MIR6878 | SE | Gain | 6 | N |
| TBC1D14 | AF | Gain | 6 | N |
| TCEAL8 | SE | Gain | 6 | N |
| TERF2IP | A5 | Loss | 6 | N |
| TEX45 | AF | Loss | 6 | N |
| THOC7 | SE | Loss | 6 | N |
| TIMP1 | SE | Gain | 6 | N |
| TMEM14C | AF | Loss | 6 | N |
| TMEM14C | SE | Gain | 6 | N |
| TMEM14C | SE | Gain | 6 | N |
| TMEM250 | AF | Gain | 6 | N |
| TMEM250 | RI | Loss | 6 | N |
| TOMM40 | A3 | Gain | 6 | N |
| TOP2A | AF | Loss | 6 | N |
| TP53 | AF | Gain | 6 | Y (upregulation) |
| TP53 | AF | Gain | 6 | Y (upregulation) |
| TP53 | AF | Gain | 6 | Y (upregulation) |
| TP53 | AF | Loss | 6 | Y (upregulation) |
| TP53 | AF | Loss | 6 | Y (upregulation) |
| TP53 | AF | Loss | 6 | Y (upregulation) |
| TRAF2/MIR4479 | AF | Loss | 6 | N |
| TRIM23 | A5 | Gain | 6 | N |
| TRMT112 | A3 | Loss | 6 | N |
| TSC22D3 | A3 | Loss | 6 | N |
| TSC22D3 | AF | Loss | 6 | N |
| TSC22D3 | AF | Loss | 6 | N |
| TSN | AF | Gain | 6 | N |
| TSN | AF | Gain | 6 | N |
| TSN | AF | Gain | 6 | N |
| TSN | AF | Gain | 6 | N |
| TSN | AF | Gain | 6 | N |
| TSPAN3 | AF | Gain | 6 | N |
| TSPAN3 | AF | Gain | 6 | N |
| TSPEAR-AS2 | AL | Loss | 6 | N |
| TST | AF | Gain | 6 | N |
| TTK | A5 | Loss | 6 | N |

| | | | | |
|---|---|---|---|---|
| TWF2-DT | A3 | Gain | 6 | N |
| TXLNG | A5 | Loss | 6 | Y (Downregulation) |
| UBE2N | A5 | Gain | 6 | Downregulation, (12h, T-REX against probe alone control), Downregulation, (12h, T-REX against DMSO) |
| UBFD1 | A5 | Loss | 6 | N |
| UHRF1 | AF | Loss | 6 | N |
| UHRF1 | AF | Gain | 6 | N |
| UQCC2 | SE | Gain | 6 | N |
| USP22 | AF | Gain | 6 | N |
| USP22 | AF | Loss | 6 | N |
| XIST | AF | Loss | 6 | N |
| YDJC | A5 | Loss | 6 | N |
| YDJC | SE | Loss | 6 | N |
| YIPF5 | A3 | Gain | 6 | N |
| YIPF5 | A5 | Loss | 6 | N |
| YIPF5 | A5 | Gain | 6 | N |
| YIPF5 | SE | Gain | 6 | N |
| YWHAE | AL | Loss | 6 | N |
| ZBTB34 | SE | Loss | 6 | Y (upregulation) |
| ZBTB7A | AF | Loss | 6 | N |
| ZFR/MIR579 | SE | Loss | 6 | N |
| ZNF580 | AF | Loss | 6 | Y (Downregulation) |
| ZNF704 | AF | Gain | 6 | N |
| ZNF764 | A3 | Loss | 6 | N |
| ZNF764 | A3 | Gain | 6 | N |
| ZNF768 | A3 | Gain | 6 | N |
| ZNF768 | AF | Gain | 6 | N |

**Supplementary Table 9. Antibodies for Western blot.**

| Antibody | Source | Dilution |
|---|---|---|
| Anti-HaloTag (mouse) | Promega G9211 | 1:1000 (WB) |
| mouse monoclonal Anti Flag | Sigma, F3165 | 1:5000 (WB) |
| Anti-HA (HRP conjugated) (Rat) | Sigma H3663 | 1:1000(WB) |
| V5 Tag Monoclonal Antibody | Invitrgen R96025 | 1:5000 (WB) 1:200 (IF) |
| Anti-β-actin (HRP conjugated) (mouse) | Sigma A3854 | 1:10000 (WB) |
| Horse anti-mouse IgG (HRP-conjugated) | Cell Signaling #7076 | 1:5000 (WB) |
| Goat anti-rabbit IgG (HRP-conjugated) | Cell Signaling #7074 | 1:5000 (WB) |
| Anti-tubulin (HRP conjugated) | Cell Signaling #12351 | 1:5000 (WB) |
| Anti- S6K1 (Rab) (antibody 1) | Cell Signaling # 2708 | 1:2000 (WB) |
| Anti- S6K1 (Rab) (antibody 2) | Cell Signaling # 9202 | 1:2000 (WB) |
| Anti- phospho-S6K1 (Rab) | Cell Signaling # 9234 | 1:1000 (WB) |
| Anti- 4EBP1 | Cell Signaling # 9452 | 1:2000 (WB) |
| Anti- phospho-4EBP1 | Cell Signaling # 2855 | 1:1000 (WB) |
| Anti- eif2alpha | Cell Signaling # 9722 | 1:2000 (WB) |
| Anti- phospho-eif2alpha | Cell Signaling # 9721 | 1:1000 (WB) |
| Anti- DACH1 | Fisher Sentific AG #16877613 | 1:1000 (WB) |
| Anti-IPO5 | Abcam # ab187175 | 1:100 (IF) |
| Anti-Halo (rb pAb) | Promega G9281 | 1:1000 (IF) |
| Anti- Calnexin (ms mAb) | Santa Cruz sc-23954 | 1:200 (IF) |
| Anti-Cand1 | Insight Biotech, sc137055 | 1:100 (IF) |
| Anti-SF3A1 | Proteintech, 15858-1-AP | 1: 2000 (WB) |
| Anti-ALYREF | Cell Signaling # 12655 | 1:1000 (WB) |
| Anti-PRP4 | Abcam ab201684 | 1:1000 (WB) |
| Anti-PHAX | Proteintech 16481-1-AP | 1:800 (WB) |
| Donkey Anti-Mouse IgG Alexa Fluor 568 | Abcam ab175472 | 1:2000 (IF) |
| Donkey Anti-Rabbit IgG Alexa Fluor 488 | Invitrogen, A21206 | 1:2000 (IF) |

### Supplementary Figures

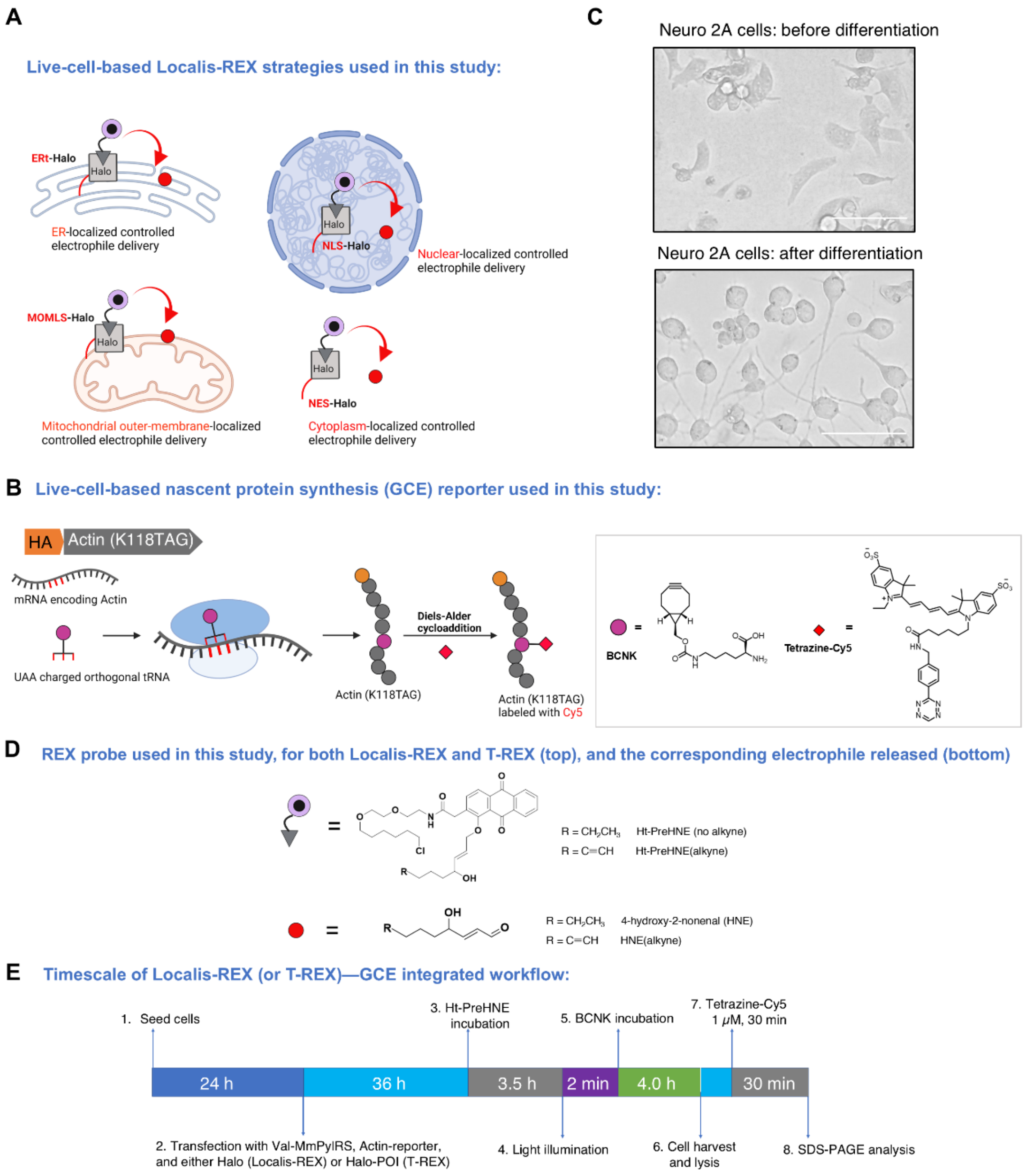


**Supplementary Fig. 1. Localis-REX, T-REX, and GCE general workflows. A)** Localis-REX setup: differentiated N2A cells ectopically expressing HaloTag and localized to 4 different subcellular locales as illustrated. The approach enables functional-electrophile responders in the specific locale, to be mapped with unparalleled control in timing, ligand chemotype and dosage, and locale. NLS, nuclear localization sequence; NES, nuclear export sequence, MOMLS, mitochondrial out membrane localization sequence; and ERt, ER-targeting sequence. **B)** GCE setup: Amber codon UAG was installed in synthetic mRNA of actin, to replace the codon encoding Lysine at 118$^{th}$ amino-acid position. Only after incubation with unnatural amino acid BCNK (see inset on the right for chemical structure), was translation of synthetic mRNA triggered, generating BCNK-integrated synthetic actin. Post cell lysis, a fluorophore can be attached to the synthetic actin, through Diels-Alder cycloaddition. *Inset:* Structure of BCNK and Tetrazine-Cy5. **C)** Representative bright field images of cultured mouse Neuro 2A (N2A) cells, before (*top*) and after (*bottom*) treatment with retinoic acid (20 µM, 24 h; differentiation inducer). Scale bar 100 µm. **D)** REX probes used in this study, for both localis-REX and T-REX (*top*), and the corresponding electrophile released (*bottom*). **E)** The workflow for GCE reporter assay, See **Method** for detail procedure.

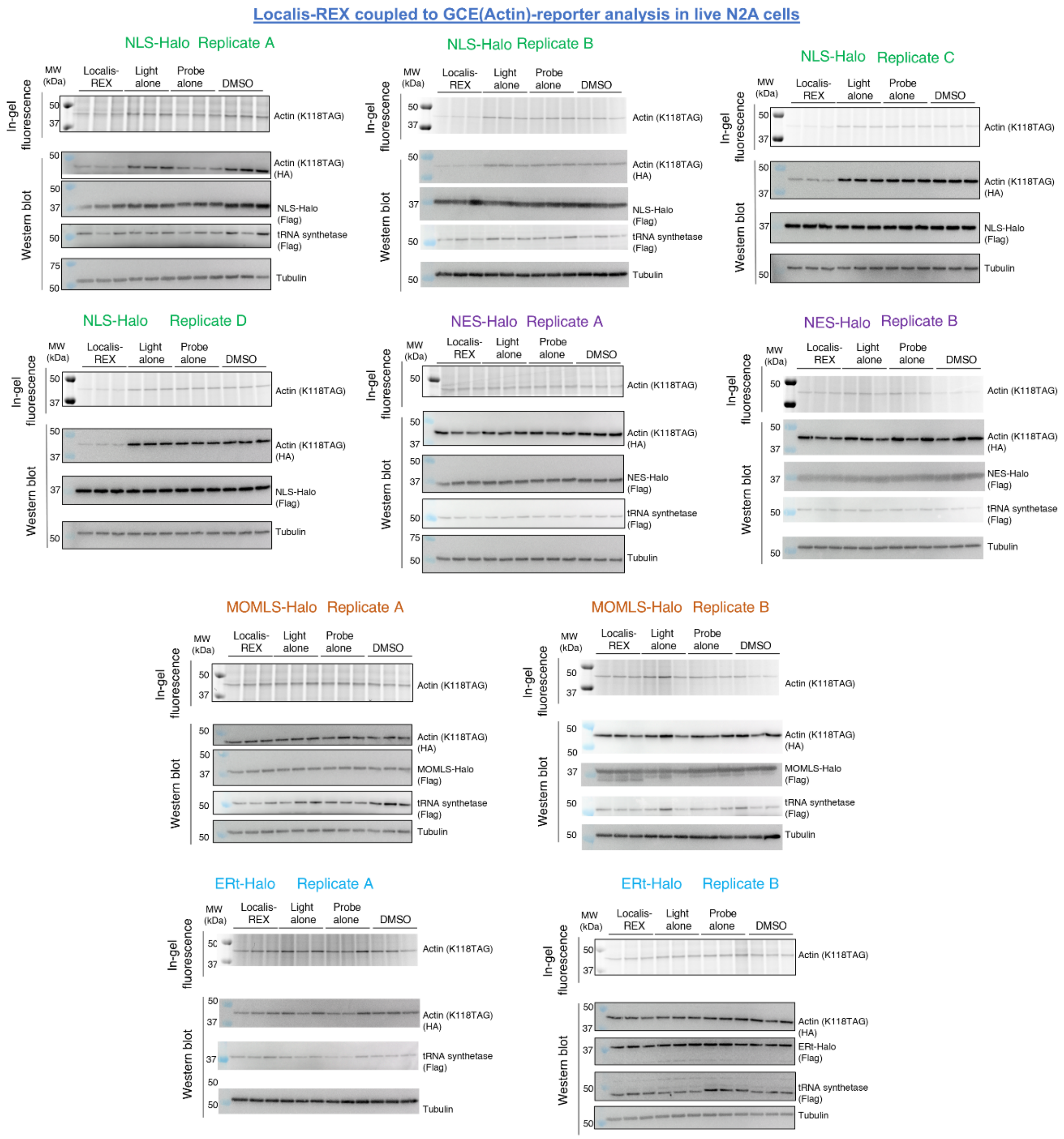


**Supplementary Fig. 2. GCE reporter assay, following Localis-REX at different subcellular locales in differentiated N2A cells, shows a prominent reduction of global protein synthesis in response to nuclear-targeted electrophile delivery.** Localis-REX—GCE assay (see **Fig. 1A-B** and **Methods**) was performed in differentiated N2A cells expressing either NLS-Halo, NELS-Halo, MOMLS-Halo, or ERt-Halo, and the data were analyzed by in-gel fluorescence and western blot (see **Methods** and also **Supplementary Fig. 1E** for details). NLS, nuclear localization sequence; NES, nuclear export sequence, MOMLS, mitochondrial out membrane localization sequence; and ERt, ER-targeting sequence. Light alone, Probe alone, and DMSO are technical negative controls associated with REX-technologies. See also **Fig. 1B-C**.

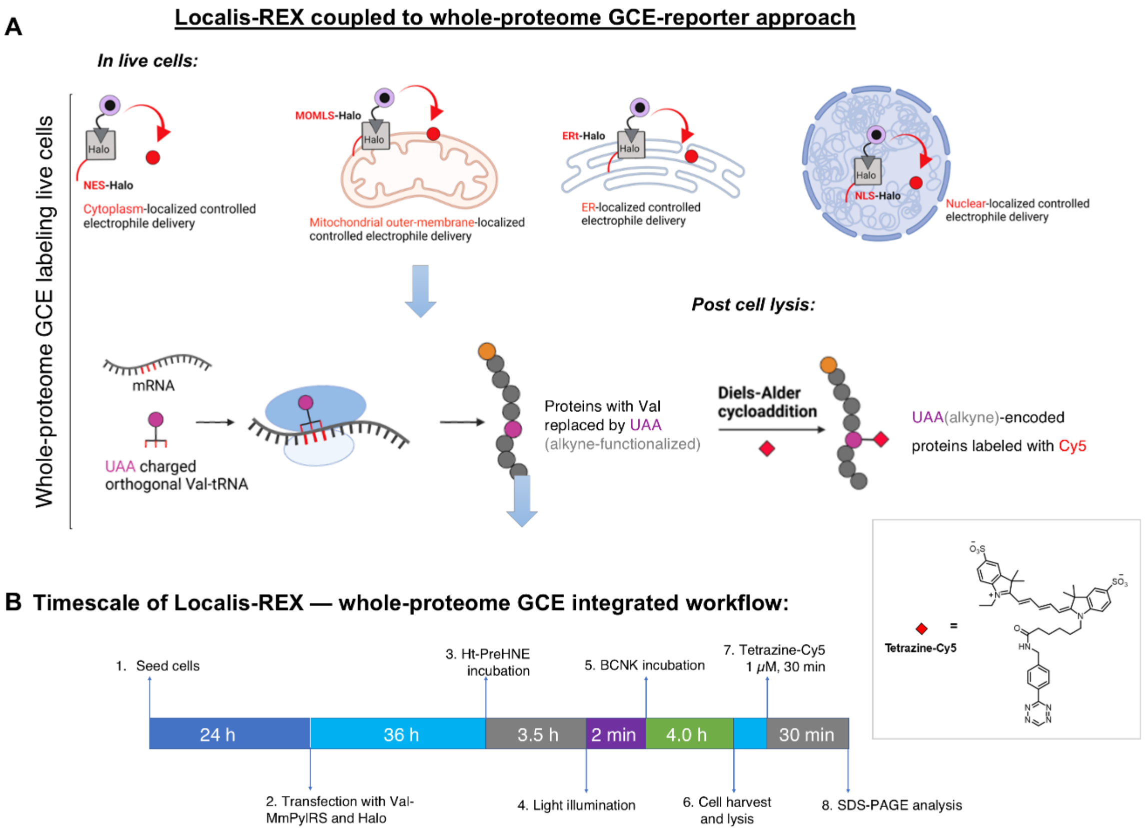


**Supplementary Fig. 3. Schematic diagram and workflow of Localis-REX coupled to whole-protemoe GCE labeling assay.** Experimental details were described in **Methods**. See also **Supplementary Fig. 4**.

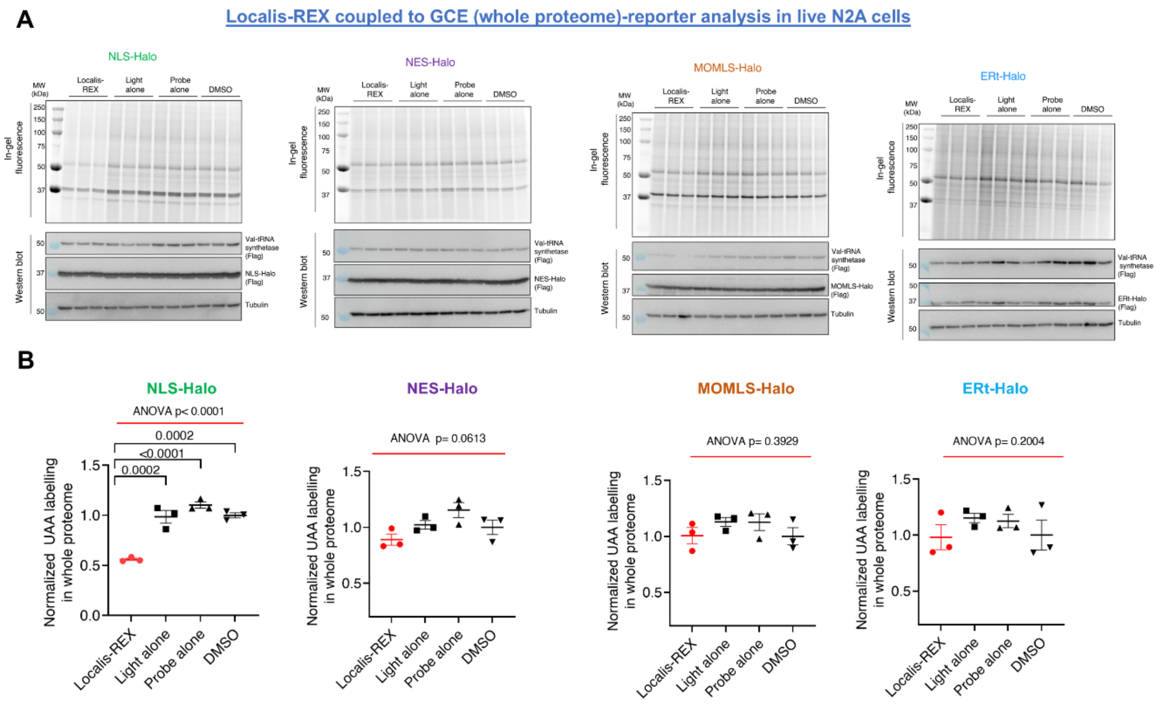


**Supplementary Fig. 4. GCE rseporter assay reporting on whole proteomic labeling, following Localis-REX at different subcellular locales, shows a prominent reduction of global protein synthesis in response to nuclear-targeted electrophile delivery. A)** The experiment was performed similarly as in **Supplementary Fig. 2**, except the omission of GCE-Actin reporter construct and the deployment of Val-MmPylRS as tRNA synthetase, whereby BCNK competed with valine to be incorporated into whole nascent proteome. (See also Schematic Workflow in **Supplementary Fig. 3**). See (B) for quantification. **B)** 'Normalized relative GCE-reporter signal' (y-axis) corresponds to relative values derived from the in-gel fluorescence signal of Cy5, normalized by that of anti-tubulin. p values were calculated using Tukey's multiple comparisons test (*n*=3). All data present mean ±SEM.

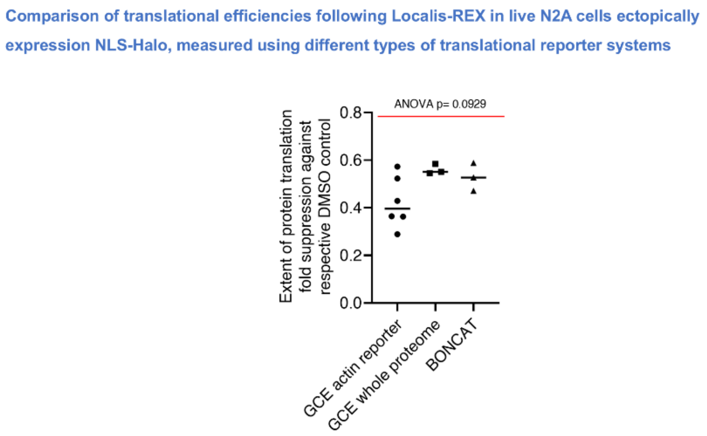


**Supplementary Fig. 5. The extent of protein translation suppression is similar across 3 different translation reporter assays, GCE actin reporter, GCE whole proteome, and BONCAT.** The degree of protein translation fold suppression upon Localis-REX in live N2A cells extopically expressing NLS-Halo (y-axis) corresponds to the relative values of either BCNK or AHA labelling, normalized to DMSO. Quantification of data shown in **Main Fig. 1B-C** and **Supplementary Fig. 2, 4 and Extended Data Fig. 3**. p values were calculated using Tukey's multiple comparisons test (*n*=6 for GCE actin reporter and n=3 for the rest). All data present mean ±SEM.

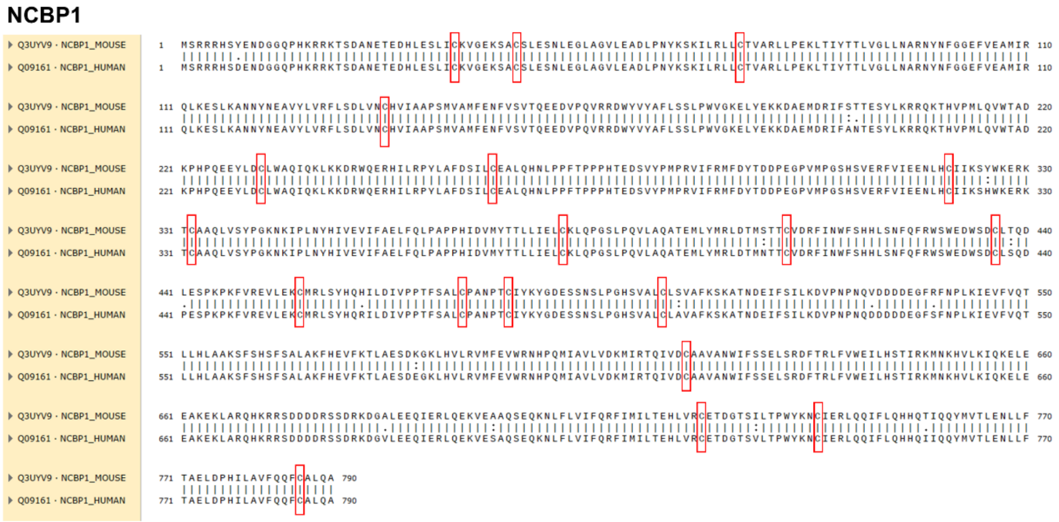


**Supplementary Fig. 6**. **Sequence alignment of human and mouse NCBP1 shows conservation of all cysteines**. Sequence alignment of human and mouse NCBP1 using SnapGene 6.1.1. Cysteines were highlighted in red boxes.

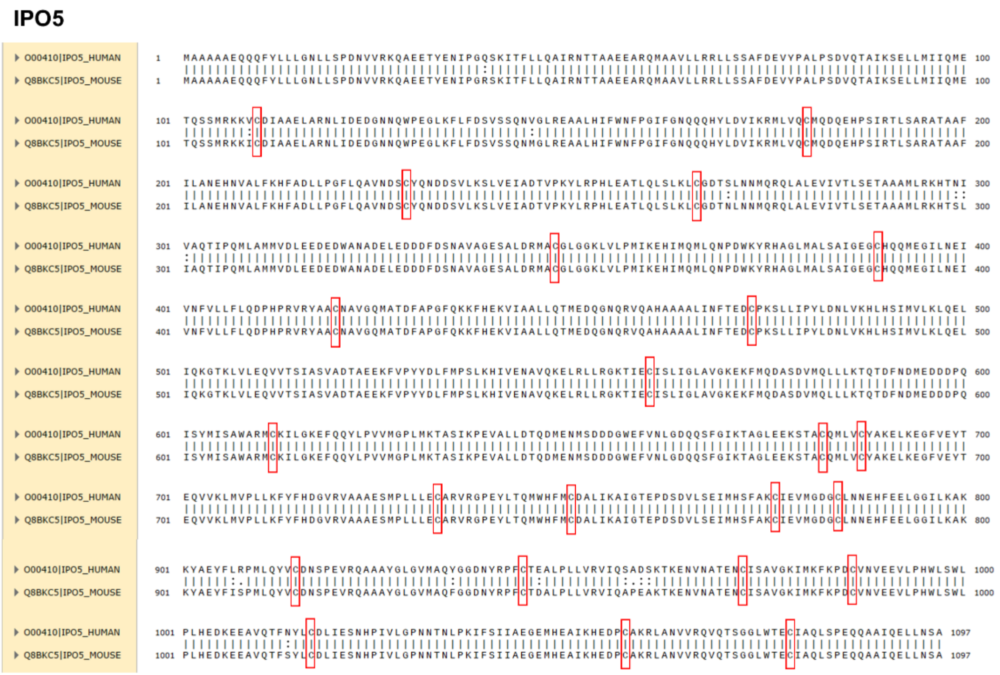


**Supplementary Fig. 7. Sequence alignment of human and mouse IPO5 shows conservation of all cysteines**. Sequence alignment of human and mouse IPO5 using SnapGene 6.1.1. Cysteines were highlighted in red boxes

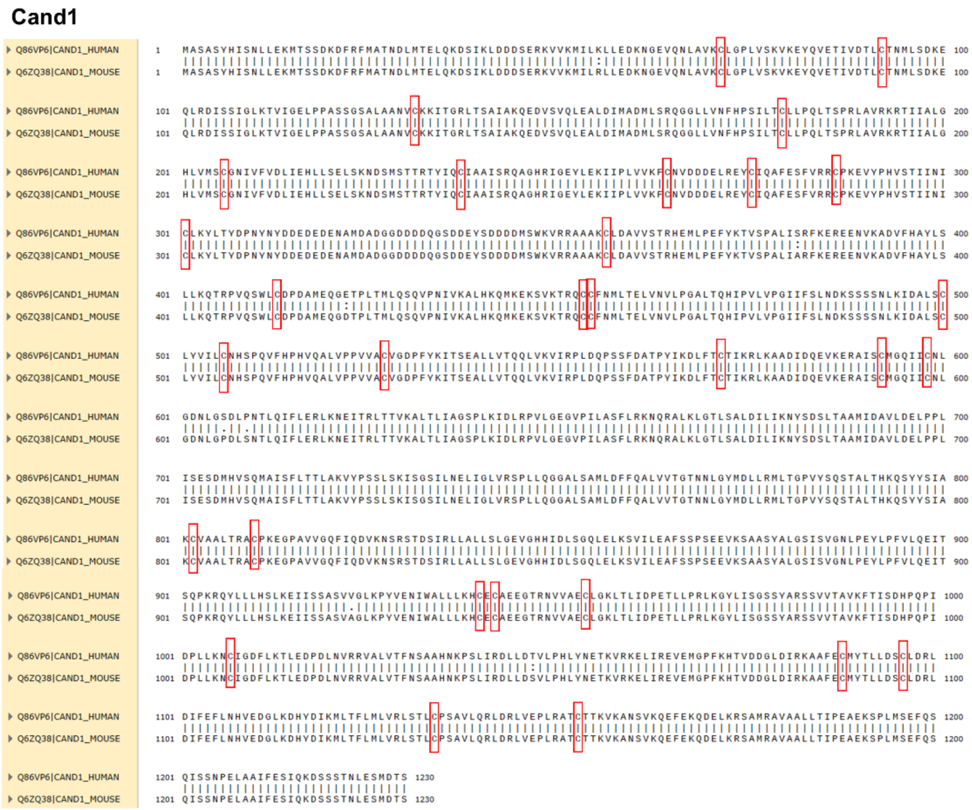


**Supplementary Fig. 8. Sequence alignment of human and mouse Cand1 shows conservation of all cysteines**. Sequence alignment of human and mouse Cand1 using SnapGene 6.1.1. Cysteines were highlighted in red boxes.

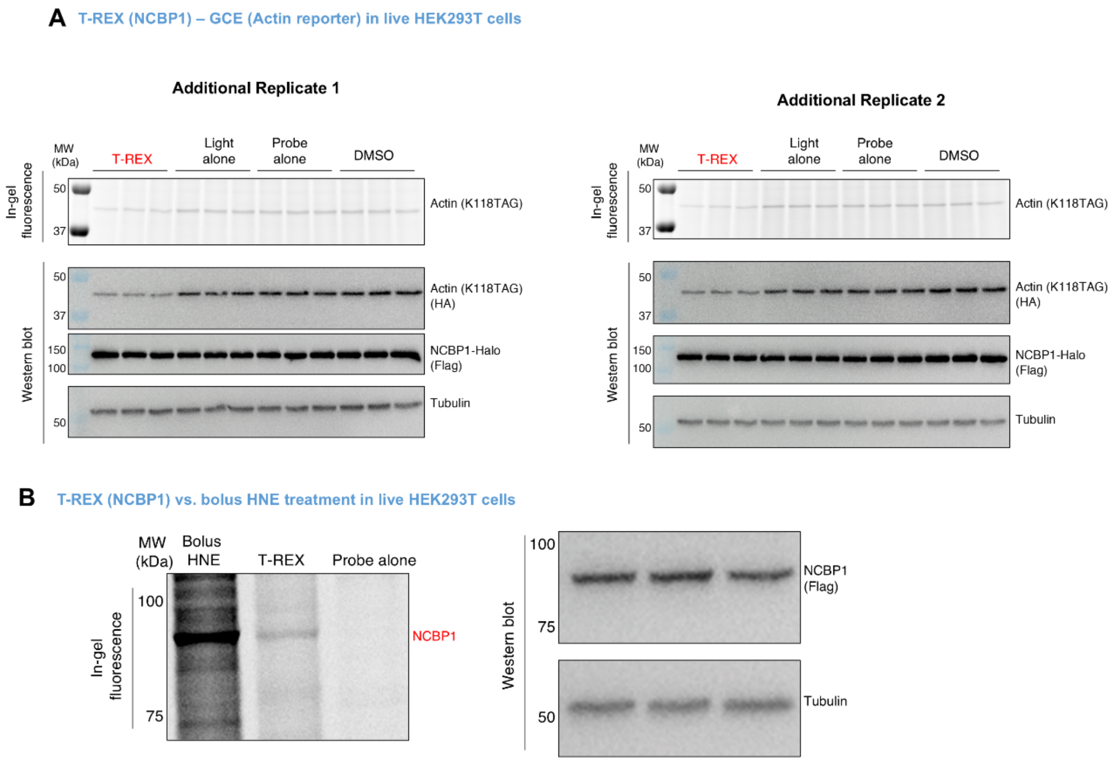


**Supplementary Fig. 9. Additional replicates of T-REX-GCE(actin) reporter assay involving NCBP1(wt)-Halo. Side-by-side comparsion between bolus HNE(alkyne) treatment and T-REX contrasts the relative extent of background non-specific modification with respect to HNEylation of ectopically expressed NCBP1-Halo. A**) Additional replicates of experiments in **Fig. 2F**. **B)** HEK293T extopically expressing NCBP1(wt)-Halo were either subjected to T-REX or photocaged probe-alone control (see **Methods**), or HNE(alkyne) (10 µM) treatment. In-gel fluorescence (top) and western blot (bottom) analyses were conducted following identical procedure as described in **Fig. 2B**.

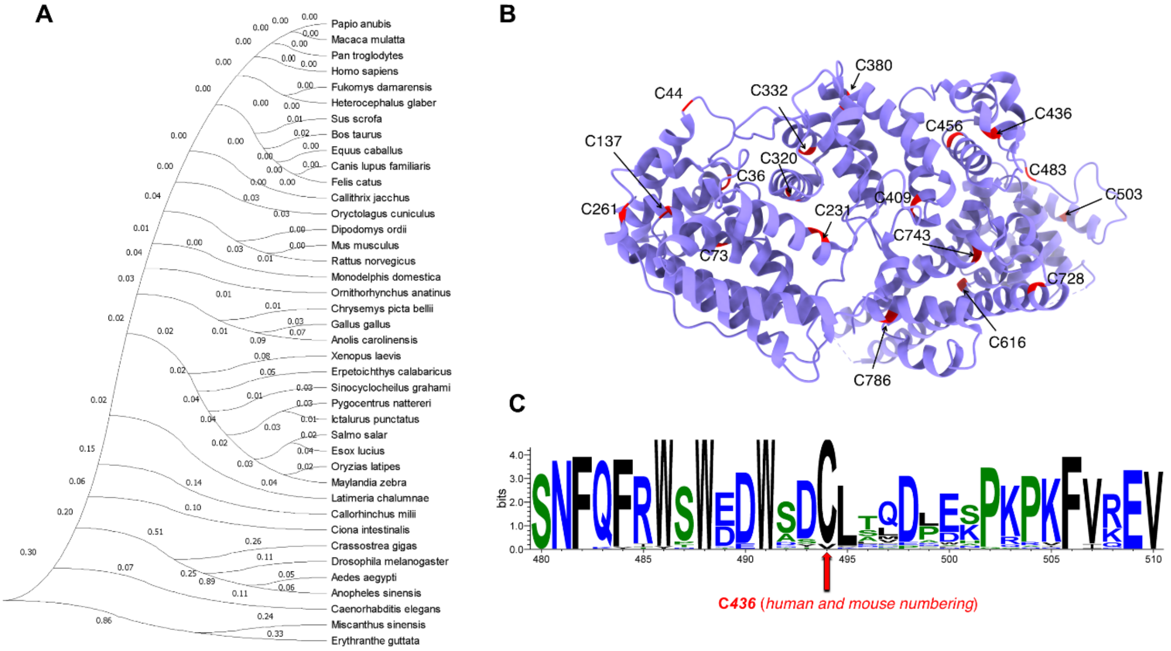


**Supplementary Fig. 10. Phylogenetic and structural analyses of NCBP1 alone and in complex with NCBP2**. **A)** The amino-acid conservation tree of NCBP1 across 414 eukaryotic species (constructed using MEGA). The number next to each node (ranging from 0 to 1, from lowest possible confidence to highest possible confidence) represents the phylogenetic confidence of the tree topology. **B)** Available single crystal X-ray structure of NCBP1 from human. Only NCBP1 is selectively shown here, from the 2.00 Å resolution single crystal structure of heterotrimeric complex of NCBP1 and NCBP2 from human (PDB: 1H6K). Also see **Fig. 3E**. 19 cysteine residues fully conserved between human and mouse NCBP1 (**Supplementary Fig. 6**) are indicated. **C)** Sequence logos featuring regions flanking the functional electrophile-signaling site discovered in this work (C436) within NCBP1. Sequence logo was generated using WebLogo.

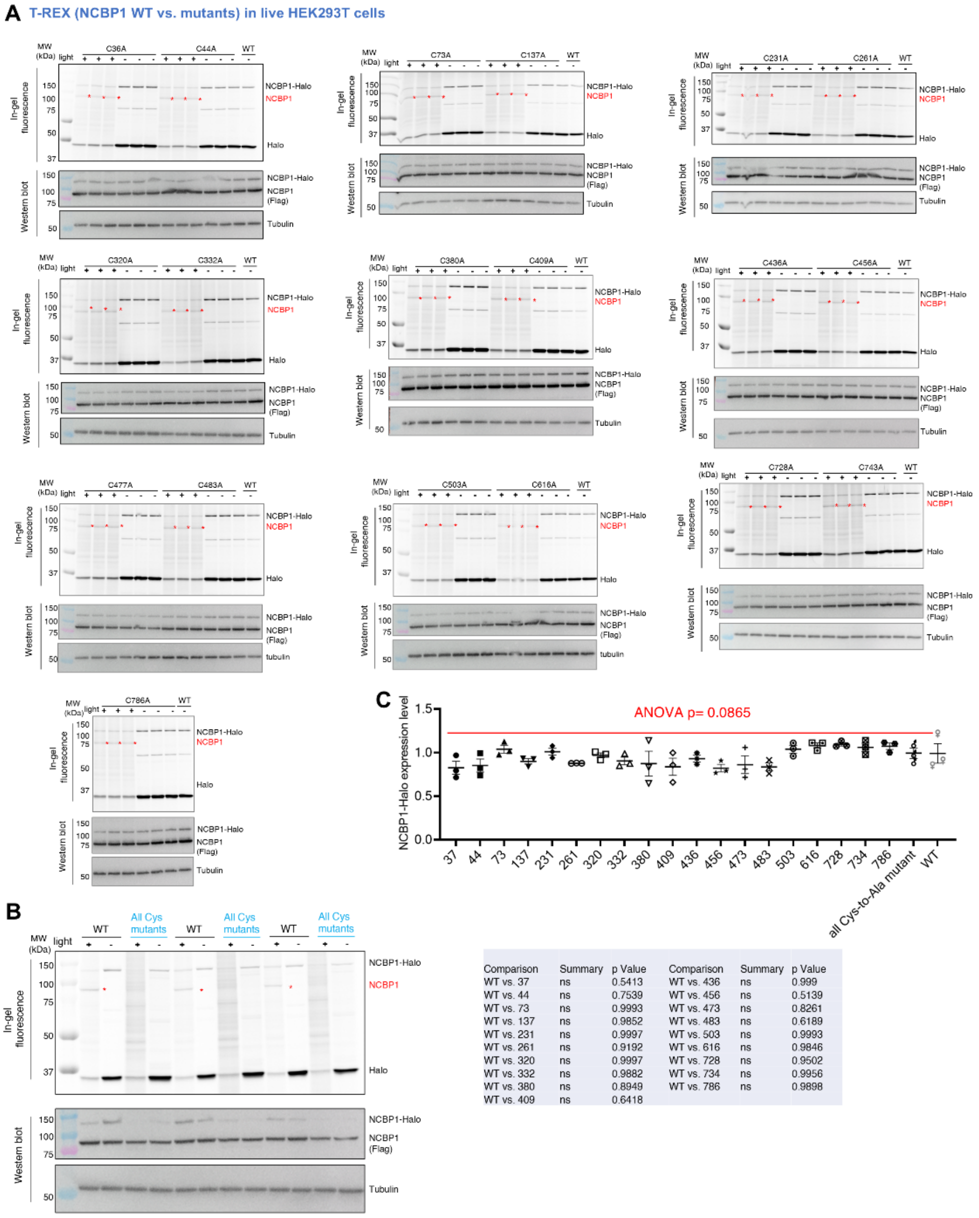


| Comparison | Summary | p Value | Comparison | Summary | p Value |
|---|---|---|---|---|---|
| WT vs. 37 | ns | 0.5413 | WT vs. 436 | ns | 0.999 |
| WT vs. 44 | ns | 0.7539 | WT vs. 456 | ns | 0.5139 |
| WT vs. 73 | ns | 0.9993 | WT vs. 473 | ns | 0.8261 |
| WT vs. 137 | ns | 0.9852 | WT vs. 483 | ns | 0.6189 |
| WT vs. 231 | ns | 0.9997 | WT vs. 503 | ns | 0.9993 |
| WT vs. 261 | ns | 0.9192 | WT vs. 616 | ns | 0.9846 |
| WT vs. 320 | ns | 0.9997 | WT vs. 728 | ns | 0.9502 |
| WT vs. 332 | ns | 0.9882 | WT vs. 734 | ns | 0.9956 |
| WT vs. 380 | ns | 0.8949 | WT vs. 786 | ns | 0.9898 |
| WT vs. 409 | ns | 0.6418 | | | |

**Supplementary Fig. 11. Functional mutagenesis coupled to T-REX evaluates potential HNE-sensing site(s) within NCBP1.** See also **Fig. 3A-B** and **Supplementary Table 1**. **A)** HEK293T ectopically expressing NCBP1-Flag-TEV-Halo (wt, or indicated Cys-to-Ala single-mutant) were subjected to T-REX (**Methods**. Also see **Fig. 2A-C** and **3A**). Briefly, datasets for each mutant show results from T-REX vs. control (without light exposure), indicated by '+' and '-', respectively) in three independent biological replicates. The band of HNEylated NCBP1 (post TeV-protease-assisted separation of NCBP1-Flag and Halo) was marked by *. See **Fig. 3A** for the corresponding relative ligand-occupancy quantification across all mutants studied vs. wild-type. Tubulin were used as a loading control. **B)** An independent biological replicate to that shown in **Fig. 2B**. **C)** Quantification of NCBP1-Halo ectopic expression levels across all 19 Cys mutants and wild-type (wt). The numbers show Cys to Ala single-mutation sites. The table below shows statistical analysis and power across pairwise comparison against wild-type. p values were calculated with Tukey's multiple comparisons test (*n*=3).

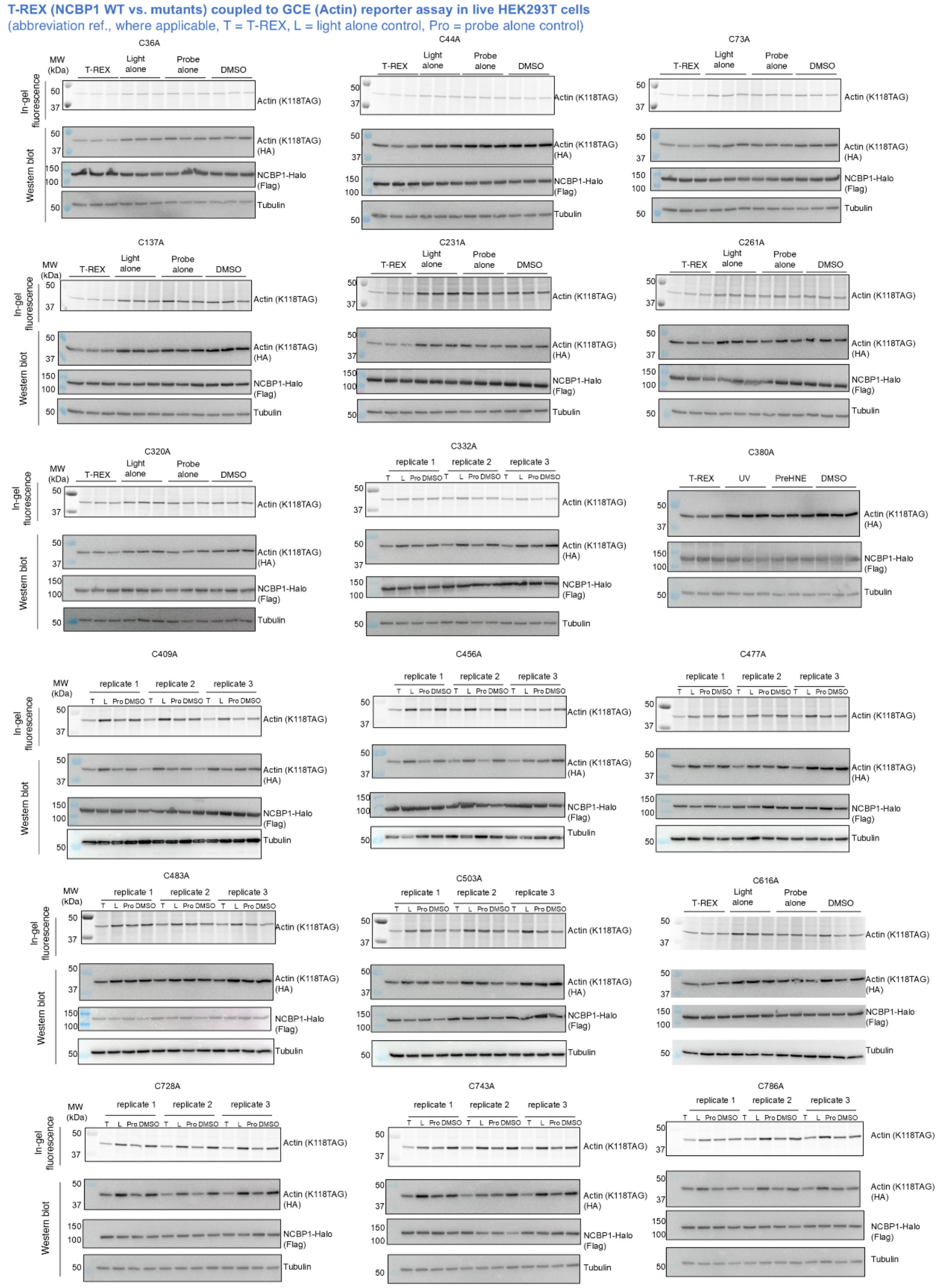


**Supplementary Fig. 12. T-REX coupled to GCE reporter assay leveraging mutant- vs. wt-NCBP1-Halo revealed that only C436-specific HNEylation functionally suppresses protein translation.** See also **Fig. 2F** and **3C-D.** Live-cell-based T-REX—GCE assay, as illustrated in **Supplementary Fig. 1E** workflow, was executed in HEK293T expressing (indicated NCBP1-mutant)-Halo (See **Methods**). See **Fig. 2F** and **3C** for comparison with results using NCBP1 (wt)-Halo, and corresponding quantification.

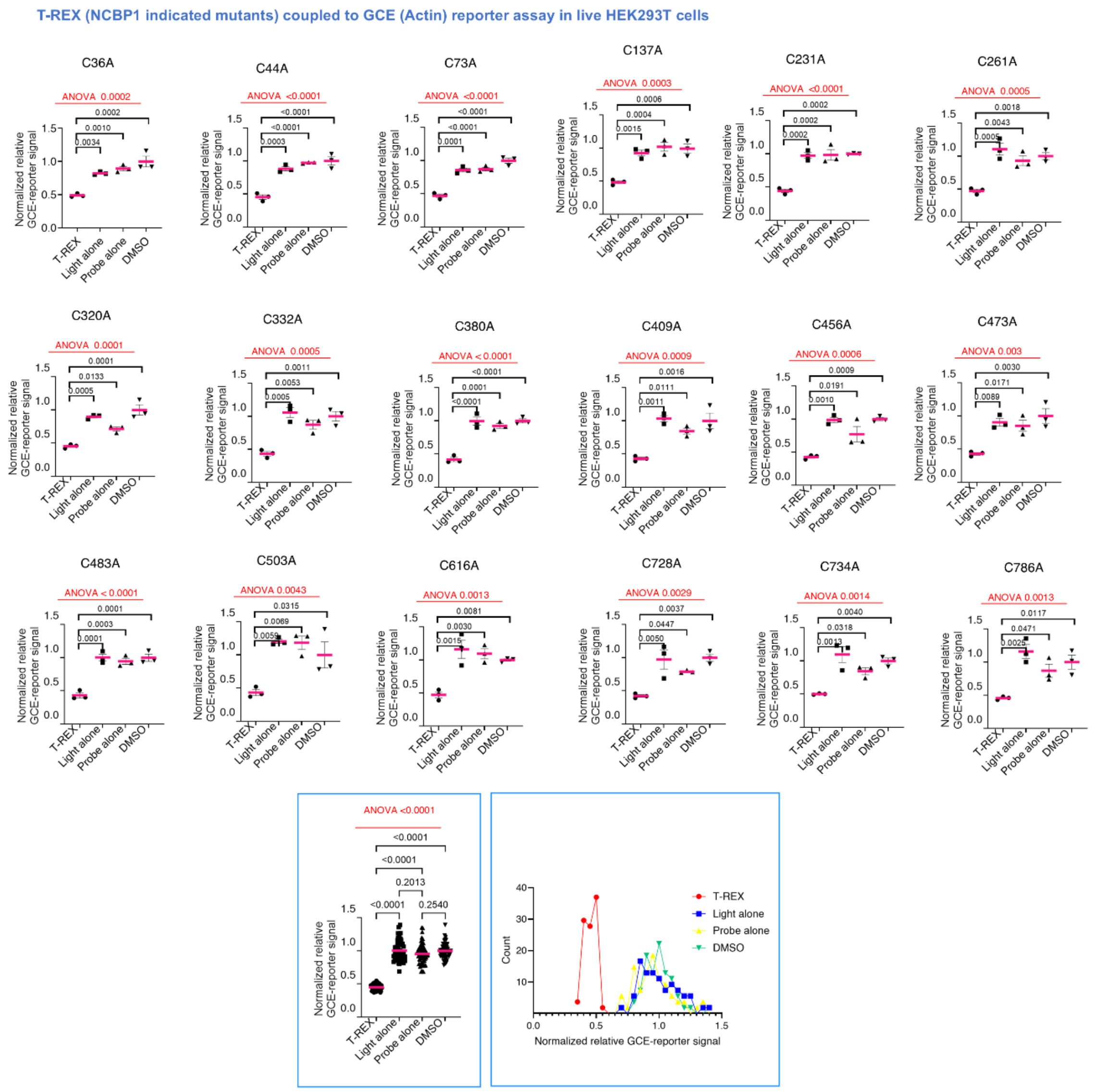


**Supplementary Fig. 13. Quantification of experimental results from Supplementary Fig. 12.** y-axis indicates relative values of GCE(actin) reporter signal normalized by NCBP1-Halo expression and tubulin expression levels. *P* values were calculated using Tukey's multiple comparison test (*n*=3). All data present mean ± SEM. The two boxed insets (bottom right) represent quantification of all the above data from all mutants combined, in two different representations (left: dot plot; right: histogram plot).

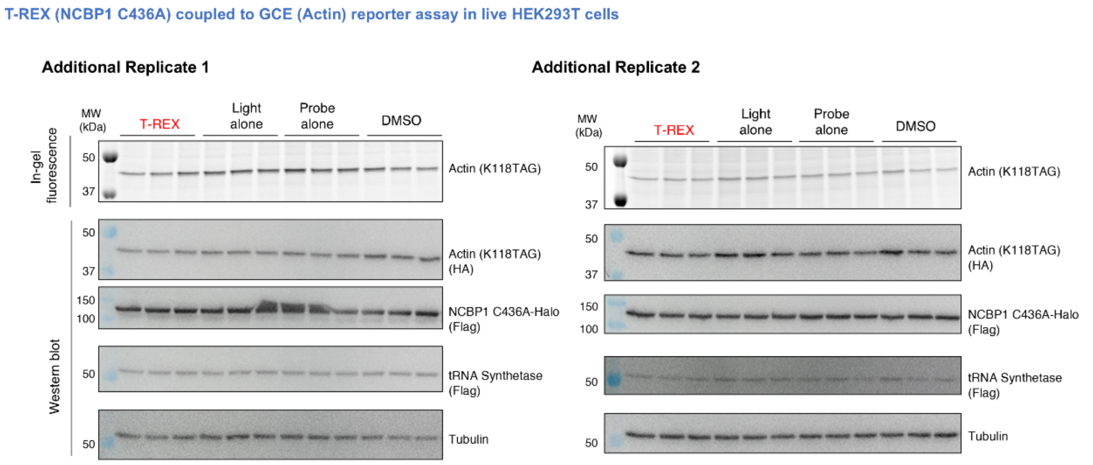


**Supplementary Fig. 14**. **T-REX coupled to GCE(actin) reporter assay involving NCBP1(C436A)-Halo.** Additional independent biological replicates to that shown in **Fig. 3C**.

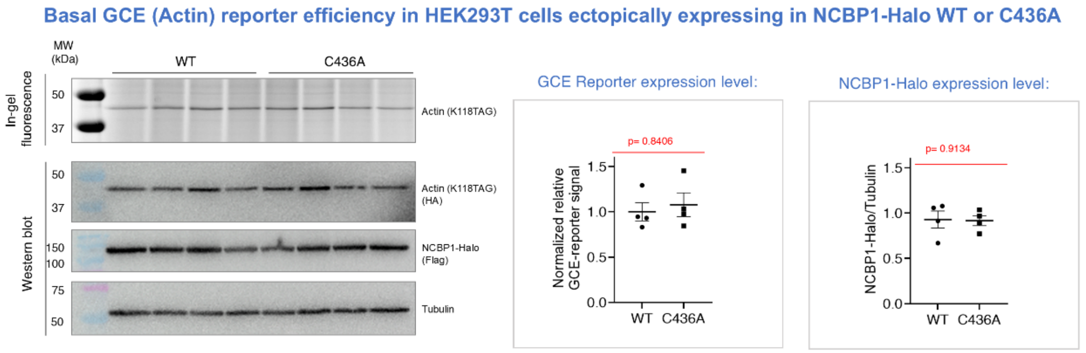


**Supplementary Fig. 15**. **The basal level of translation efficiency remained the same between cells expressing NCBP1(wt)-Halo and those expressing NCBP1(C436A)-Halo.** Live-cell-based GCE assay, as illustrated in **Supplementary Fig. 1E**, was executed in HEK293T expressing NCBP1(wt)-Halo or NCBP1 (C436A)-Halo (see **Methods**). Basel level of translation efficiency was examined for DMSO-treated control samples, as reported by Actin(K118TAG) expression level, normalized against NCBP1 (wt or C436A) and against loading control (tubulin). Insets show quantification of GCE reporter expression level and NCBP1-Halo expression level normalised over that of tubulin. p values were calculated with unpaired two-tailed student's *t*-test (*n*=4). All data present mean ±SEM.

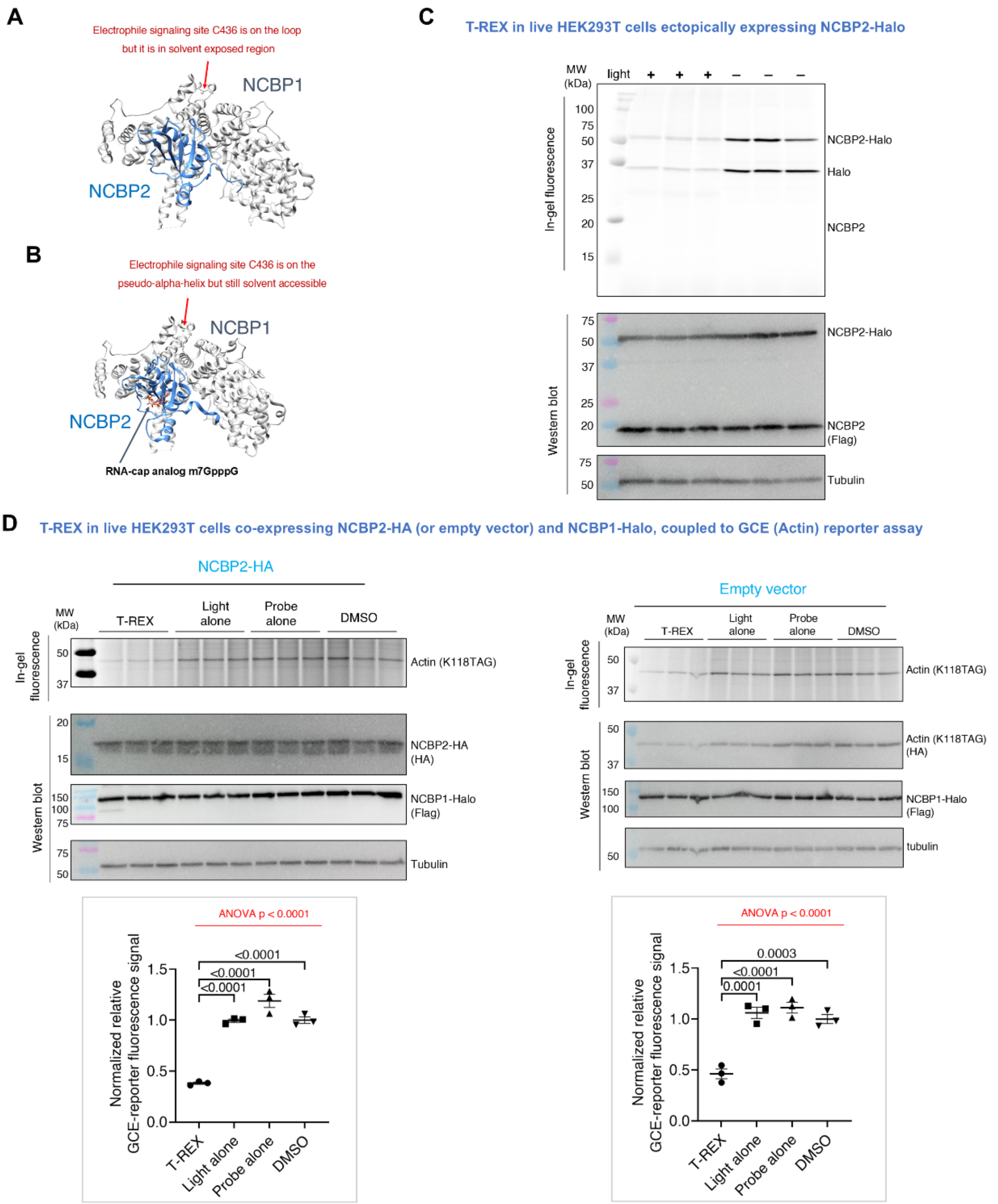


**Supplementary Fig. 16. NCBP1 binding partner, NCBP2, has no effect on the extent of translation suppression induced by NCBP1-HNEylation. A and B.** Crystal structures of NCBP1 and NCBP2, bound **(A)** and unbound **(B)** to capped mRNA. C436 (indicated with a red arrow) is in the solvent-exposed region in both cases. PDB: 1N52 for **A** and 1N54 for **B**. **C.** HEK293T expressing NCBP2-Flag-TeV-Halo were subjected to T-REX (as described in **Methods** and in **Fig. 2B-C** and **3A**). In-gel fluorescence analysis showed no detection of HNE-modified NCBP2. **D.** HEK293T co-expressing NCBP1-Flag-Halo, and either NCBP2-HA or empty vector (EV), replete with GCE-reporter constructs, were subjected to T-REX-GCE (as described in **Methods**). Cells were treated with BCNK (50 µM, 4 h). Post cell lysis, and coupling with tertrazine-Cy5, samples were analyzed by in-gel fluorescence (Cy5) reporting protein translation. *Insets at the bottom:* Quantification. p values were calculated using Tukey's multiple comparisons test (*n*=3). All data present mean ±SEM.

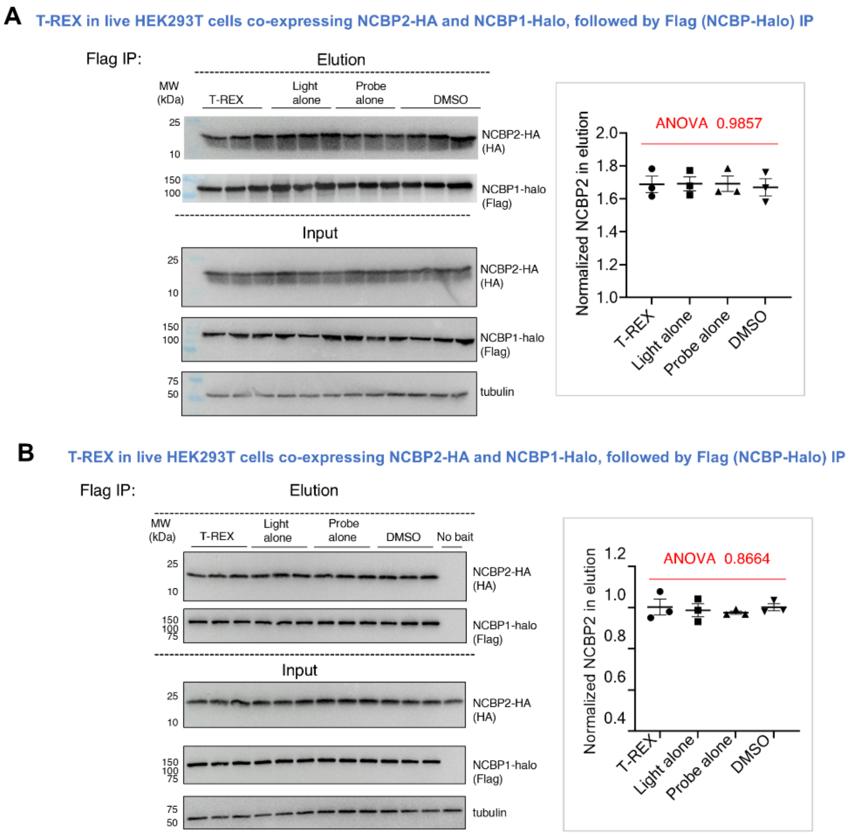


**Supplementary Fig. 17. NCBP1-NCBP2 intreaction is not affected by NCBP1-HNEylation. A)** Following T-REX-enabled NCBP1-HNEylation in HEK293T (see **Methods** for details) ectopically expressing NCBP1-Flag-Halo and NCBP2-HA, against indicated T-REX technical controls, Anti-Flag pulldown (see **Methods** for detail) was performed. Input and elution samples were subjected to SDS-PAGE, and western blot analyses using indicated antibodies. *Right panel* quantification of the protein level of eluted NCBP2-HA normalized by NCBP1-Halo expression. *p* values were calculated using Tukey's multiple comparison test (*n*=3). All data present mean±SEM. **B)** The experiment was performed identically to (**A**) above, except the addition of 'no bait' control (i.e., HEK293T cells transfected with empty vector in place of NCBP2-HA). *Right panel:* quantification of the protein level of eluted NCBP2-HA normalized by NCBP1-Halo expression. *p* values were calculated using Tukey's multiple comparison test (*n*=3). All data present mean±SEM.

T-REX (NCBP1) coupled to RNA-Seq at 6 h post T-REX in live HEK293T cells

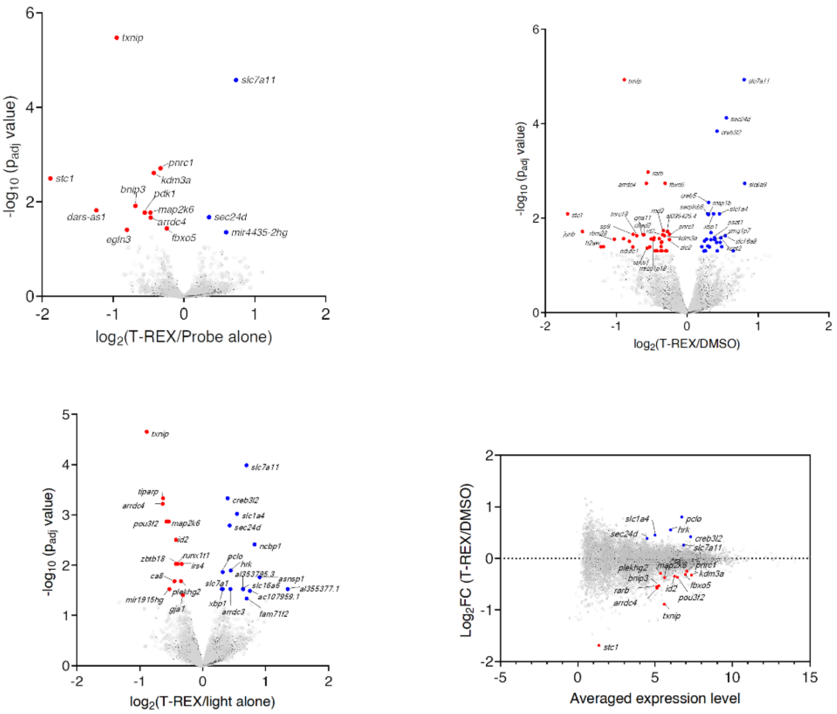


**Supplementary Fig. 18. Volcano and MA plots derived from RNA-seq analysis showing statistically-significant differentially-expressed genes (SDEs) under T-REX with respect to indicated controls (*6 h post T-REX*).** As in **Extended Data Fig. 9B-E**, except datasets are from 6 h time-point post T-REX. Select genes that were downregulated SDEs under T-REX were marked with **red** dots, and upregulated SDEs, with **blue** dots. Gene names were shown for arbitrarily-selected SDEs. See also **Supplementary Table 4**.

T-REX (NCBP1) coupled to RNA-Seq at 12 h post T-REX in live HEK293T cells

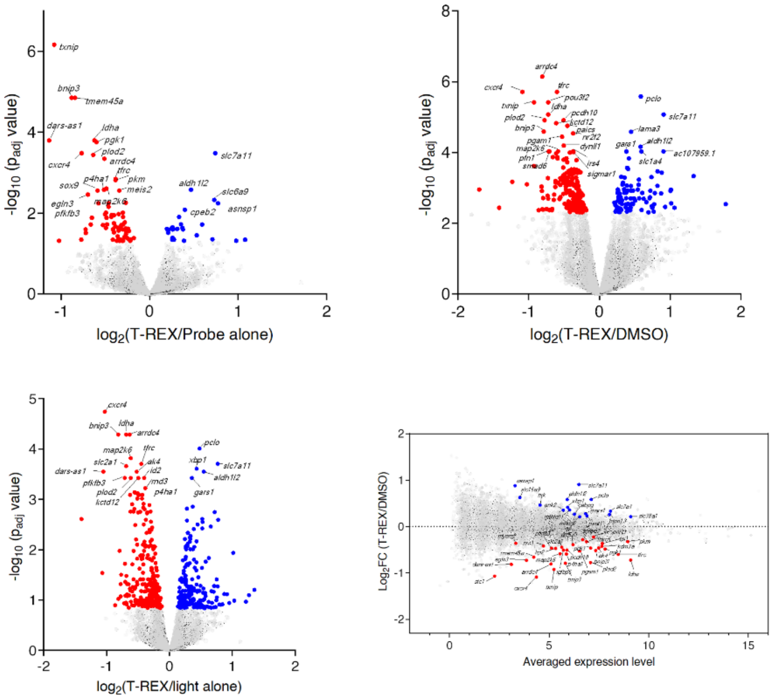


**Supplementary Fig. 19. Volcano and MA plots derived from RNA-seq analysis showing statistically-significant differentially-expressed genes (SDEs) under T-REX with respect to indicated controls (*12 h post T-REX*).** As in **Extended Data Fig. 9B-E**, except datasets are from 12 h time-point post T-REX. Select genes that are downregulated SDEs under T-REX were marked with **red** dots, and upregulated SDEs, with **blue** dots, and gene names are shown for arbitrarily-selected SDEs. See also **Supplementary Table 4**.

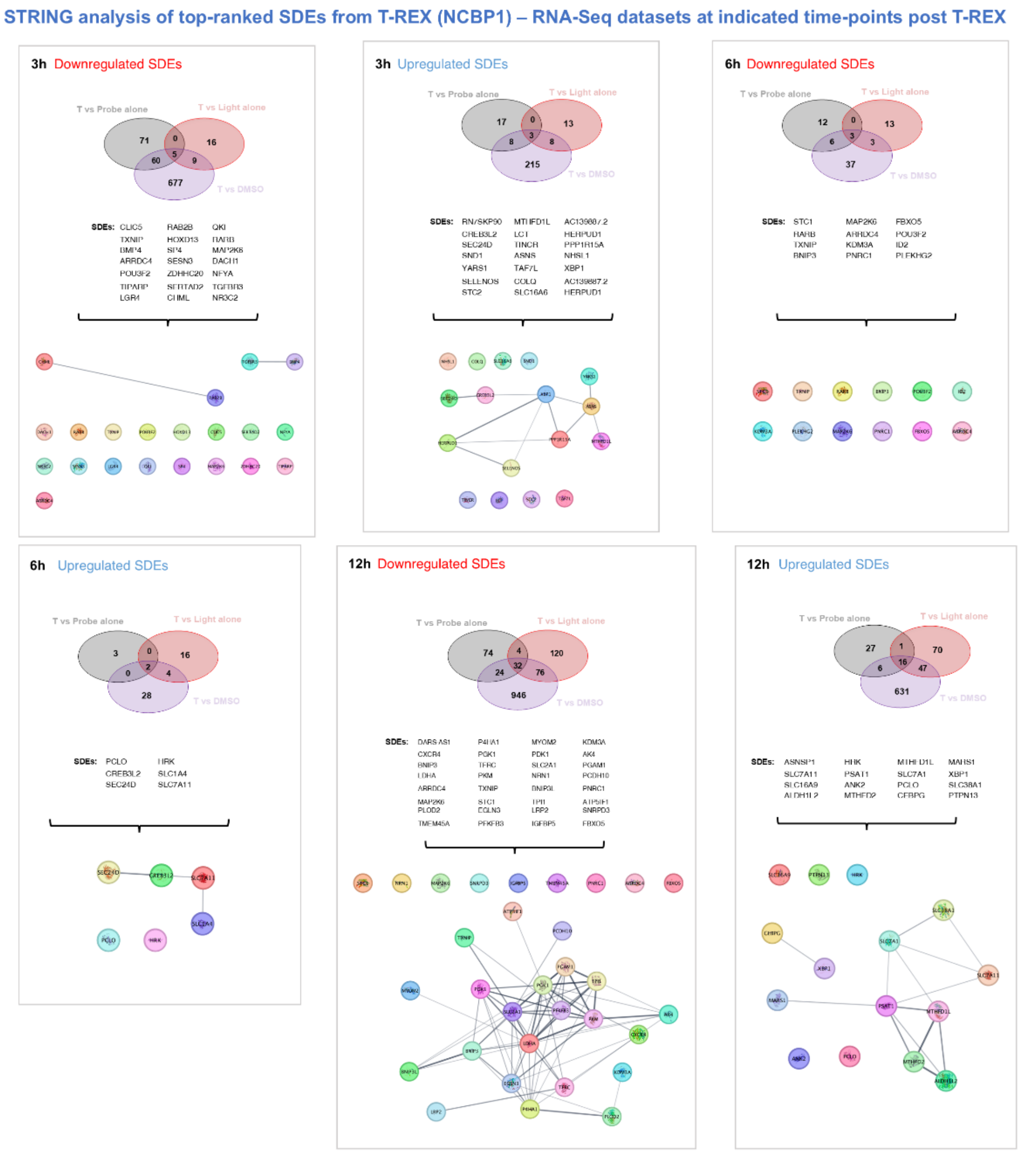


**Supplementary Fig. 20. STRING analysis of top-ranked SDEs from RNA-seq datasets.** The numbers within each Venn diagram correspond to the number of differentially-expressed genes (DEGs) altered in response to NCBP1-HNEylation, with respect to each indicated control group (light alone, probe alone, or DMSO treatments), in an either down- or upregulated manner, at a given time-point post NCBP1-HNEylation. [For instance, (71 (unique) +60 (overlapping with DMSO) +5 (overlapping with light alone and DMSO) DEGs were downregulated following T-REX with respect to probe-alone control, 3h post uncaging]. Subsequently, DEGs manifesting $p \leq 0.05$ under at least two experimental sets, out of the 3 sets of comparisons deployed (namely: T-REX against the 3 respective controls), were considered as statistically-significant DEGs (SDEs), which are listed underneath each Venn diagram, along with the output from STRING analysis, using default cut-off thresholds in Cytoscape (v3.10.1). See also **Supplementary Table 5**.

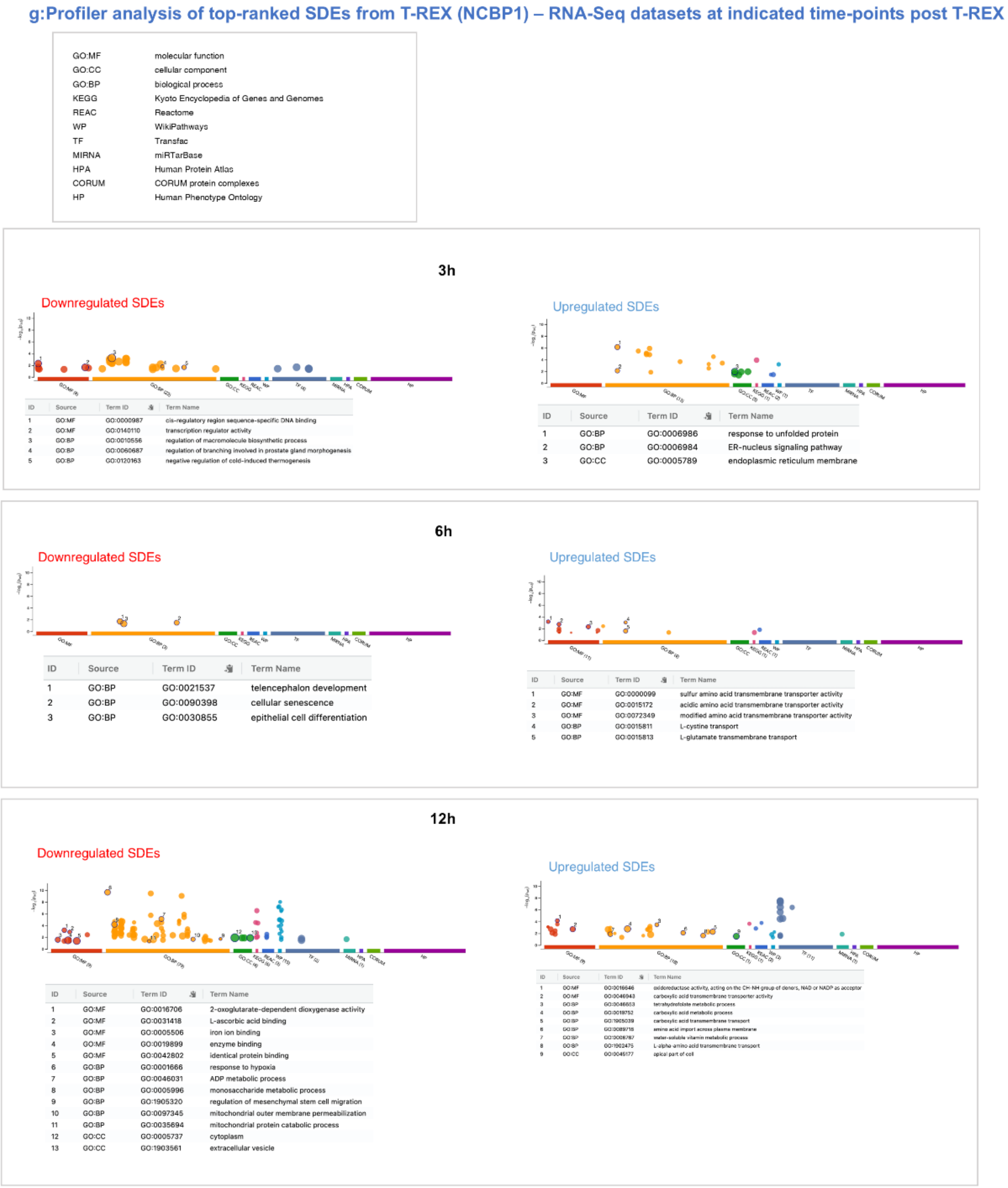


**Supplementary Fig. 21. *g:Profiler* analysis of top-ranked SDEs from RNA-seq datasets.** SDEs shown in the **Supplementary Fig. 20** were subjected to online software *g:Profiler* (https://bio.tools/gprofiler). See also **Supplementary Table 5**.

Primer efficiency analysis for primers targeting *dach1* mRNA or pre-mRNA

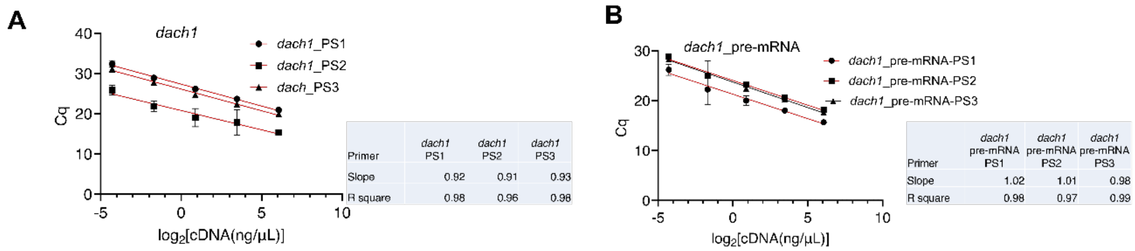


**Supplementary Fig. 22. Primer efficiency tests of *DACH1* mRNA and pre-mRNA.** Three sets of primers were designed and tested for each indicated gene as described in **Methods**. Briefly, five or six serial dilutions (1:6) of cDNA obtained from HEK293T were tested to obtain the quantification for cycle (Cq) values. [n = 6 replicates (3 biological replicates with 2 technical replicates); error bars represent SEM; linear regression analysis was performed, yielding the results shown in the table]. Primer efficiency values of 90–110% were considered as the selection criteria for primer sets. The respective primer set (PS) 1 was selected in each case: (A) *dach1* mRNA and (B) *dach1* pre-mRNA. See also **Supplementary Table 6**.

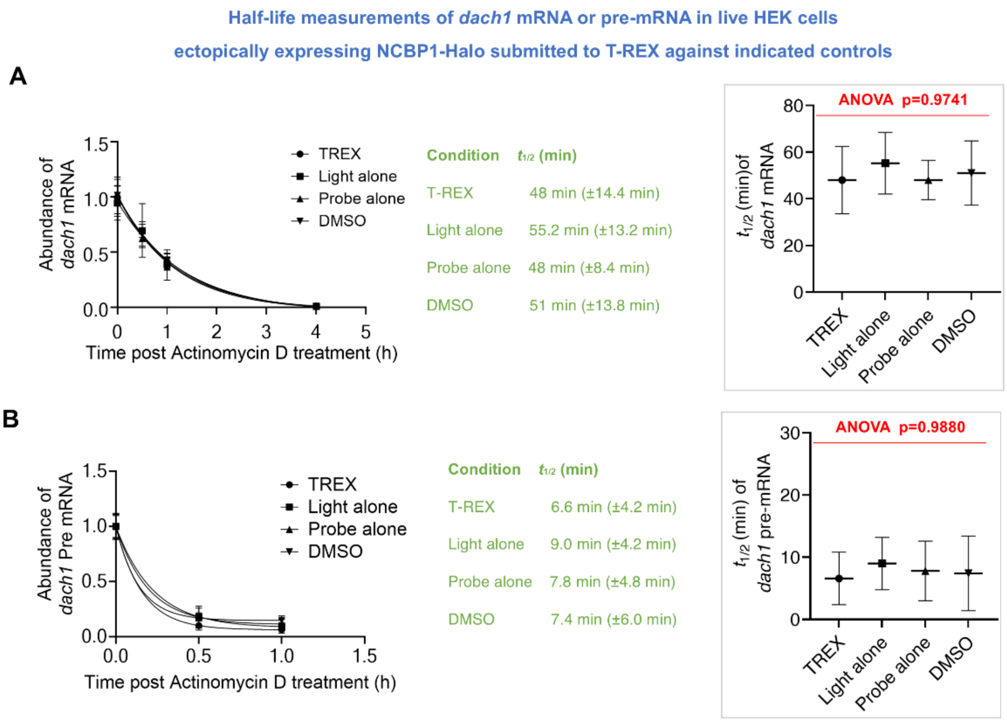


**Supplementary Fig. 23. NCBP1-HNEylation does not significantly perturb the half-life of *dach1*-pre-mRNA and -mRNA** HEK293T expressing NCBP1-Halo were exposed to T-REX and all other three controls, then Actinomycin D (5 µg/mL) was incubated into cells for the time period indicated (see detail protocol in **Methods**). RNA was extracted and the initial level of mRNA or pre-mRNA was set at 1, and data for other time points were normalized against that from time zero. Data were fit to one-phase exponential decay: y = (1-Plateau)*exp (-*k*x) + Plateau, where y is the RNA transcript levels, *k* is the rate constant, and x is the time, using Prim v9.4.0. **A)** results for *dach1*-pre-mRNA **B)** results for *dach1*-mRNA. $t_{1/2}$ of pre-mRNA decay shown in **A**) ($t_{1/2}$ under T-REX is 6.6 min (±4.2 min), $t_{1/2}$ under Light alone is 9.0 min (±4.2 min), $t_{1/2}$ under Probe alone is 7.8 min (±4.8 min), $t_{1/2}$ of DMSO is 7.4 min (±6.0 min). $t_{1/2}$ of mRNA decay shown in **B**) ($t_{1/2}$ under T-REX is 48 min (±14.4 min), $t_{1/2}$ under Light alone is 55.2 min (±13.2 min), $t_{1/2}$ under Probe alone is 48 min (±8.4 min), $t_{1/2}$ of DMSO is 51 min (±13.8 min). *p* values were calculated withTukey's multiple comparison test (*n*=9). All data present mean ± SEM.

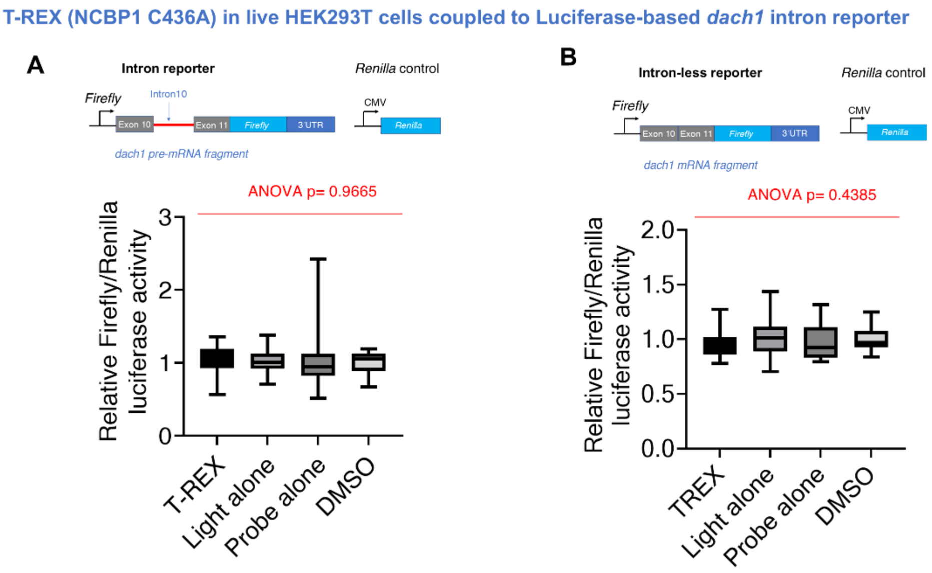


**Supplementary Fig. 24. NCBP1(C436A), HNE-signaling-dead mutant, is recalcitrant to NCBP1-HNEylation-induced perturbation of DACH1 splicing observed with NCBP1(wt).** See also **Fig. 4B-D.** The experimental setup was identical to that in **Fig. 4D** expect that HEK293T cells were transfected with NCBP1(C436)-Halo in place of NCBP1(wt)-Halo. **A)** Schematic of the *firefly* luciferase intron reporter was used, and **B)** Schematic of the *firefly* luciferase intron-less reporter was used. *p* values were calculated using ANOVA and Tukey's multiple comparison test (*n*=18). All data present mean±SEM. Box and whisker plots: line-median; box-25-75 percentiles; whiskers-1-99 percentiles.

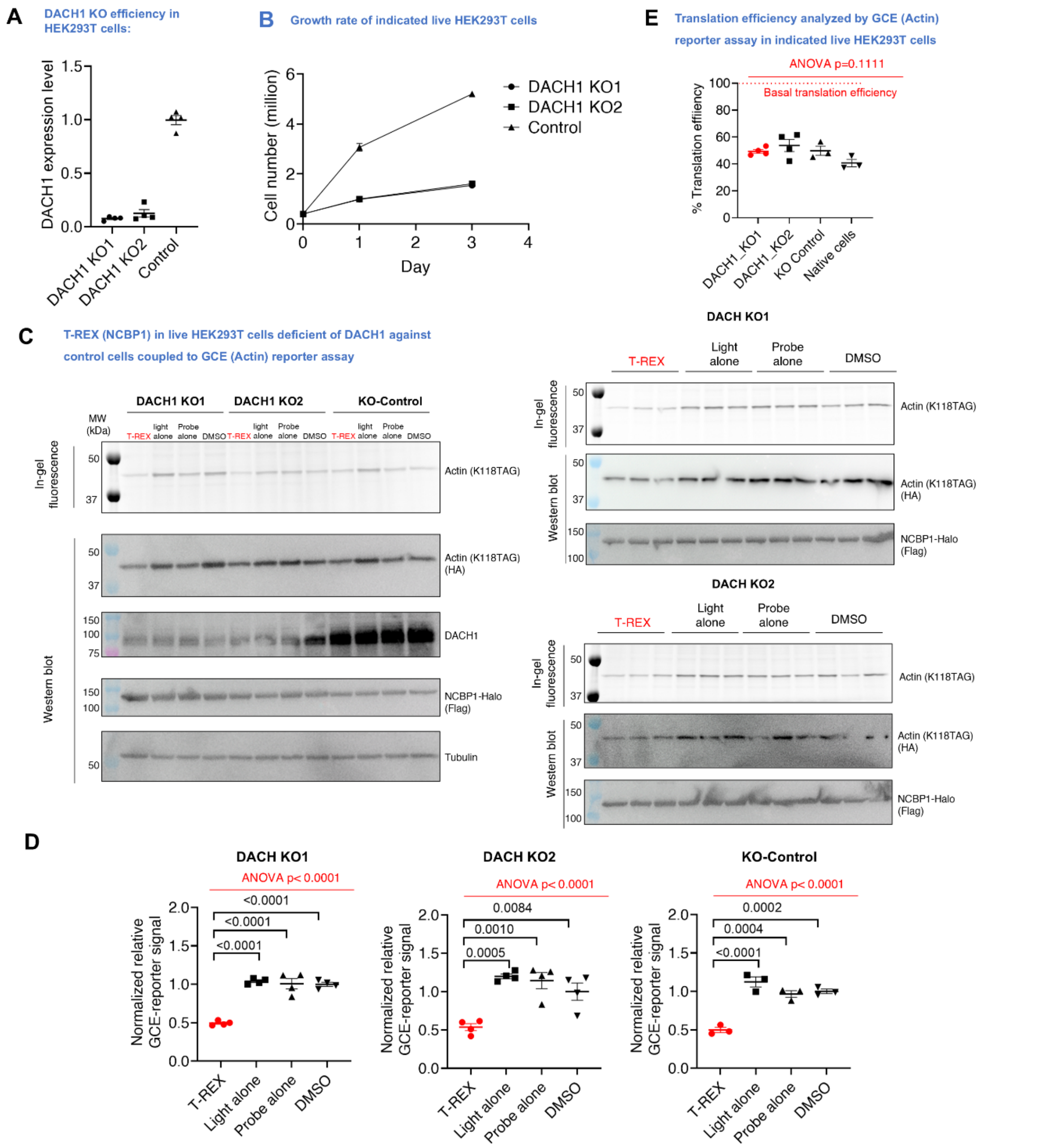


**Supplementary Fig. 25. HNEylation of NCBP1 downregulates protein translation independent of DACH1. A)** Two different DACH1 knockout (KO) and KO-control HEK293T lines were generated as described in **Methods**. KO efficiencies (92 ± 1% for KO line 1 and 88 ± 4% for KO line 2) were analyzed by western blot using anti-DACH1 antibody (showed in **B**). Tubulin served as loading control. n=4. **B)** HEK293T deficient of DACH1 grew slower compared to KO control. 0.4 Million HEK293T with DACH1 KO1, KO2 or control cells were seeded in individual 60 mm dish. The cells were harvested and counted using Countess II Cell Counter (ThermoFisher) at the indicated time. n=3. **C)** Representative gels and blots from T-REX–GCE integrated assay (see **Methods** and **Supplementary Fig. 1E**) conducted in DACH1 KO 1, 2 and KO-control lines**. D)** Data quantification is shown in **B**. *p* values were calculated using Tukey's multiple comparison test (n=4,4,3). **E)** Relative extent of translation efficiency under T-REX normalized to DMSO control. Data of native cells are derived from **Fig. 2F** (DMSO condition). 100% on the y-axis designates the basal translation efficiency. *p* values were calculated using Tukey's multiple comparison test. All data present mean±SEM. *n*=4,4,3,3 independent expreiments.

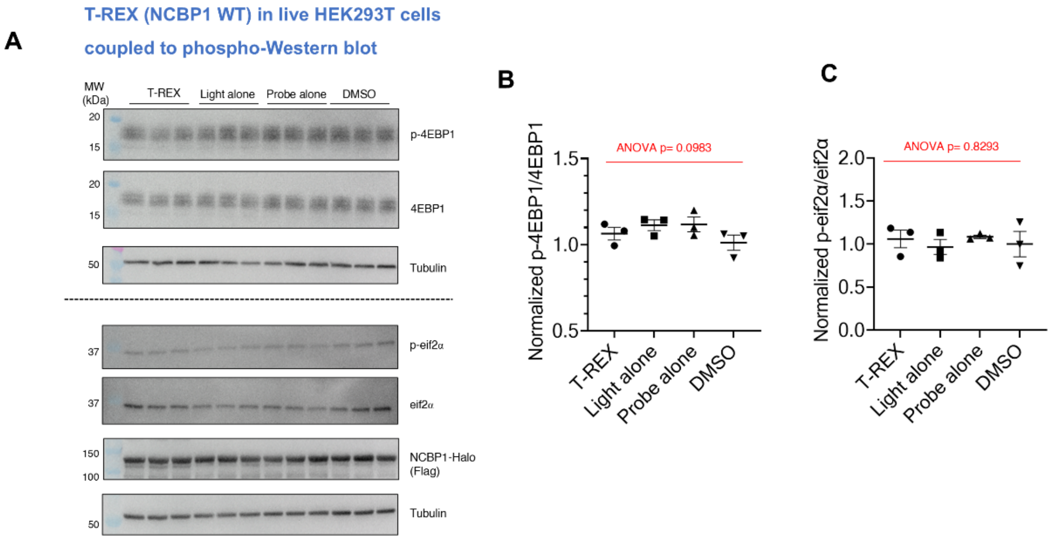


**Supplementary Fig. 26. NCBP1-HNEylation does not affect phosphorylation of endogenous 4EBP1 and eIF2α.** The expression levels of endogenous 4EBP1 and eIF2α, and those of the respective phosphorylated proteins, were analyzed 2 h post T-REX in HEK293T expressing NCBP1-Halo, against indicated controls. Post cell lysis, indicated antibodies were used to perform western blot analyses. See **Methods** for detailed lysis conditions to preserve phosphorylation states of proteins. **A)** Representative western blots: 4EBP1 (*top*); eIF2α (*bottom*). **B-C)** Quantification. *p* values were calculated using Tukey's multiple comparison test (*n*=3). All data present mean±SEM.

*Sequence alignment of S6K1-isoform 1 and isoform X*

| | | |
|---|---|---|
| isoform 1 | MRRRRRRDGFYPAPDFRDREAEDMAGVFDIDLDQPEDAGSEDELEEGGQ | 1-49 |
| isoform X | MRRRRRRDGFYPAPDFRDREAEDMAGVFDIDLDQPEDAGSEDELEEGGQ | 1-49 |
| isoform 1 | LNESMDHGGVGPYELGMEHCEKFEISETSVNRGPEKIRPECFELLRVLGK | 50-99 |
| isoform X | LNESMDHGGVGPYELGMEHCEKFEISETSVNRGPEKIRPECFELLRVLGK | 50-99 |
| isoform 1 | GGYGKVFQVRKVTGANTGKIFAMKVLKKAMIVRNAKDTAHTKAERNILEE | 100-149 |
| isoform X | GGYGKVFQVRKVTGANTGKIFAMKVLKKAMIVRNAKDTAHTKAERNILEE | 100-149 |
| isoform 1 | VKHPFIVDLIYAFQTGGKLYLILEYLSGGELFMQLEREGIFMEDTACFYLAE | 150-201 |
| isoform X | VKHPFIVDLIYAFQTGGKLYLILEYLSGGELFMQLEREGIFMEDTACFYLAE | 150-201 |
| isoform 1 | ISMALGHLHQKGIIYRDLKPENIMLNHQGHVKLTDFGLCKESIHDGTVTHT | 202-252 |
| isoform X | ISMALGHLHQKGIIYRDLKPENIMLNHQGHVKLTDFGLCKESIHDGTVTHT | 202-252 |
| isoform 1 | FCGTIEYMAPEILMRSGHNRAVDWWSLGALMYDMLTGAPPFTGENRKKTI | 253-302 |
| isoform X | FCGTIEYMAPEILMRSGHNRAVDWWSLGALMYDMLTGAPPFTGENRKKTI | 253-302 |
| | Coded by Exon 11 | |
| isoform 1 | DKILKCKLNLPPYLTQEARDLLKKLLKRNAASRLGAGPGDAGEVQAHPFF | 303-352 |
| isoform X | DKILKCKLNLPPYLTQEARDLLKKLLKRNAASRLGAGPGDAGEVQAHPFF | 303-352 |
| | Coded by Exon 12 / Coded by Exon 13 | |
| isoform 1 | RHINWEELLARKVEPPFKPLLQSEEDVSQFDSKFTRQTPVDSPDDSTLSE | 353-402 |
| isoform X | RHINWEELLARKVEPPFKPLLVHHPDLCLHVHMVFYLCAYLWPNFPFL | 353-400 |
| | Coded by Exon 14 | |
| isoform 1 | SANQVFLGFTYVAPSVLESVKEKFSFEPKIRSPRRFIGSPRTPVSPVKFSP | 403-453 |
| | Coded by Exon 15 | |
| isoform 1 | GDFWGRGASASTANPQTPVEYPMETSGIEQMDVTMSGEASAPLPIRQPN | 454-502 |
| isoform 1 | SGPYKKQAFPMISKRPEHLRMNL | 503-525 |

**Supplementary Fig. 27. Alignment of amino acid sequences of S6K1 full length (FL) and S6K1-isoform X** Sequences coded by exons 11, 12, 13, 14, 15 are indicated using lines of different color. The sequence at the C-terminus of isoform X, which is different from isoform 1 due to alternative splicing, is highlighted in red. **Note:** Kinase domain and ATP-binding site within S6K1-**FL** remained intact in S6K1-**X**, and so did the mTORC1 stimulatory phosphorylation site (T389, highlighted in yellow).

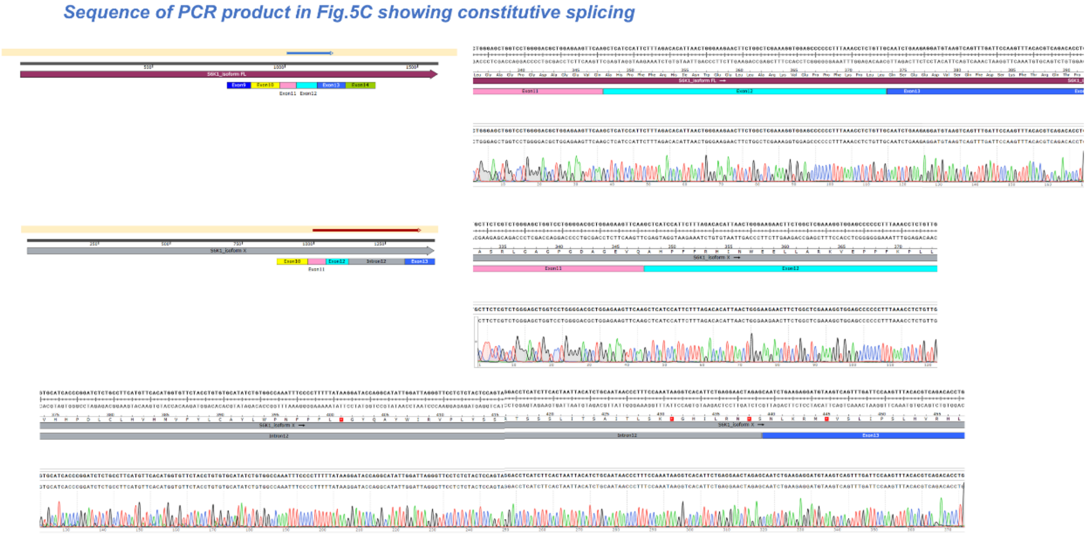


**Supplementary Fig. 28. (Note: raw sequencing output data in this figure are split over two pages for clarity). Relating to datasets in Fig. 5C-D, sequencing validation of S6K1 full length (FL) and isoform X.** The requisite amplicons (the lower band in the gel in **Fig. 5C from S6K1 FL** and the upper band in the gel in **Fig. 5C from S6K1 isoform X**) were excised and gel-extracted (A9285, Promega), and sequenced using the corresponding forward primer used in the experiment that gave rise to the dataset shown in **Fig. 5C-D**. The sequencing results covering regions E11—E12—E13 are shown.

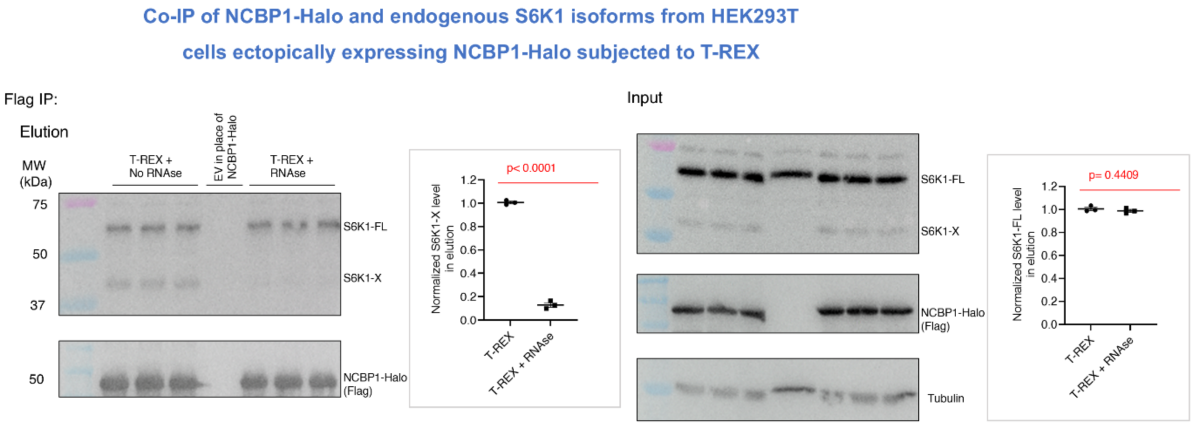


**Supplementary Fig. 29. S6K1-X–NCBP1 interaction, but not S6K1–NCBP1 interaction, is RNA dependent.** The live-cell-based T-REX experiments were performed similarly as described in **Fig. 5E**, except that post lysis, samples were treated with either RNase A (100 ug/mL) or buffer alone and incubated (30 min, 37°C), prior to further incubation with Flag beads. *Insets* show quantification *p* values were calculated with an unpaired, two-tailed t-test (*n*=3). All data present mean ±SEM.

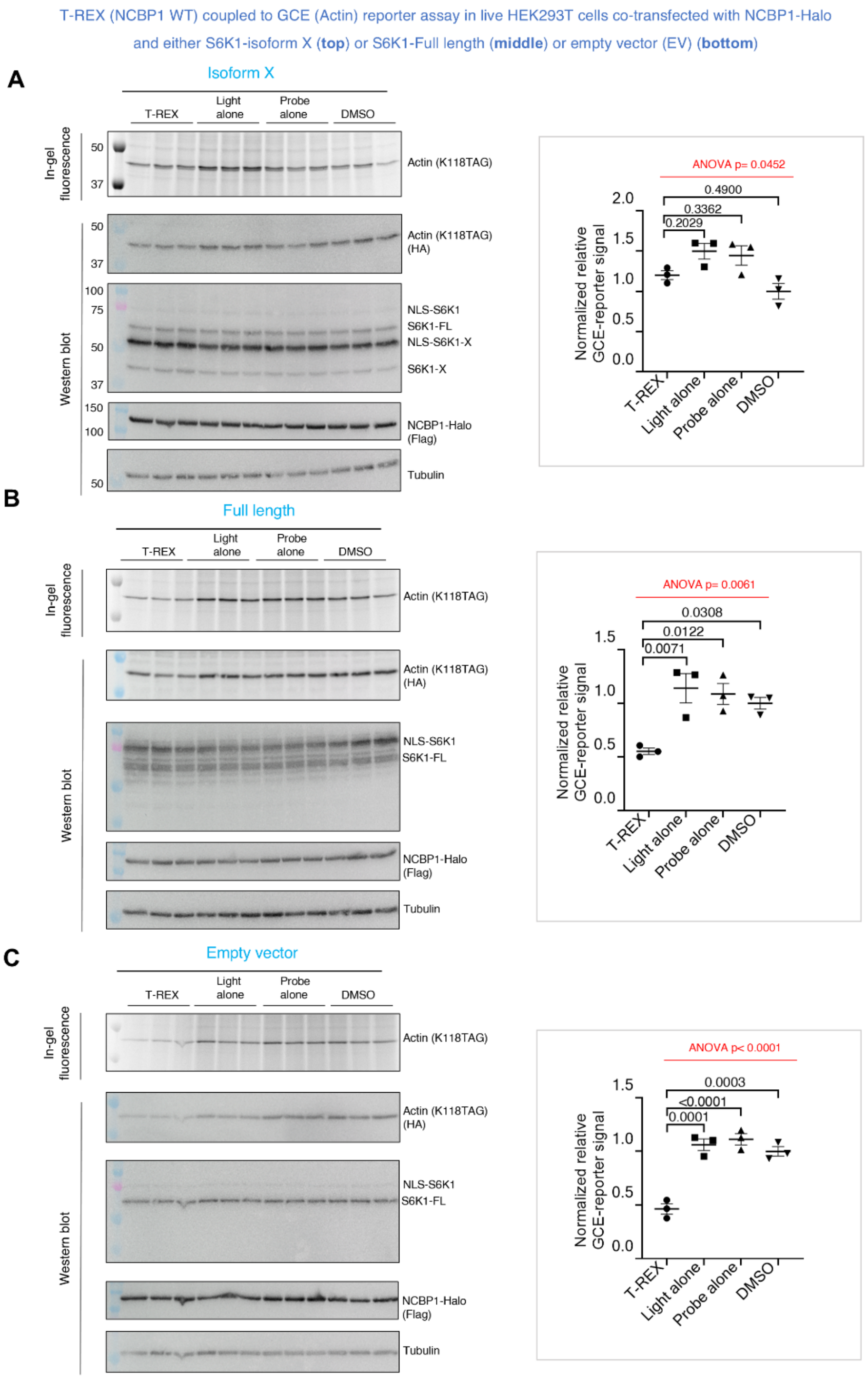


**Supplementary Fig. 30. Effects of overexpression of full-length S6K1 and truncated isoform S6K1-X on NCBP1-HNEylation-induced suppression of protein translation**. HEK293T were co-transfected with plasmids encoding either **(A)** S6K1-isoform X, **(B)** S6K1-full length (FL), or **(C)** empty vector (EV), and GCE-reporter constructs as described in **Methods** and **Supplementary Fig. 1E** for T-REX—GCE integrated workflow. The ratio of plasmids of S6K1-FL or S6K1-X (or EV): NCBP1-Halo : Actin (K118TAG) : tRNA synthetase is 1:1:1:0.2. The cells were subjected to T-REX against indicated controls, followed by treatment with BCNK (50 µM, 4 h). Post cell lysis, and coupling with tertrazine-Cy5, the samples were analyzed by in-gel fluorescence and western blot analyses using indicated antibodies. *Insets* in each case show respective quantification. p values were calculated using Tukey's multiple comparison test (*n*=3). All data present mean±SEM.

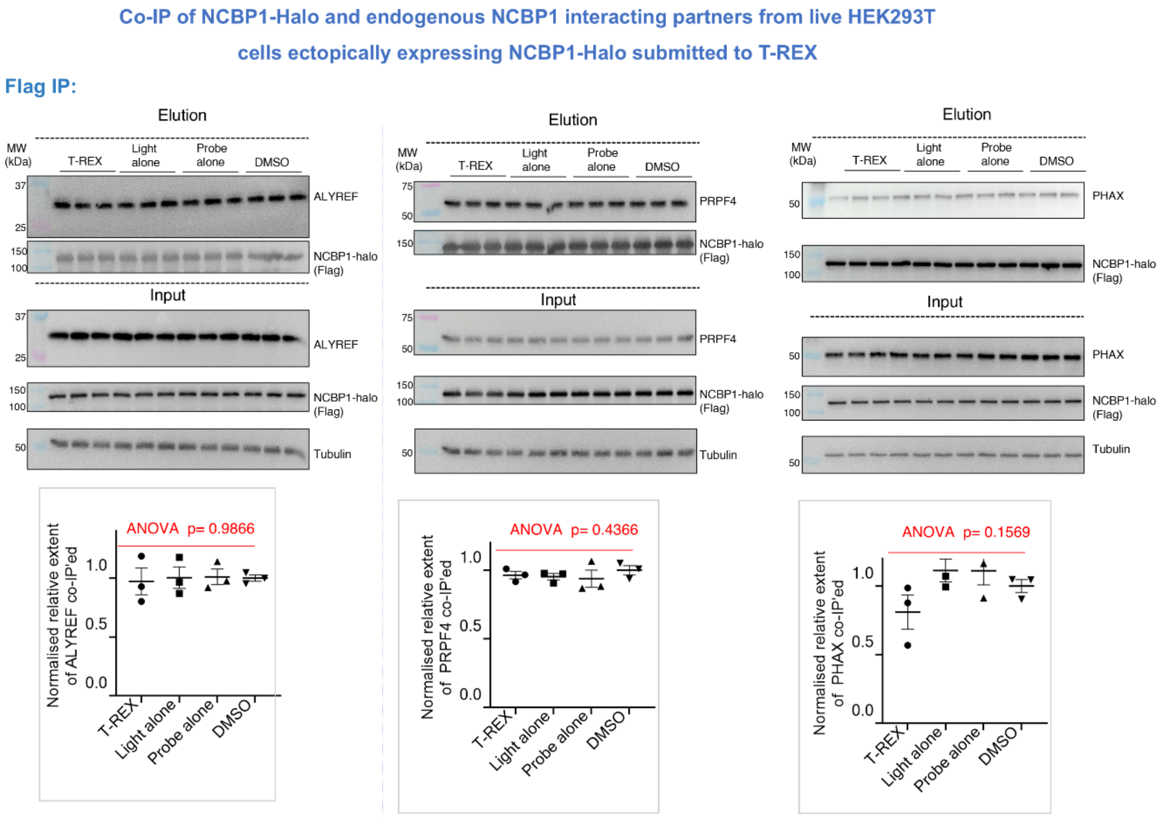


**Supplementary Fig. 31. Relating to Fig. 6B, investigating the possible HNE-dependent effects on association between NCBP1 and either ALYREF, PRPF4, or PHAX shows no statistically significant change in association upon NCBP1-HNEylation.** The experiment was performed similarly to **Fig. 6B**. Following T-REX-enabled NCBP1-HNEylation in HEK293T (see **Methods** for details) expressing NCBP1-Flag-Halo, against indicated T-REX technical controls, Anti-Flag pulldown (see **Methods** for details) was performed. Input and elution samples were subjected to SDS-PAGE, and western blot analyses using indicated antibodies. *Right panel* quantification of the relative extent of ALYREF, PRPF4, or PHAX co-eluted with NCBP1-Flag-Halo in the absence and presence of NCBP1-HNEylation. Data were normalized by NCBP1-Halo expression. *p* values were calculated using ANOVA and Tukey's multiple comparison test (*n*=3). All data present mean±SEM.

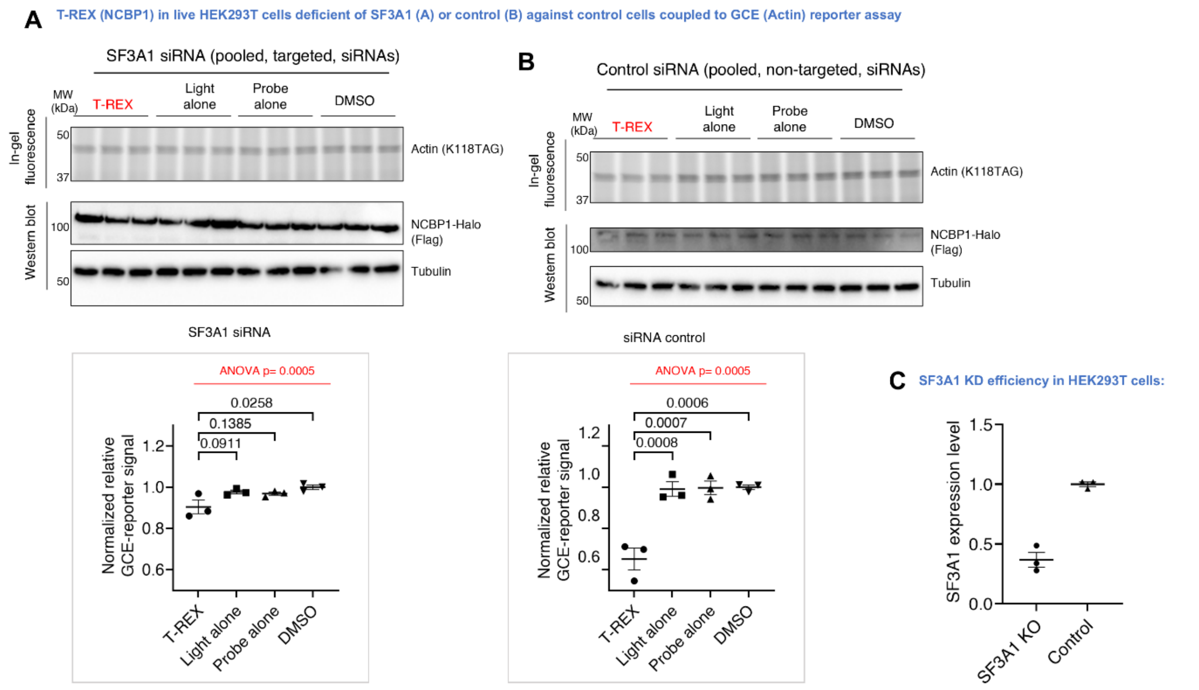


**Supplementary Fig. 32. SF3A1 depletion ablated translation inhibition driven by NCBP1-HNEylation.** HEK293T co-transfected with NCBP1-Halo and GCE(actin) reporter constructs were subjected to either SF3A1 siRNA (**A**), or non-targeted control siRNA (**B**), using DharmaFECT Duo Transfection Reagent following manufacturer's protocol. T-REX-GCE assay was performed as described in **Methods**. Data quantification is shown in *Right panel*. *p* values were calculated using Tukey's multiple comparison test ($n$=3). All data present mean±SEM. **C)** KO efficiency of SF3A1 (63 ± 7%) analyzed by western blot using anti-SF3A1 antibody (also see **Fig. 6C**).

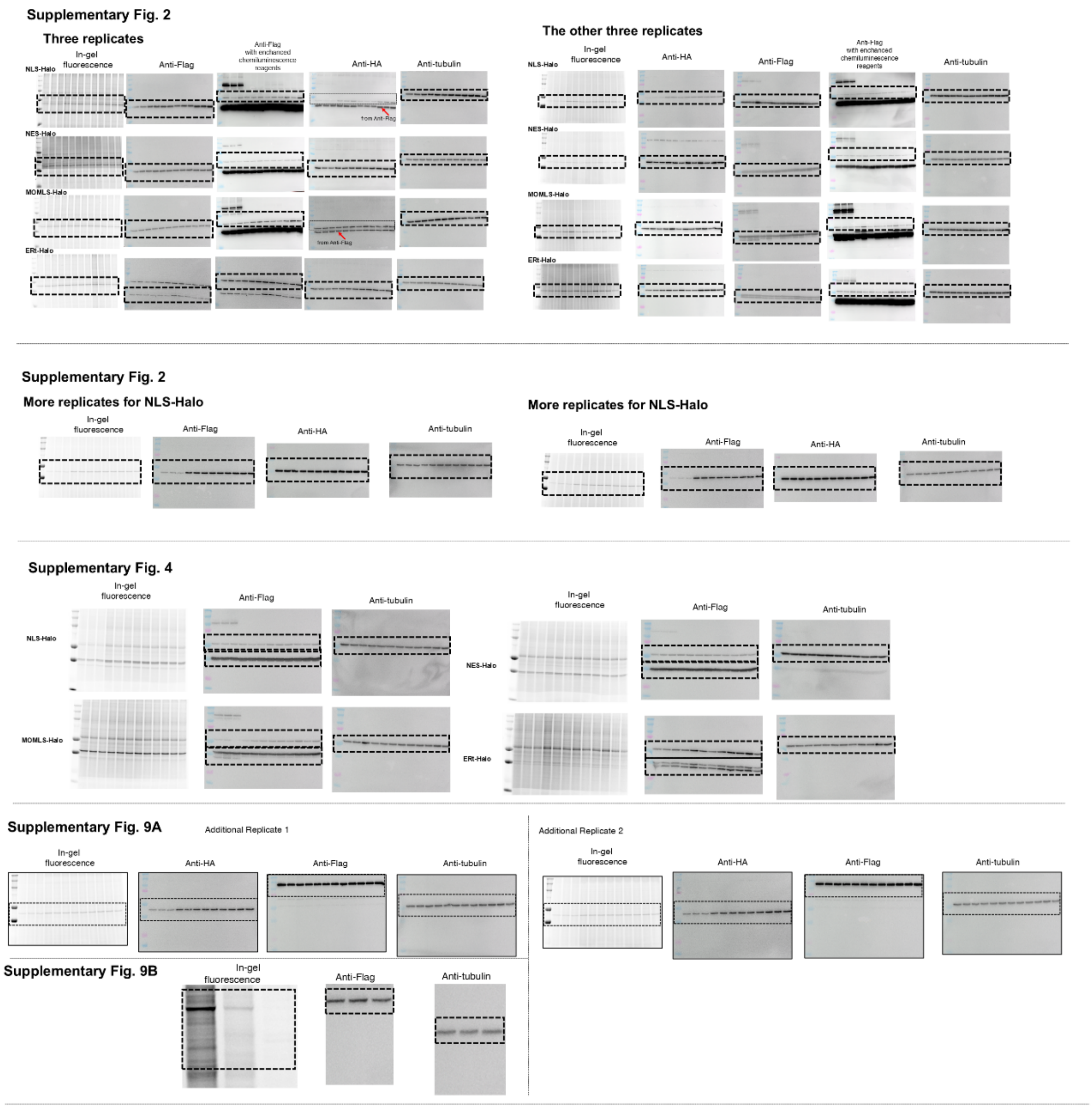


**Supplementary Fig. 33. Unprocessed western Blots for Supplementary Figures.**

**Supplementary Fig. 11A**

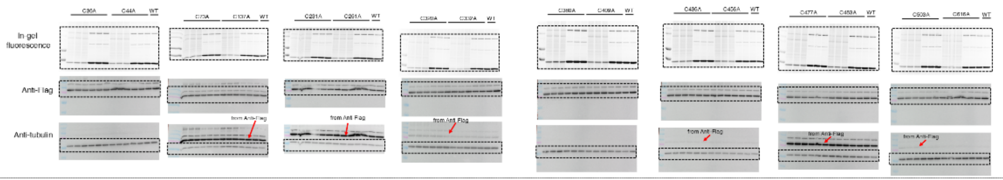


**Supplementary Fig. 11A**

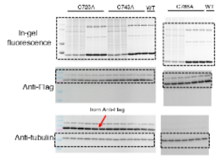


**Supplementary Fig. 11B**

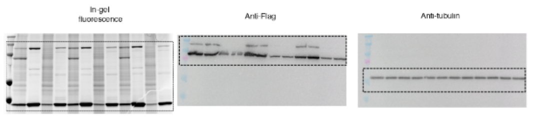


**Supplementary Fig. 12**

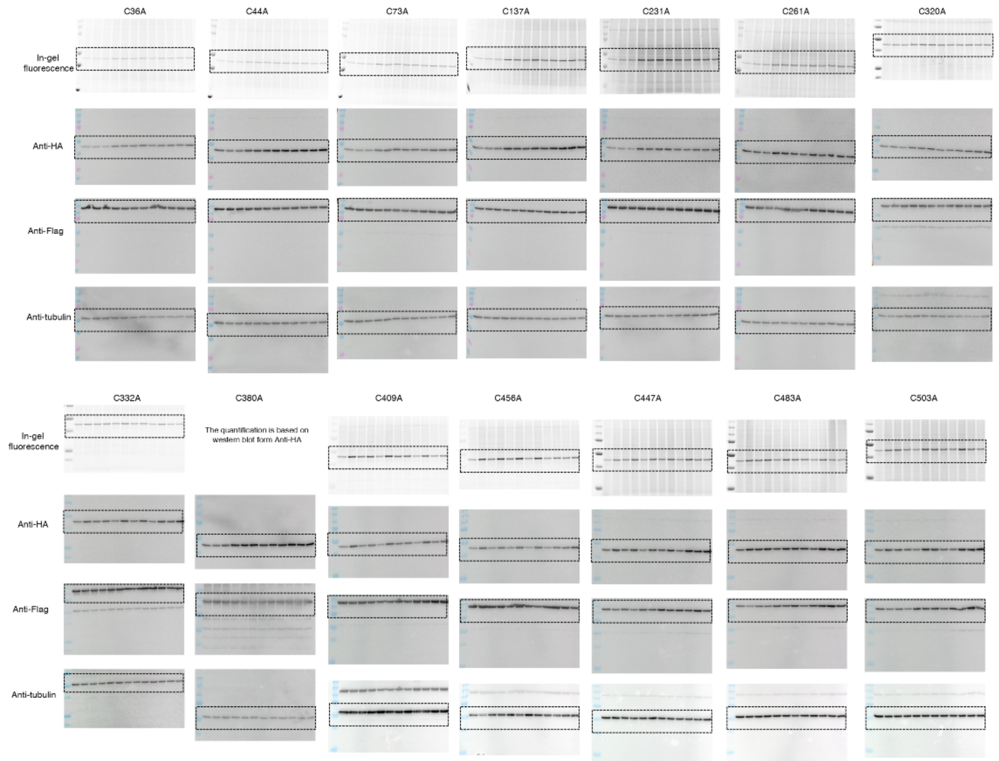


**Supplementary Fig. 34. Unprocessed western Blots for Supplementary Figures.**

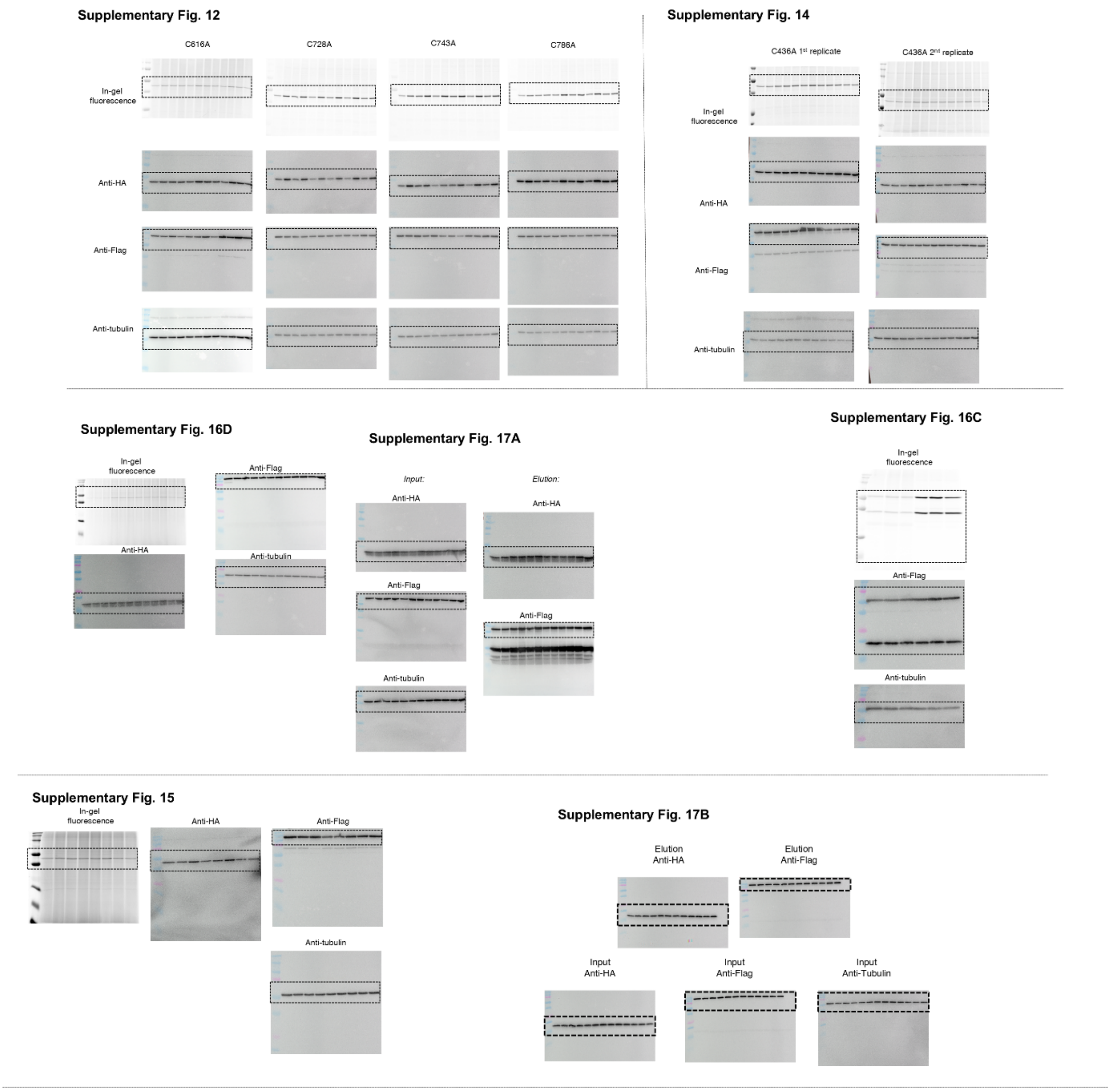


**Supplementary Fig. 35. Unprocessed western Blots for Supplementary Figures.**

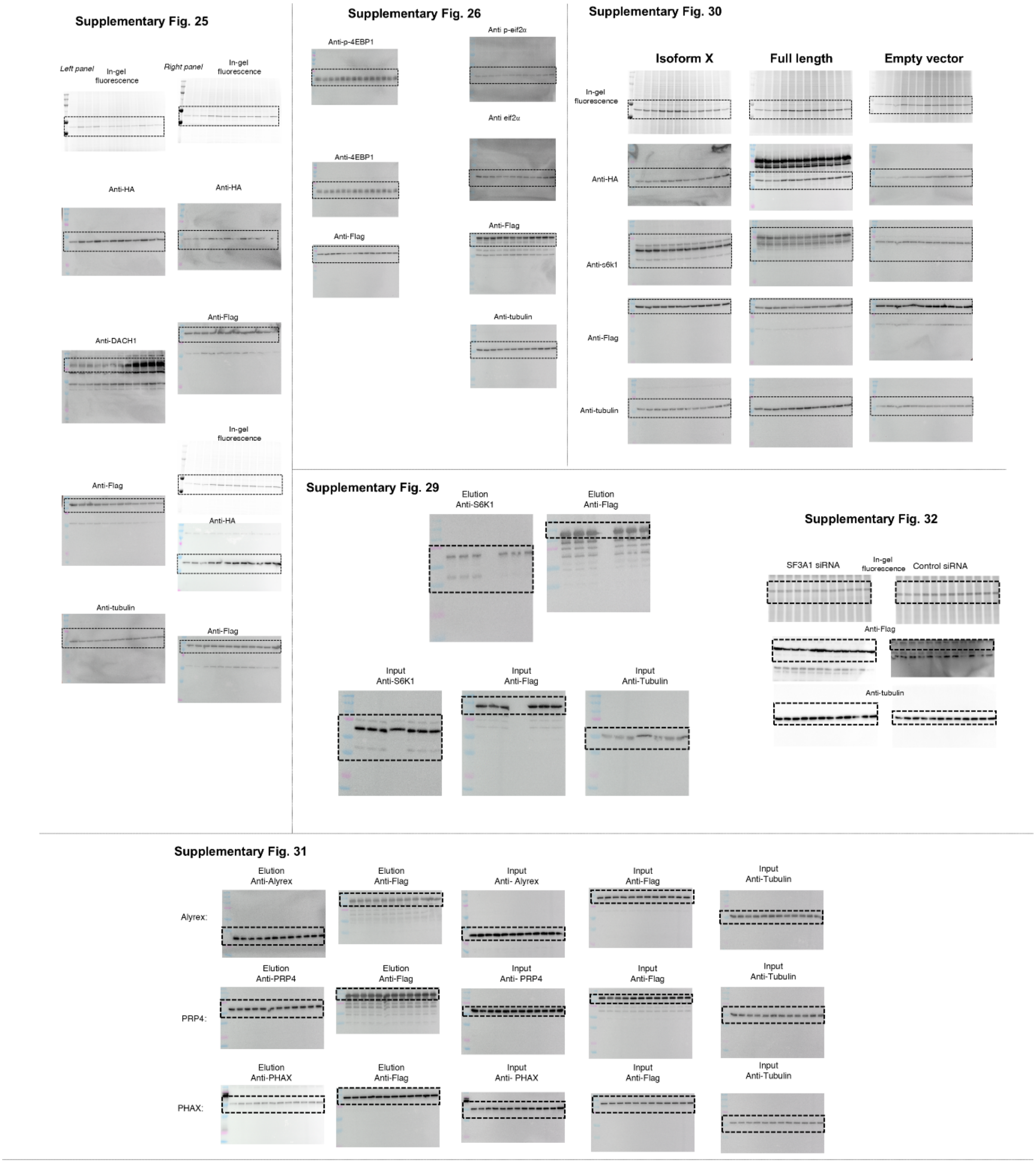


**Supplementary Fig. 36. Unprocessed western Blots for Supplementary Figures.**
